# Sensitivity and robustness of comorbidity network analysis

**DOI:** 10.1101/752964

**Authors:** Jason Cory Brunson, Thomas P. Agresta, Reinhard C. Laubenbacher

## Abstract

1

**Background:** Comorbidity network analysis (CNA) is an increasingly popular approach in systems medicine, in which mathematical graphs encode epidemiological correlations (links) between diseases (nodes) inferred from their occurrence in an underlying patient population. A variety of methods have been used to infer properties of the constituent diseases or underlying populations from the network structure, but few have been validated or reproduced.

**Objectives:** To test the robustness and sensitivity of several common CNA techniques to the source of population health data and the method of link determination.

**Methods:** We obtained six sources of aggregated disease co-occurrence data, coded using varied ontologies, most of which were provided by the authors of CNAs. We constructed families of comorbidity networks from these data sets, in which links were determined using a range of statistical thresholds and measures of association. We calculated degree distributions, single-value statistics, and centrality rankings for these networks and evaluated their sensitivity to the source of data and link determination parameters. From two open-access sources of patient-level data, we constructed comorbidity networks using several multivariate models in addition to comparable pairwise models and evaluated differences between correlation estimates and network structure.

**Results:** Global network statistics vary widely depending on the underlying population. Much of this variation is due to network density, which for our six data sets ranged over three orders of magnitude. The statistical threshold for link determination also had strong effects on global statistics, though at any fixed threshold the same patterns distinguished our six populations. The association measure used to quantify comorbid relations had smaller but discernible effects on global structure. Co-occurrence rates estimated using multivariate models were increasingly negative-shifted as models accounted for more effects. However, only associations between the most prevalent disorders were consistent from model to model. Centrality rankings were likewise similar when based on the same dataset using different constructions; but they were difficult to compare, and very different when comparable, between data sets, especially those using different ontologies. The most central disease codes were particular to the underlying populations and were often broad categories, injuries, or non-specific symptoms.

**Conclusions:** CNAs can improve robustness and comparability by accounting for known limitations. In particular, we urge comorbidity network analysts (a) to include, where permissible, disaggregated disease occurrence data to allow more targeted reproduction and comparison of results; (b) to report differences in results obtained using different association measures, including both one of relative risk and one of correlation; (c) when identifying centrally located disorders, to carefully decide the most suitable ontology for this purpose; and, (d) when relevant to the interpretation of results, to compare them to those obtained using a multivariate model.

## 2 Introduction

### 2.1 Background

The importance of comorbidity to medical research and practice is fundamental. Usually, the term refers to morbidity that complicates an index condition for an individual patient, which produces population-level associations in epidemiological studies and clinically relevant differences in prognosis or response to treatment and which analyses must take into account (Akker et al. 1996; Valderas et al. 2009). Recently, several research teams have taken a systems approach to population comorbidity by analyzing “comorbidity networks” (also “disease networks”, “disease graphs”, and maps of “disease space” or of the “diseasome”) aggregated from measures of co-occurrence between pairs or among small groups of disorders (Emmert-Streib et al. 2013; Capobianco and Liò 2015). Their investigations seek to uncover novel clinical associations, to stratify patient populations, and to predict disease progression, among other aims (Brunson and Laubenbacher 2017).

In this section we review the recent comorbidity network analysis literature and articulate some concerns with the methods used. In subsequent sections, we test how sensitive the network statistics produced, and by extension the kinds of inferences drawn, in these studies are to the source of data and the method of network construction. We conclude with a set of recommendations that we think will strengthen the practical value of future comorbidity network analyses.

### 2.2 Conventional comorbidity network analysis

*Systems medicine* consists in the adoption into medical research of principles and techniques from systems biology, which in turn are described as global, integrative, and holistic (Hood et al. 2013; Ayers and Day 2015; Kirschner 2016). Networks are a prime illustration, having become a staple of systems biology (Pavlopoulos et al. 2011) and seen extensive use in systems medicine (Gietzelt et al. 2016). Since 2009, alongside an expansion of the scope and scale of social network analysis in medicine, network analysis has accelerated the systems approach to human health (Brunson and Laubenbacher 2017).

Any complex system can be conceptualized as a network, and a range of structures are called network models, but *network science* rests predominantly on the theory of mathematical graphs, and therefore on the discretization of underlying relations. Converting subject-level expression or incidence data to (often unweighted) network models results in the loss of covariance, sign, and magnitude information. This conversion makes the analysis of large data sets more tractable, as network models occupy far less memory than incidence data and can be analyzed and simulated using highly efficient algorithms. In most applications, however, network analysis relies on theoretical assumptions that do not necessarily hold for association data (Brandes et al. 2013). As a consequence, systems analyses of high-dimensional data may be presented as network analyses despite making little or no use of graph theory, an observation made by Williams et al. (2014) of systems biology and later by Brunson and Laubenbacher (2017) of systems medicine. For example, comorbidity network studies are often strictly dyadic—focused on pairwise relations but not on dependencies among these relations—as in the search for new or poorly understood associations between clinical concepts such as diagnosed disorders (e.g. Hanauer et al. 2009; Hanauer and Ramakrishnan 2013). Meanwhile, classical techniques such as hierarchical clustering (e.g. Roque et al. 2011) and ordination (e.g. Lyalina et al. 2013) preserve more information from association data than graph-tailored alternatives like community detection and force-directed layouts.

There is also a conceptual mismatch between the tools designed to investigate actor-oriented networks and the questions motivating variable-oriented association studies. Much of contemporary network science appropriates graph-theoretic operationalizations of such sociological concepts as community, centrality, and brokerage, which have natural and important applications in population health, health economics, and other socio-structural research using relational data. However, these concepts have also been applied to network models of association data, without the motivations and interpretations that would attend a theoretical foundation independent of these social scientific roots, and health informatics is no exception. This begins with the very concept of a relation, or link, which is usually defined for pairs of clinical concepts by imposing an evidential or evaluative cutoff on a measure of association between binary variables, e.g. rejecting a null hypothesis of no association at *p* < .05 based on a chi-squared test (Davis and Chawla 2011) or having an odds ratio greater than 3 (Hanauer et al. 2009). The observations of Hidalgo et al. (2009), for example—that the lethality of a disease correlates with its connectivity and that health states tend to progress from less to more connected diseases—may depend on disease prevalences and other features of the population (c.f. Chmiel et al. 2014) and in part be artifacts of sample size (c.f. Blair et al. 2013) or of patient-level covariates (c.f. Rzhetsky et al. 2007).

We do not suggest that applications of graph theory to population comorbidity data are inherently problematic. The aforementioned observations of Hidalgo et al. (2009) have been reproduced (Glicksberg et al. 2016), the strengths of association between disorders are consistent across a range of data sets (Blair et al. 2013), and several studies using graph-theoretic methods have yielded interesting results. For example, Chen and Xu (2014) used a random walk–based notion of distance to rank the comorbidities of different cancers, in an effort to identify targets for fruitful follow-up laboratory research (Chen et al. 2015). Other studies have illustrated ways in which network analysis complements geometric techniques: Lyalina et al. (2013) visualized co-occurrence data using both principal component analysis biplots and co-occurrence network layouts, from which they gained complementary insights to the problem of differentiating between distinct disorders with shared symptoms. Nevertheless, the irrigidity with which links—the observational units of network analysis—are determined and interpreted in this setting call for additional caution.

### 2.3 Objectives

We articulate four concerns with the conventional approach:^1^

1. Sources of disease incidence data differ in their conventions, completeness, and representativeness, and the consequent differences in comorbidity network structure have not been explored.
2. The method of link determination, meaning the determination of whether to link two disorder codes in the network model based on a statistical signal or strength measure on the cooccurrence data, varies across studies. What effect does imposing an error rate correction on the p-values obtained from pairwise *χ*^2^ tests, or using a binary correlation coefficient to measure the strength of association instead of the odds ratio, have on the properties of the resulting network?
3. A network model aggregated from pairwise correlations discards information about interactions among larger subsets of variables, though these potential effects are important both to the network perspective and to the systems paradigm.
4. The use of network statistics like connectivity, distance, and centrality to characterize networks of disorders depends on correspondences between the definitions and values of these statistics and the theoretical constructs they are being used to measure. Little theoretical guidance is available for the study of networks representing statistical associations rather than directly observed relations.

We address concerns 1 and 2 in a sensitivity analysis. In the following sections, we describe an analysis pipeline to characterize a pairwise co-occurrence dataset using several single-valued network summary statistics and node centrality. We evaluate the sensitivity of the results to the source of data, the choice of association measure, and the strength of evidence and of association used to determine links. We address concern 3 in a comparison of a typical pairwise construction to three alternative constructions that control for progressively more factors when estimating co-occurrence rates. For this analysis we focus on the effects of the modeling choice on these estimates. Finally, we touch upon concern 4 in the Discussion section after reviewing the results of these analyses. Our goal in the present study is to better understand any systematic biases built in to the comorbidity network approach and how they might be addressed.

## 3 Summary of Analyses

We conducted robustness and sensitivity analyses of several network-analytic results of the kind that have been reported in the CNA literature. See the full Methods and Results sections below for detailed discussions and citations.

Six of our data sets, all used for pairwise analysis, were provided by the authors of previous CNAs; one, the intensive care unit database MIMIC-III, is freely available for research use; and one, results of the 2011 National Ambulatory Medical Care Survey (NAMCS), was obtained from the website of the Centers for Disease Control and Prevention. Our pairwise study examined the effects of the data source; the test-wise error rate (TWER) at which links were determined (evidential threshold); how, if at all, TWERs were corrected for multiple comparisons; the binary association measure (BAM) used to weight links; and the value of the BAM, if any, at which to prune links (evaluative threshold). A second study compared the pairwise approach to multivariate network constructions on the available incidence-level data sets. These comparisons held other parameters fixed and examined correlation structures directly as well as their network models.

Comorbidity networks have been described as “scale-free” based on the power-law appearance of their degree distributions. However, rigorous tests of this hypothesis, based on competing interpretations of power-law fitting, agree that comorbidity network degree distributions do *not* follow power laws. This held true across the range of pairwise constructions. (The degree distributions tended to more closely follow log-normal distributions.)

CNAs often summarize networks in terms of global statistics such as degree assortativity, clustering coefficient, and modularity, but the choices of summary statistics are evidently arbitrary. Regressions of several global network statistics on the construction parameters revealed that different data sources yield highly distinct networks, though specific cases suggest that pre-processing choices can be equally important. Most global statistics change as expected with graph density, whether due to stricter evidential or evaluative cutoffs; the exception was the clustering coefficient, which may depend on the interaction between these cutoffs due to indirect comorbidities such as common risk factors. A principal components analysis demonstrated that variability was explained primarily by graph density, followed by data source, and that networks from different sources were better distinguished under stricter cutoffs.

Studies often identify the disorders that occupy highly central positions (“hubs”) in comorbidity networks, with the expectation that they are likely to be more fruitful targets of epidemiological, biological, or clinical follow-up. We quantitatively compared full rankings of three common centrality measures between networks constructed from data sets using the same ontology, and we described and compared the several most central disorders within and between ontologies. Full rankings were highly sensitive to the evidential cutoff (TWERs) and/or the BAM, and which parameter was more determinative varied by data source. Networks constructed from regional EHR data had assorted and often non-specific symptoms and disorders at their centers, including epilepsy, limb pain, respiratory problems, vitamin deficiency, benign neoplasms, and tuberculosis. Other data sources produced their own distinctive hubs. These were usually explained by sheer numbers of epidemiological comorbidities, in part due to high prevalence consistent with their specific subpopulations. In both settings the same disorders usually topped the rankings by different centrality measures.

Though comorbidity networks are aggregated from pairwise associations in incidence data, rigorous tests of epidemiological comorbidity account for patient-level covariates as well as associations with other population-level disorders. We compared pairwise constructions to two multivariate constructions using partial correlations and joint distribution models (JDMs) on the NAMCS and MIMIC-III data, separated into subpopulations by care unit. As expected, comorbidities between less prevalent disorders were often indiscernible in the multivariate models, which controlled for the effects of disorders outside each pair. Compared to pairwise correlations, which in all networks were frequently strong and overwhelmingly positive, partial and JDM correlations were on average negative-shifted, including many more negative associations. Consequently, the resulting network models were much sparser and included many more negative links. At least for highly prevalent disorders, JDM correlation estimates were better predicted by pairwise correlations than by partial correlations. A centrality analysis yielded similar results to that of the pairwise constructions: Hubs were robust to the choice of model, while overall centrality rankings were only weakly concordant and no group two models produced similar rankings across all units.

## 4 Discussion

We summarize the results of our analyses in Section 6, with references to supporting figures and tables. In this section, we discuss the implications of these results for the practice of comorbidity network analysis and recommend an analytic workflow informed by these implications. CNA ranges widely in motivations and methodologies, but we believe that a study that follows our recommendations will arrive at more robust and interpretable results.

### 4.1 Sensitivity of pairwise network structure

We explored the dependence of certain network properties on certain parameters of comorbidity network construction. These parameters include the data source (usually an EHR or claims database), the evidential threshold (p-value) used to discern links, the method of correcting for multiple comparisons (if any), and the binary association measure (BAM) used to prune weak links and/or to weight remaining links. The properties considered are network density; shape of the degree sequence; several single-valued global statistics having to do with the degree sequence, local link dependencies, and geodesic distances; and several measures of node centrality. These are a fraction of the possible constructions and properties and may represent only a minority of those that have so far been published. Nevertheless, our observations allow us to draw several provisional but clear conclusions about the methodological robustness of CNA.

One observation, about the shapes of their degree sequences or distributions, is unequivocal. While the proper way to define and detect power laws from empirical data is contested (Broido and Clauset 2018; Voitalov et al. 2018), proposed standards from both camps (Gillespie 2015; Voitalov 2018) lead to the same conclusion, that the degree sequences of comorbidity networks do *not* follow power laws. This runs counter to a frequent claim within the literature. While most CNA researchers have inferred from supposed scale-freeness only the existence of highly-connected “hubs” (Steinhaeuser and Chawla 2009; Ball and Botsis 2011; Divo et al. 2018), the assumption underpins some more advanced analytic techniques used to prune weak associations prior to analysis (Jiang et al. 2018) or to model network growth based on the accumulation of database records (Scott et al. 2014). Such techniques may yet be useful, but the weaknesses at their foundations should not be overlooked.

A second observation concerns the structure of comorbidity networks more generally. While the network architecture is sensitive to every parameter of construction, the underlying population is far more determinative than the choices of link determination or of weighting scheme. This holds true for all of the properties we evaluated. This is a favorable result for the genre, since the goal of many recent comorbidity network applications has been to assess differences in the structures of comorbidity networks derived from different populations (Warner et al. 2015; Feldman et al. 2016; Glicksberg et al. 2016). For example, Divo et al. (2018) introduced a case–control study design to CNA to compare network hubs between COPD and non-COPD patient populations, in order to better identify potential targets for follow-up research or clinical intervention specific to COPD. While our secondary use of processed data prevents us from separating the effects of population, practice, and pre-processing, our analysis supports the premiss that it can be meaningful to identify comorbidity network properties that are characteristic of certain populations.

While comorbidity networks constructed from different data sets are clearly discriminable by their global properties, these differences are largely orthogonal to those due to changes in network density: decreased density (increased sparsity), due to tightening of evidential or evaluative cutoffs, results in more skewed degree distributions and less modular structure. Meanwhile the underlying populations are better distinguished by rates of overall connectedness, assortative linking, and triad closure. To characterize comorbidity networks from a moderate number of populations, then, it may be more useful to report such statistics as connected component size distribution, assortativity, and clustering coefficient than the average geodesic distance, Gini index, or modularity.

When comorbidity networks are weighted, on the other hand, the choice of weight may strongly impact the results. This is based on three observations: First, the shape of the weight distribution differs substantially between relative risk–like BAMs (the Forbes coefficient and the odds ratio) and correlation coefficients (the Pearson binary and the tetrachoric). Second, BAM cutoffs appear to be more important determinants than p-value cutoffs. Third, node centrality can depend dramatically and idiosyncratically on which BAM is used to measure and prune network links. Sensitivity to the choice of weights has long been observed in the literature (Hidalgo et al. 2009), but the motivation for selecting one measure over others for CNA often goes unstated. Comparisons are only possible between some published comorbidity network studies because certain exploratory conventions, in particular the combined use of *F* and *ϕ*, have been adopted by later investigators (see Section 5.1.2).

Finally, as mentioned above, node centralities are quite robust to the choice of evidential cutoff, but less robust to the choice of BAM used to impose a evaluative cutoff. This is especially true for “hubs”, the nodes in each network with the highest centrality. Different disorders may be identified as hubs by different centrality measures, but the same small subset of hubs tends to appear in networks constructed from any specific population.

### 4.2 Multivariate network models

We compared two multivariate network modeling approaches, partial correlation networks and joint distribution networks, to the conventional pairwise correlation network approach. In this comparison, all models were based on latent, normally-distributed risk factors and produced estimates of the correlations among them. Both multivariate approaches had the intended and expected effects of reducing both the estimated values of and the nominal statistical evidence for most pairwise associations, which are interpreted as comorbidities. Models deeper into the systems paradigm represented comorbidities as being less aligned with each other or with a single dimension of poor health. As a result, more negative associations, particularly involving codes for depression and cancer versus those for metabolic and cardiac disorders, emerged. Also as expected, associations between more prevalent disorders were more robust to the choice of model and would therefore lead to more consistent interpretations.

In addition to having fewer links, the multivariate models estimated consistently more negative associations than the pairwise, at least among more prevalent disorders. Indeed, the overall reduction in association estimates included some positive pairwise associations shaking out as negative in the multivariate models. In this way the multivariate models appear to be preventing errors both of magnitude and of sign (Gelman and Carlin 2014). Based on these patterns, we expect in general that models that account for more sources of variation will discern strong evidence for fewer epidemiological comorbidities but for more population-level disassociations.

The partial correlation and joint distribution models yielded radically different alternatives to the pairwise model, the former large and sparse, the latter small and dense. The pairwise correlation estimates among more prevalent disorders were roughly linearly related both to the partial correlation estimates and to the joint distribution estimates. However, this relationship broke down among the less prevalent disorders. Taking the joint distribution models to provide more realistic summaries, this suggests that estimating pairwise disease associations from a larger diseaseome introduces a uniform positive bias in the measured associations, which increases the apparent strengths of the comorbid relations and mistakes some null relationships, and possibly some relationships of mutual exclusion, as comorbidities. Moreover, introducing exogenous covariates into a multivariate model yielded bimodally-distributed effect estimates, which indicate that much of the perceived interaction among disorders, in addition to their prevalence, can be accounted for by demography and environment.

The linearity of some of these relationships stands in stark contrast to the non-linear relationships observed between estimates obtained using joint distribution models and classical co-occurrence indices not based on the same latent structure (Pollock et al. 2014). This reaffirms the importance of comparing like with like, insofar as this is possible, even amidst fundamentally different modeling approaches.

We emphasize that these multivariate models no more reveal “true” relationships than pairwise models. Indeed, the inclusion or exclusion of certain patient-level predictors or other diseases has the potential to again upend the results. Rather, as the authors of several comorbidity networks have stressed, these “phenome-wide” analyses must be understood as exploratory; the interplay between specific diseases is best understood through more narrowly-targeted research. Certainly far more clinically relevant epidemiological comorbidities exist in critical care populations than were captured by the partial correlation and joint distribution networks. Nevertheless, these networks serve as valuable checks on conventional constructions, which are certain to contain many indirect and spurious associations.

### 4.3 Interpretability of network statistics

Though we focus on the robustness of the calculations—the reliability of the numerical results— equally important is the soundness of the interpretations—their validity. Networks are an increasingly popular analysis tool for high-dimensional data sets. Though the ways in which their properties are interpreted draws heavily from analyses of social networks and networks based on other relational data (e.g. electrical signals, protein interactions), their construction is fundamentally different. The conclusions we draw about complex systems from network models of their constituent interactions must be informed by the process that converts the raw data to the network model.

The use of centrality measures is a case in point. The degree of a disorder, calculated as the number of disorders it is comorbid with in a patient population, is sensible enough a measure of its “total comorbidity” (Hidalgo et al. 2009) and a useful concept both epidemiologically and clinically. The weights (using BAMs) associated with these comorbid disorders are also clearly useful for discriminating between stronger and weaker co-occurrence rates, hence higher or lower risk factors for patients with an index disorder. However, we found that the choice of disease ontology has a significant impact on comorbidity rankings, so much so that the centralities of disorders before crosswalking to a coarser ontology are not predictive of the centralities of their counterparts after crosswalking. (See the Results section for more detail.)

Meanwhile, none of the weights commonly used to quantify the strengths of comorbidities are *additive*: For example, using one typical network construction,^2^ amebiasis, a gastrointestinal infection rare in the United States, and rheumatoid arthritis, a common chronic autoimmune disorder, have 5 and 67 comorbid relations, respectively. Though having very different etiologies and afflicting very different patient populations, these disorders have approximately the same weighted degree (529 and 561).

This does not translate to their being similarly severe in any recognized sense, or to their belonging at a similar ranking amidst the other disorders in the ontology.

Betweenness and closeness centrality rely on a different version of additivity that is equally problematic. Take the same network, except invert the link weights^3^ to indicate distance rather than strength of association. The geodesic distance between type 1 diabetes and breast cancer (in female patients) is then the same as that between multiple epiphyseal dysplasia (MED) and hepatitis E (HepE), approximately .30, though the former two disorders are significantly correlated (i.e. directly linked, *p* < 10^−27^) while the latter two can only be reached from each other via three intermediate disorders: multiple epiphyseal dysplasia ↔ Albright–Sternberg syndrome ↔ cerebral palsy ↔ hepatitis C ↔ HepE. Yet this indirect sequence of associations leading to HepE is does not have an established clinical interpretation, nor does it imply a natural comparison to the relative risk of MED encoded by the direct link. Indeed, controlling for covariates and subsetting populations may significantly alter the magnitudes and signs of the associations in the sequence, with unpredictable effects on the resulting indirect distance measure.

These limitations are better revealed by weighting schemes, but they arise from the network model itself, which is premised on a principle of “guilt by association” that implicates one node in the effects of another according to their proximity in the network. As Hou et al. (2014) point out in the related field of genomics, this principle “does not reflect the dynamic nature of biological networks”. The same may be said of epidemiological networks. As in genomics, comorbidity network centrality analysis is demonstrably effective at prioritization, but without underlying theory or validation it will be difficult to know what critical diagnoses it may fail to identify.

Such concerns are not specific to comorbidity network analysis. Inconsistency and uncertainty over the interpretations of centrality measures in the study of human communication networks led Freeman (1978) to propose the concise set of conceptualizations and measures discussed above: *degree*, based on the idea of communication activity with other nodes; *betweenness*, based on the control of communication among other actors, and *closeness*, based on either independence from the control of others or efficiency of dissemination. These interpretations can be naturally extended to other kinds of resource exchange, but they do not have straightforward interpretations on correlation networks. Such interpretations are needed if disorders are to be characterized with respect to specific types of centrality—for example, the betweenness centrality of acute posthemorrhagic anemia in the MIMIC-III intensive care population—rather than merely being observed to be centrally located in the network in a non-specific sense.

### 4.4 Recommendations

We enumerate here some recommendations for future comorbidity network studies. To pre-empt and mitigate the aforeraised concerns, in light of our observations, we urge CNA researchers to include in their analysis, according to their aims and methods, several steps:

- Make patient–diagnosis incidence data available for secondary use. This enables future researchers to perform multivariate analyses that are essential for our understanding of epidemiological comorbidity, as will be increasingly recognized as comorbidity network analysts leverage the gains already made by ecologists and psychologists. The release of such data will require additional processing work, some of it entailed by privacy requirements, e.g. the removal of low-frequency disorders or unique patient profiles; but, the sooner these tasks are pioneered, the sooner they can be more widely adopted. If patient–diagnosis data cannot be released, then at least pairwise frequency tables, which enabled our sensitivity analysis, should be.
- Provide theoretical justification for the choice of disorder ontology. The clinical ontologies used by CNA data sets were designed for different uses, e.g. treatment versus billing versus research, and differences between them due to geography and time as well as purpose are unavoidable. Global and local properties may vary dramatically between networks constructed from the same incidence data before versus after crosswalking to an alternative ontology. If a coarser ontology more accurately identifies patient cohorts that have a well-defined index disorder, then the comorbidity network should be constructed after conversion from the original ontology.
- Provide theoretical justifications for weighted CNAs. Indirect network relations such as those underpinning most centrality measures are highly sensitive to the choice of association measure, and the conventional interpretations of these measures at the pairwise level do not translate naturally to the network setting. If weighting is important but no specific weight is entailed, then obtain and report results obtained using multiple weight types—e.g. unit, relative risk–like, or correlation.
- To summarize the global topology of a comorbidity network, report the size of its largest component, its degree assortativity, and its global clustering coefficient, in addition to any other desired summary statistics. These statistics are commonly calculated and effectually discriminate between comorbidity networks. They are therefore especially appropriate to include in comparisons of networks for different populations or disorders of interest.
- Provide theoretical justification for taking a pairwise versus a partial correlation approach. Controlling for confounding effects among all disorders in an ontology can radically change the structure of a comorbidity network, in particular revealing a large number of negative associations. Since free software allows correlation matrices to be efficiently converted to partial correlation matrices, this approach is not prohibitively more expensive.
- Use joint distribution or other multivariate models to validate associations among the most common disorders or within small subsets of disorders. This modeling approach may be too costly for whole-diseaseome analysis, and it may fail to recover comparatively weak but highly discernible comorbidities outside a core set of more prevalent disorders, but it may help distinguish and prioritize the associations of greatest importance to the interactions among a subset of interest, for example the known risk factors and complications of an index disorder.

In addition to these methodological recommendations, we stress the importance of involving clinicians in CNA, to assist both in formulating network-analytic goals or hypotheses that are meaningful and relevant and in understanding the (usually administrative) data on which the analysis is performed. Many CNA studies to date have been (co-)authored by clinicians, other physicians, or medical researchers, and medical background knowledge is evident and crucial in the discussions of several seminal analyses. For example, in one of the earliest CNAs, Hanauer et al. (2009) parse the medical plausibility of different explanations for the novel associations they uncovered and provide an essential discussion of likely sources of inconsistency. In a different type of example altogether, Schafer et al. (2014) develop a novel network construction based on triads, motivated by a three-disease conception of multimorbidity, that eliminated much of the low-relevance periphery of association networks but that has not to our knowledge been used elsewhere. As applications proliferate, it will be increasingly important to ensure that clinical authors bring their training and experience to bear on these projects.

### 4.5 Future work

While previous work has compared different false discovery rate corrections on network data at the pairwise level (Koo et al. 2014), we are not aware of any research, theoretical or empirical, into whether FWER-versus FDR-based corrections more faithfully or usefully recover network properties such as centrality and clustering. Therefore, while we observe some dependence of these properties on the choice of correction (if any), we cannot recommend any method over any other. To inform this choice, future work should evaluate such corrections on the recovery of network properties based on incidence data generated from realistic covariance matrices and on the ability to draw useful inferences from network models of real-world incidence data.

The network analytic approaches used here represent only a subset of the broader range of systems analytic techniques, which includes structural equation models and (discrete or continuous) dynamical models. Few if any studies of disease incidence and co-occurrence have tested the agreement of inferences drawn from such methods with respect to a common topic or question, whether applied to the same or independently collected data. These more computationally intensive methods benefit from the target-pruning of network analysis, and it is important to ensure that potentially important targets are not missed.

## 5 Methods

### 5.1 Pairwise constructions

To address the first two concerns from Section 2.3, we test the dependence of degree distributions, global network statistics, and node centrality rankings to the source of data, the method of link determination, and the choice of network tools. Our results speak to the generalizability of such properties.

#### 5.1.1 Data sets

The data sets in Table 1 were pre-processed and made available by the authors of previous comorbidity network studies (Rzhetsky et al. 2007; Hidalgo et al. 2009; Roque et al. 2011; Hanauer and Ramakrishnan 2013; Bagley et al. 2016), except for MIMIC-III (which is freely available for research use) (Johnson et al. 2016). The data sets vary widely in the underlying patient population, in the collection of their data by healthcare institutions, *and* in the researchers’ pre-processing protocols; variation along each of these dimensions contributes to overall variation due to the data source. Disentangling these factors would require a more thorough study using several sources of patient-level data, and we make no attempt to do so here.

**Table 1:**
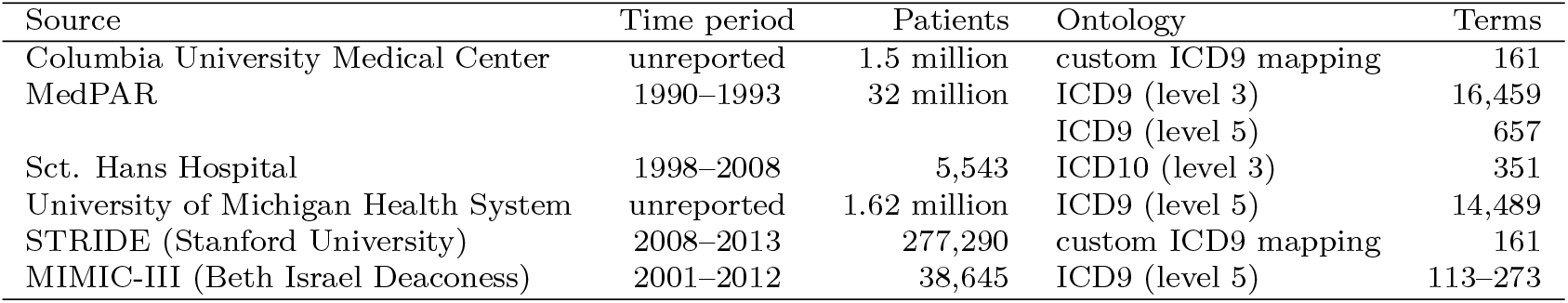
Sources of pairwise disorder co-occurrence data used in this study, originally aggregated from patient-level data for previous studies and made available by their authors (except MIMIC-III).

#### 5.1.2 Link determination

The majority of comorbidity networks in our surveyed literature (Brunson and Laubenbacher 2017) have been aggregated from pairwise co-occurrence data, meaning that they can be constructed from the entries *a, b, c, d* in the 2 × 2 contingency tables for disorder pairs (*D*_1_, *D*_2_):

| | $D_2$ | $\neg D_2$ |
| --- | --- | --- |
| $D_1$ | $a$ | $b$ |
| $\neg D_1$ | $c$ | $d$ |

We calculated pairwise p-values via Fisher’s exact test,^4^ optionally adjusting for multiple comparisons using the family-wise error rate (FWER) Bonferroni correction or the false discovery rate (FDR) Benjamini–Hochberg correction, both of which have been used in the CNA literature (Roque et al. 2011; Roitmann et al. 2014; Bhavnani et al. 2015; Bagley et al. 2016; Kim et al. 2018).

In addition to the option of leaving links unweighted (the “unit” measure) we adopted four measures of binary association: two risk ratios and two correlation coefficients The *odds ratio* 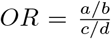 (Kraemer 1995; Parzen et al. 2002); *Pearson’s binary correlation coefficient* 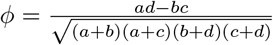 (Hubalek 1982), *Forbes’ coefficient of association* 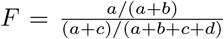 (Hubalek 1982), and the *tetrachoric correlation coefficient r_t_* calculated using a latent bivariate normal model (Drasgow 2006).

*OR* is recommended as a standard measure of epidemiological comorbidity (Kraemer 1995) and has been used in several CNA studies (Hanauer et al. 2009; Hanauer and Ramakrishnan 2013; Kim et al. 2016). Several other studies have used *ϕ* and *F* together, in part to check the robustness of their results (Hidalgo et al. 2009; Folino and Pizzuti 2012; Chen and Xu 2014; Chmiel et al. 2014; Lai 2015), though these misleadingly refer to *F* as “relative risk” (Zhang and Yu 1998). *OR, ϕ*, and *F* have been used in previous CNA studies. *r_t_* complements the mix as a second correlation coefficient and provides methodological continuity with the later multivariate models. Each measure is symmetric, so that the roles of *b* and *c* in the above table are interchangeable.

Each comorbidity network was thus constructed based on five parameters: the source of data *D*; the evidential cutoff (TWER) *α*;, the choice of error rate correction *C*, if any; the BAM *m*; and an optional evaluative (BAM) cutoff *θ*. We use the notation *N*(*D, α, C, m, θ*) to denote specific networks and substitute bullets • for values to indicate families of networks taken over all values in the following ranges:

- *D*: Columbia, MedPAR(3), MedPAR(5), Sct. Hans, Michigan, Stanford, Columbia*, MIMIC
- *α*: 10^−*p*^, *p* = 1,…, 6
- *C*: none (Ø), Bonferroni (*B*), Benjamini–Hochberg (*BH*)
- *m*: 1 (unit), *OR, ϕ, F, r_t_*
- *θ*: each of four values specific to each measure:^5^ *θ_OR_* = 1, 2, 6, 60, *θ_ϕ_* = 0, 0.005, 0.05, 0.2, *θ_F_* = 1, 2, 6, 60, *θ_r_t__* = 0, 0.1, 0.4, 0.6

#### 5.1.3 Power laws and scale-freeness

Like many other empirical networks, comorbidity networks have been described as following a power law, and thereby being scale-free, usually based on visual inspections of their degree distributions (Steinhaeuser and Chawla 2009; Ball and Botsis 2011; Divo et al. 2018). Other studies have taken scale-freeness as the basis for using pre-processing steps designed for scale-free networks (Jiang et al. 2018) or testing whether such networks are consistent with the preferential attachment growth model (Scott et al. 2014). However, rigorous definitions of power-law behavior in empirical data are contested (Broido and Clauset 2018; Voitalov et al. 2018), and neither the visual methods often used to determine scale-freeness (Li et al. 2005; Clauset et al. 2009) nor the implications that power-law degree sequences are often thought to have for the structural and generative properties of a network (Willinger et al. 2004; Mitzenmacher 2004; Li et al. 2005) are reliable. (One CNA that applied formal statistical methods to test for power-law degree sequences drew negative conclusions (Davis and Chawla 2011).)

We began by testing whether comorbidity networks have power-law degree sequences, using two methodologies that have come to represent the terms of the debate in the network literature. Following the methodology of Clauset et al. (2009) and Gillespie (2015), we fit four pure models— Poisson (expected of Bernoulli random graphs), power law, exponential, and log-normal—to the degree sequence tails of the unweighted networks *N*(•, 0.05, •) and compare the goodnesses of fit of the model families, statistically and visually. Then, following the methodology of Voitalov et al. (2018) and Voitalov (2018), we assumed the degree sequences were drawn from regularly varying distributions and compare several estimators of the power-law exponent for well-definedness and consistency.

#### 5.1.4 Global structure of comorbidity

To assess the effects of the construction parameters on the resulting network structure, we calculated a battery of single-valued global statistics on the unweighted networks *N*(•, •, •): the proportion of nodes in the largest connected component “LCP”, the graph density *δ*, the mean node degree 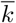, the Gini index *G* of the degree sequence (Badham 2013), the degree assortativity *r* (Newman 2003), global triad closure *C*, the average internode distance 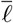, the modularity *Q* based on a Walktrap node partition (Newman and Girvan 2004; Pons and Latapy 2006), and the maximum-likelihood estimates of the parameters of the model family found to best fit the most degree sequence tails, which turned out to be the parameter estimates 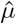 and 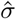 of the log-normal family.

For each statistic *s*, we fit two families of multiple linear regression models to its values *s_i_* at the networks *N_i_* = *N*(*D_i_, α_i_*, Ø, *m_i_, θ_i_*). In each case we take as the response variable the difference 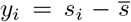 between *s_i_* and the average value 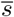 on the networks *N*(•, •, Ø). In this way, the effects of both kinds of cutoff indicate the direction and magnitude of change in the statistic as more strict evaluative cutoffs are imposed. The first model (Equation 1) regresses *y_i_* on the data source *d_i_*, treated as categorical, and log(*α_i_*), treated as continuous, using only data from networks *N*(*D_i_, α_i_*, Ø). We omit an intercept term, so that each dataset has an associated coefficient that indicates the direction in which *s* deviates, on this population, from its values on the others. The second model (Equation 2) additionally includes interaction terms between the BAMs and evaluative cutoffs and is fit to the values on the networks *N*(•, •, Ø, •, •). These terms encode the fact that restricting cutoffs moves each BAM-specific family of networks away from a common origin.^6^

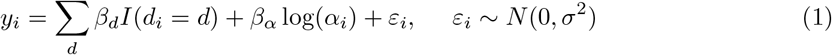

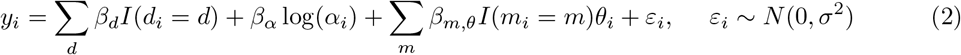

We complemented these regression models with a principal components analysis (PCA) on the values of all 10 centered and scaled statistics measured across *N*(•, •, Ø, •, •). A row-principal biplot of the data points and variable axes along the first two principal components (PCs) characterizes the dimensions of greatest variance in terms of the loadings of the global statistics.

#### 5.1.5 Centrality rankings of disorders

Network models enable researchers to characterize the structural positions of individual nodes (Brandes 2016). Analyses of comorbidity networks have usually invoked any of three standard measures of centrality developed for the analysis of social networks (Freeman 1978): *degree*, the number of actors linked to an index actor; *betweenness*, the proportion of geodesics (shortest paths) connecting other actors that pass through an index actor; and *closeness*, the reciprocal sum of the geodesic distances (lengths of shortest paths) to an index actor.

Several analyses have characterized disorders by their centrality in a comorbidity network: Hidalgo et al. (2009) and Schafer et al. (2014) used (weighted and unweighted, respectively) degree centrality to measure the connectedness of disorders, which we can think of as their “total” epidemiological comorbidity; Schafer et al. (2014) also used betweenness centrality to measure the potential influence of an index disorder on a patient’s comorbidities. Several other teams invoked degree, betweenness, and closeness centrality as general indicators of a disorder’s importance to the larger diseaseome (Ball and Botsis 2011; Chen and Xu 2014; Liu et al. 2016). In terms of specific results, three studies corroborated the exceptionally high centrality of hypertension in comorbidity networks (Chen and Xu 2014; Schafer et al. 2014; Feldman et al. 2016), while another examined the centralities of disorders comorbid with hypertension (Liu et al. 2016). An increasingly popular approach is to examine differences in disease centrality across study populations: Feldman et al. (2016) compared the betweenness centralities of central diagnoses between demographic subgroups such as low- and high-income populations, and Divo et al. (2018) compared the degree centralities of disorders between COPD and non-COPD populations in a case–control design.

Our sensitivity analysis concerns the effects of link determination on disease centralities and centrality rankings. These effects may be qualitatively different from the effects on global statistics described in the previous section. If changes in the thresholds have pronounced effects on global network statistics, this might be understood primarily as an effect of changes in network density. Certainly, changes in density—the loss or gain of a significant proportion of links—will affect node centralities, but *relative* centralities could nevertheless remain stable. This has been tested in a few instances, with significant changes in rankings being observed: Feldman et al. (2016) found that the betweenness rankings of disorders were sensitive to their link pruning procedure, and Ball and Botsis (2011) noted that the centralities of adverse events in their VAERS networks changed noticeably from month to month.

To measure this affect systematically, we calculated degree, betweenness, and closeness centralities in the networks *N*(•, 0.05, •, •, −∞). For the strength-based measure (degree), we weighted each link by its BAM. For the path-based measures (betweenness and closeness), we weighted each link by the reciprocal of its BAM, so that stronger associations yield shorter internode distances. We imposed no evaluative threshold. For each fixed data source and centrality measure, we compared the disorder rankings obtained using each p-value correction and each BAM using Kendall rank correlations (Kendall 1938, 1945) as a test of the robustness of the approach. We analyzed the rankings geometrically via eigendecompositions of the pseudo-correlation matrices of Kendall values. We report the proportions of inertia explained by each construction parameter and summarize the correlations among the rankings using biplots.^7^

It may be that overall centrality rankings are sensitive to network construction while the identification of exceptionally central “hubs”, the focus of most applications, is robust: nodes on the periphery tend to be structurally similar and of lower degree, so that the addition or removal of fewer links would be needed to permute their ranks. It is also important, for the generalizability of CNA, to know whether the high centrality of certain disorders is an artifact of the network construction, a robust property of a specific patient population, or a generalizable epidemiological fact. To address these “hubs” specifically, we inspected the several most central disorders from each network.

Any specific ontology was only used by two to four data sets. In order to test the robustness of centrality rankings across data sets with different ontologies, we used many-to-one maps from finer to coarser ontologies to identify groups of nodes in finer comorbidity networks with single nodes from coarser networks. Where such mappings were available (level-5 to level-3 ICD9 and level-5 ICD9 to the ontology of Rzhetsky et al. (2007)), we compared group centrality measures for concepts in the finer ontology with node centralities for concepts in the coarser ontology (Everett and Borgatti 1999). Because group betweenness centrality was computationally prohibitive, we only considered degree and closeness centrality.

### 5.2 Multiple and multivariate constructions

Our sensitivity analysis of pairwise comorbidity networks was enabled by the several research teams who shared their pairwise data. With access to patient-level incidence data, advanced statistical software, and adequate computational resources, a wider family of network models becomes available. Multivariate models that incorporate all disease associations simultaneously can address concerns of multiple comparisons and confounding without resort to post hoc corrections. In this phase, we compare comorbidity relationships obtained from two publicly-available comorbidity datasets using the conventional pairwise construction, a partial correlation construction that corrects for confounding, and a recently-proposed multivariate model that also incorporates patient-level covariates. The comparisons will reveal the effects of systems-level interactions on observed pairwise relations.

#### 5.2.1 Datasets

We draw from two data sets, one low-dimensional but high-volume, in the sense of covering the incidence of a small number of disorders for a large number of patients, and one comparatively high-dimensional and low-volume.

As a low-dimensional use case, we use data for the year 2011 from the National Ambulatory Medical Care Survey (NAMCS), coordinated by the CDC and conducted by hundreds of physicians and community health centers each year (https://www.cdc.gov/nchs/ahcd/index.htm). Each entry describes a single encounter, and each physician collects data for two weeks out of the year, *without* linking records for encounters with the same patient. Thus, the data does not contain long-term data for any patient, and any multiple diagnoses were recorded at a single encounter. The survey data include a range of patient demographics, provider characteristics, and clinical information such as diagnoses and procedures pertaining to the encounter. While these diagnosis data are suitable for certain study designs, e.g. cross-sectional descriptions of specialty- or location-specific patient populations, they are inappropriate for comorbidity network analysis, which relies on the recovery of comprehensive health profiles for all patients in a sample.

NAMCS asks providers to indicate whether each patient has each of several chronic disorders, including thirteen that are reported in the Public Use Dataset: arthritis, asthma, cancer, cerebrovascular disease, chronic obstructive pulmonary disease, congestive heart failure, ischemic heart disease, depression, diabetes mellitus, hyperlipidemia, hypertension, obesity, and osteoporosis. These are widely-recognized conditions that do not require specialized training to diagnose and that will often already appear on patients’ records. Therefore, despite the limitations of NAMCS, we expect that the data much more reliably reflect the actual distribution and co-occurrence of these chronic disorders in the clinic-going population. To make fitting joint distribution models (see Section 5.2.3) more computationally feasible, we took a 50% cluster sample from this year of NAMCS encounter data, after restricting to cases for which all variables were recorded, clustered by participating practices and weighted by number of encounters within the practice.

As a contrast to NAMCS, we draw a second data sample from MIMIC-III, described previously. MIMIC-III is similar to NAMCS in containing only one encounter for most patients, but dissimilar in covering a narrow range of practice and therefore a limited population. To increase the clinical homogeneity of our sample, we restrict our attention to patients admitted to specific units, which include medical intensive care (MICU), surgical intensive care (SICU), the neonatal ward (NWARD), neonatal intensive care (NICU), coronary care (CCU), cardiac surgery recovery (CSRU), and trauma/surgical intensive care (TSICU). For computational feasibility, we grouped the ICD-9 diagnosis codes recorded in MIMIC-III according to the Clinical Classification Software (CCS) ontology (Agency for Healthcare Research and Quality 2012). Care unit populations ranged in the number of distinct recorded CCS categories from 113 (NICU) to 273 (MICU).

#### 5.2.2 Partial correlation networks

Conventional measures of comorbidity fail to account for an important source of variation in patient-level incidence data: incidence rates of other disorders. Two disorders that are clinically unrelated may be epidemiologically comorbid due to having risk factors or complications in common, and CNA researchers have pointed out that these are important potential explanations for clustering patterns observed in comorbidity networks (Hanauer et al. 2009). Partial correlations account for these confounding effects by generalizing the calculation of correlations from regression coefficients: the full partial correlation 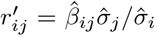 between response variables *y_i_* and *y_j_* is a standardized effect estimate from the regression model of *y_i_* on all other responses, including *y_j_* (Epskamp and Fried 2017). This concept relies on the normality assumptions of classical regression; we use a matrix formulation to obtain a matrix 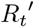 of *partial tetrachoric correlations* 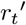 from the matrix *R_t_* of tetrachoric correlations.

#### 5.2.3 Joint distribution networks

Epidemiological comorbidities can also arise from patient-level covariates, as when clinically relevant subpopulations (e.g. elderly or infirm patients) are at heightened risk of multiple, otherwise etiologically unrelated disorders. Several comorbidity network studies incorporated demographic predictors into their binary association tests (Rzhetsky et al. 2007; Feldman et al. 2016; Glicksberg et al. 2016), though this information was unavailable for our sensitivity analysis. To simultaneously account for the *endogenous* effects of other disorders and the *exogenous* effects of patient-level covariates, we adapted the *joint distribution model* (JDM) designed to incorporate both species interactions and environmental factors into ecological models (Pollock et al. 2014).

In making this choice, we propose and appeal to an *ecological–epidemiological analogy:* Disorders afflicting persons and communities are similar in many respects to species occupying geographical sites. In the case of viral, bacterial, and fungal infections, the former is in fact a special case of the latter. Insofar as the assumptions underlying an ecological analytic technique are met by epidemiological data, the technique is an appropriate one. The analogy posits that this will frequently be the case, or at least that ecological techniques will be no less appropriate than competing ones. Indeed, association network analysis itself is rooted in ecology, which produced many if not most of the measures commonly used to weight association networks (Johnston 1976; Podani 2000). More recently, ecologists have honed several other methods to account for the same limitations of pairwise network construction discussed here (Ulrich and Gotelli 2007; Ovaskainen et al. 2010; Cazelles2016; Morueta-Holme et al. 2016); see Elith and Leathwick (2009) and Kissling et al. (2012) for useful reviews.

In the JDM, each patient’s distribution has the same covariance matrix, from which a correlation matrix P = (*ρ_ij_*) is obtained, and is centered at a linear transformation B = (*β_ki_*) of their *q* exogenous variables (1 ≤ *k* ≤ *q*), which can be interpreted in the same way as classical regression coefficients. The latent normality allows us to directly compare the estimated endogenous correlations 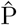 to *R_t_* and 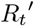, and the combined estimated effects of the patient-level covariates can be organized into a matrix 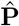 of exogenous components of correlation. We fit the JDM to the *n* × *m* patient-disorder incidence matrices, using either of two matrices *X* of exogenous patient-level covariates: an *n* × 1 intercept matrix to encode only the baseline prevalence of each disorder, and an *n* × *q* matrix augmented with demographic variables. The respective models are designated JDM0 and JDM1 and estimates indexed with 0 or 1 accordingly. See the SI Text for more details.

#### 5.2.4 Correlation structure

We first examined the differences in correlation structure between models produced using these three approaches to comorbidity, independently of link determination. From the NAMCS data, we generated four correlation matrices for the 13 chronic disorders described in Section ref{sec:datasets-multivar}: pairwise (*R_t_*), full partial 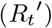, and joint distribution with prevalence-only and with full demographic exogenous covariates (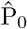 and 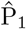). In the last model, the exogenous variables included decadal age groupings (0–14, 15–24, etc., to 65–74, with 75+ the baseline), gender (male, with female the baseline), race/ethnicity (Asian, Black/African-American, American Indian/Alaska Native/Pacific Islander, white, and Hispanic/Latinx, with unknown/other the baseline), insurance status (Private, Medicare, and Medicaid, with others or none the baseline) region (Midwest, South, West, with Northeast the baseline), and metropolitan status (Metropolitan Statistical Area, with non-MSA the baseline). From the MIMIC data, we only considered the three endogenous models (pairwise, full partial, and joint distribution without exogenous covariates), in part to reduce computational cost and in part to limit the scope of the analysis. Each model included every CCS code.

To assess the effect of moving from a pairwise to a systemic approach to comorbidity, we focused on differences among *R_t_*, 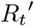, and 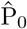. The additional effect of controlling for patient-level covariates manifested in differences between 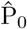 and 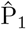. We compared the correlation matrices *R_t_*, 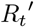, 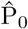, 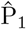 among the 13 chronic disorders in NAMCS using correlation biplots. We visualized the relationships among the point estimates of association strength *r_t_*, 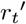, 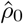, 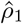, across all 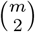 pairs disorders from each data source, using scatterplots.

#### 5.2.5 Link determination

Differences in correlation structure translate into differences in network structure through the link determination process. To evaluate these differences, we adopted a uniform evidential threshold of *α* = 5% by which to categorize the pairwise associations in each model as discernibly positive, discernibly negative, or indiscernible. We sought patterns of differences between the pairwise determinations using alluvial diagrams (Brunson 2018) and visualized the resulting networks using a circular layout.

#### 5.2.6 Centrality rankings of disorders

To evaluate differences in the resulting network structures, we compared centrality rankings as in Section 5.1.5, using Kendall rank correlations between the different network models of a common data set and visualizing them using pseudo-correlation biplots.

### 5.3 Software

We performed our analyses using the R statistical programming language (R Core Team 2016), unless stated otherwise. We processed and analyzed data using packages from the tidyverse collection (Grolemund 2016) and manipulated and visualized networks using the igraph (Csardi and Nepusz 2006), tidygraph (Pedersen 2018a), and ggraph (Pedersen 2018b) packages.

Depending on the application, we used the implementation of *r_t_* in the psych package (Revelle 2017) or implemented the approximation method of Bonett and Price (2005) in order to avoid infinities and to calculate standard errors. We also used different implementations to compute partial correlations *R′* from pairwise correlations *R*: For high-volume, low-dimension data, we used psych package implementation `partial.r()`, which uses squared multiple correlations. For low-volume, high-dimension data, this became infeasible, and we first calculated shrinkage estimates (Schäfer and Strimmer 2005) for *R′* using `cor.shrink()`, together with the partial correlation function `cor2pcor()`, from the corpcor package (Schäfer et al. 2017).

We organized effect estimates from regression models using the stargazer package (Hlavac 2015). Biplots were rendered using the ordr package (Brunson 2019). English descriptions of codes from the International Classification of Diseases, 9th and 10th Revisions, Clinical Modifications (ICD9 and ICD10), were obtained from and formatted using the icd package (Wasey 2017).

We adapted our JDM workflow from the tutorial provided by Pollock et al. (2014), using JAGS (Plummer 2003) to perform Bayesian model fitting through the R2jags package (Su and Yajima 2015).

Full code to reproduce our analyses will be made available on Bitbucket upon acceptance for publication.

### 5.4 Ethical considerations

This study did not involve human or other animal subjects. We conducted secondary analysis on data sets collected and aggregated by other researchers, which are available either publicly or upon request. Of these, patient-level data were only available in the MIMIC-III database, but our analysis relied exclusively on aggregated data.

## 6 Results

### 6.1 Pairwise analysis

#### 6.1.1 Link determination

Table 2 presents network density for each dataset and each of several common TWERs. In most cases the original authors provided co-occurrence data only for a subset of pairs, according to their own study designs; the network density at the 100% TWER is the proportion of pairs they included. Often among these were data on negative associations. In each case our 5% TWER cutoff excluded many additional pairs, including all negative associations. The Bonferroni correction excludes the vast majority of the remaining links from networks for which more pairs were originally available (MedPAR(5), MIMIC) but fewer than one third from networks that had already been pruned of weak associations (Sct. Hans, Stanford, Columbia*). For any fixed evidential cutoff, the networks range in density over 1–3 orders of magnitude. Quintiles calculated for each BAM and data source indicated that the evidence for a comorbid association may not be predictive of its strength, so that some network properties may be more sensitive to an evaluative cutoff than to an evidential one (see the SI Text).

**Table 2:** Densities (proportion of pairs of nodes that are linked) of comorbidity networks constructed using different evidential thresholds. Family-wise error rate (FWER) correction uses the Bonferroni procedure; false discovery rate (FDR) correction uses the Benjamini–Hochberg procedure.

| p-value | Correction | Columbia | MedPAR(3) | MedPAR(5) | Sct.Hans | Michigan | Stanford | Columbia* | MIMIC |
| --- | --- | --- | --- | --- | --- | --- | --- | --- | --- |
| 1 | none | 0.624 | 0.560 | 0.027 | 0.029 | 0.083 | 0.073 | 0.110 | 0.040 |
| 1 | FWER | 0.174 | 0.156 | 0.002 | 0.025 | 0.048 | 0.065 | 0.104 | 0.001 |
| 1 | FDR | 0.451 | 0.465 | 0.006 | 0.029 | 0.077 | 0.073 | 0.110 | 0.009 |
| 0.01 | none | 0.240 | 0.244 | 0.006 | 0.029 | 0.073 | 0.073 | 0.110 | 0.009 |
| 0.01 | FWER | 0.142 | 0.138 | 0.002 | 0.013 | 0.042 | 0.054 | 0.089 | 0.001 |
| 0.01 | FDR | 0.210 | 0.215 | 0.003 | 0.029 | 0.067 | 0.068 | 0.109 | 0.002 |
| 0.0001 | none | 0.175 | 0.181 | 0.003 | 0.029 | 0.063 | 0.064 | 0.102 | 0.003 |
| 0.0001 | FWER | 0.126 | 0.125 | 0.001 | 0.008 | 0.037 | 0.046 | 0.081 | 0.001 |
| 0.0001 | FDR | 0.161 | 0.168 | 0.002 | 0.015 | 0.057 | 0.057 | 0.093 | 0.001 |
| 0.000001 | none | 0.144 | 0.153 | 0.002 | 0.014 | 0.054 | 0.053 | 0.089 | 0.002 |
| 0.000001 | FWER | 0.113 | 0.115 | 0.001 | 0.006 | 0.033 | 0.040 | 0.073 | 0.001 |
| 0.000001 | FDR | 0.136 | 0.145 | 0.002 | 0.009 | 0.049 | 0.047 | 0.082 | 0.001 |

#### 6.1.2 Degree sequence distributions

Almost all comorbidity networks were better-modeled by log-normal distributions, and then by exponential distributions, than by power-law distributions. Based on our sample, this pattern was not dependent on the construction parameters, except that the preference for log-normal versus exponential versus power-law distributions were less clear for networks constructed using stricter evidential thresholds. Most regular variation–based power-law models failed to converge or else were inconsistent, reaffirming that power-law models were inappropriate for these networks’ degree sequences. See the SI Text for details.

#### 6.1.3 Single-valued summary statistics

Table 3 reports the effect estimates obtained by fitting Equation 2 to the various global network statistics. The coefficients within each model (column) associated with the data sources can be directly compared using differences, though the scale is interval, not ratio (i.e. the relative position of 0 is arbitrary). The evaluative cutoffs *θ_m_* are included only as interaction effects with a categorical variable encoding the measure *m*, because the range of values of *θ_m_* varied across *m*. Roughly, the effect estimates can be compared after scaling by the ranges of the *θ_m_* (Section 5.1.2). For example, the relative effect estimates of 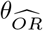 and *θ_r_t__* on average degree 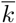 were in proportion to –1.14 × 59: –125.11 × 0.6, or 9: 10. The effect estimates obtained using Equation 1 were qualitatively similar, for those predictors included in both models (Table S6).

**Table 3:** LMs of network statistics on data source, test-wise error rate, and binary association measure.

|  | *Dependent variable:* |  |  |  |  |  |  |  |  |
| --- | --- | --- | --- | --- | --- | --- | --- | --- | --- |
| | LCP<br>(1) | $r$<br>(2) | $G$<br>(3) | $k$<br>(4) | $\hat{\mu}$<br>(5) | $\hat{\sigma}$<br>(6) | $\ell$<br>(7) | $Q$<br>(8) | $C$<br>(9) |
| Columbia | 0.24***<br>(0.03) | 0.06*<br>(0.03) | -0.19***<br>(0.02) | -51.04***<br>(11.46) | -0.001<br>(0.18) | -0.34***<br>(0.05) | -0.68*<br>(0.27) | -0.03<br>(0.03) | 0.01<br>(0.02) |
| MedPAR(3) | 0.31***<br>(0.03) | 0.13***<br>(0.03) | -0.18***<br>(0.02) | -31.79**<br>(11.46) | 0.31*<br>(0.17) | -0.08<br>(0.05) | 0.30<br>(0.27) | 0.12***<br>(0.03) | -0.09***<br>(0.02) |
| MedPAR(5) | 0.21***<br>(0.03) | -0.12***<br>(0.03) | 0.03*<br>(0.02) | -39.54***<br>(11.46) | 0.24<br>(0.17) | 0.31***<br>(0.05) | 1.38***<br>(0.27) | 0.18***<br>(0.03) | -0.28***<br>(0.02) |
| Sct.Hans | 0.13***<br>(0.03) | 0.04<br>(0.03) | -0.11***<br>(0.02) | -59.58***<br>(11.46) | -1.16***<br>(0.18) | 0.03<br>(0.05) | 0.20<br>(0.27) | 0.10**<br>(0.03) | -0.22***<br>(0.02) |
| Michigan | 0.51***<br>(0.03) | -0.18***<br>(0.03) | -0.11***<br>(0.02) | 158.43***<br>(11.46) | 2.07***<br>(0.17) | 0.23***<br>(0.05) | 0.31<br>(0.27) | 0.12***<br>(0.03) | -0.12***<br>(0.02) |
| Stanford | -0.21***<br>(0.03) | 0.39***<br>(0.03) | -0.07***<br>(0.02) | -60.66***<br>(11.46) | -2.20***<br>(0.19) | 0.23***<br>(0.05) | -1.50***<br>(0.27) | -0.13***<br>(0.03) | 0.09***<br>(0.02) |
| Columbia* | -0.19***<br>(0.03) | 0.07*<br>(0.03) | -0.14***<br>(0.02) | -58.96***<br>(11.46) | -0.93***<br>(0.19) | -0.45***<br>(0.05) | -1.63***<br>(0.27) | -0.14***<br>(0.03) | 0.01<br>(0.02) |
| MIMIC | 0.31***<br>(0.03) | 0.02<br>(0.03) | -0.07***<br>(0.02) | -37.21**<br>(11.46) | -0.09<br>(0.17) | 0.43***<br>(0.05) | 1.62***<br>(0.27) | 0.15***<br>(0.03) | -0.21***<br>(0.02) |
| $\log(p)$ | 0.02***<br>(0.002) | -0.003<br>(0.002) | -0.01***<br>(0.001) | 1.95*<br>(0.80) | 0.06***<br>(0.01) | -0.01**<br>(0.003) | -0.05**<br>(0.02) | -0.01***<br>(0.002) | -0.01***<br>(0.001) |
| $F \times \theta_F$ | -0.01***<br>(0.001) | 0.001*<br>(0.001) | 0.002***<br>(0.0003) | -1.17***<br>(0.20) | -0.04***<br>(0.004) | -0.003*<br>(0.001) | 0.02***<br>(0.005) | 0.002***<br>(0.001) | -0.003***<br>(0.0003) |
| $\widehat{OR} \times \theta_{\widehat{OR}}$ | -0.01***<br>(0.001) | 0.003***<br>(0.001) | 0.002***<br>(0.0003) | -1.14***<br>(0.20) | -0.03***<br>(0.004) | -0.003**<br>(0.001) | 0.02***<br>(0.005) | 0.003***<br>(0.001) | -0.003***<br>(0.0003) |
| $\phi \times \theta_\phi$ | -3.24***<br>(0.15) | 2.20***<br>(0.17) | 1.46***<br>(0.09) | -422.14***<br>(59.45) | -19.35***<br>(1.08) | 0.47<br>(0.30) | 3.70**<br>(1.38) | 0.57***<br>(0.16) | 0.51***<br>(0.11) |
| $r_t \times \theta_{r_t}$ | -0.63***<br>(0.05) | 0.19***<br>(0.05) | 0.19***<br>(0.03) | -125.11***<br>(17.70) | -4.50***<br>(0.30) | -0.04<br>(0.08) | 2.64***<br>(0.41) | 0.34***<br>(0.05) | -0.22***<br>(0.03) |
| Observations | 576 | 568 | 576 | 576 | 504 | 504 | 576 | 576 | 568 |
| Adjusted R<sup>2</sup> | 0.78 | 0.59 | 0.57 | 0.66 | 0.79 | 0.51 | 0.43 | 0.56 | 0.65 |

Note:
\*p<0.1; \*\*p<0.01; \*\*\*p<0.001

The vast majority of effect estimates in both tables are discernible with *p* < .001. This is expected from variation in the evidential and evaluative thresholds (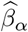 and 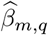), which correspond to dramatic changes in the density of the graph. The data source effects 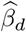 are likewise discernible, except in some cases with respect to degree distribution (*δ*, 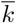, *σ*), indicating that none of the data sets produces in any sense an “average” comorbidity network. Some of the data sets produce similar networks, for example MedPAR(3) and Michigan, or Stanford and Columbia*. The latter use the same ontology and were processed in the same way, though the former have no such similarities. The MedPAR(3) and MedPAR(5) networks differ only in the resolution of their ontologies, and their global properties are broadly similar; but the Columbia and Columbia* networks share these background similarities (and the same ontology) yet exhibiting very different structure.

Each statistic varies widely (given the range of its possible values) across these comorbidity networks, though much of this variation can be explained by network density, which is largely a product of population or sample size. Increasing the number of links in a graph, whether by increasing *α* or by decreasing *θ_m_*, has the expected effect on LCP (more nodes in the largest component); on *δ*, 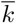, and *μ* (greater density); and on 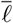 (shorter geodesics). It also tends to decrease the correlation between the total comorbidities of comorbid disorders (*r*) and the modular structure of the network (*Q*). Since these networks tend to be denser than other empirical networks to begin with, this suggests that graphs obtained via more relaxed cutoffs are saturated with connections in a way that obscures these hierarchical properties. Triad closure increases with stricter evidential cutoffs but decreases with stricter evaluative cutoffs. This suggests that cliques of three or more associated disorders often have strong evidential support while their pairwise associations vary widely in strength. This may be understood as some comorbid relations being artifacts of others, i.e. “transitive correlations” (Tao 2014).

Several patterns emerge from the PCA biplot (Figure 1): Judging from the variable loadings, PC1 captures a spectrum between highly connected and dense graphs with shorter internode distances, in which communities are difficult to detect, and sparser, more modular graphs. This spectrum aligns with the differences in density, encoded as the opacity of the plotting symbol, produced from a single dataset by varying the evaluative cutoff. PC2 discriminates between more homogeneous graphs, in terms of degree distribution (low scores), and those high in degree assortativity and triad closure (high scores). Graphs on finer ontologies, such as the level-5 ICD9 codes used by Hidalgo et al. (2009) and Hanauer and Ramakrishnan (2013), varied more widely across different choices of BAM; while graphs on coarser ontologies, in particular the level-3 ICD9 codes used Hidalgo et al. (2009) and the custom ICD9 mapping of Rzhetsky et al. (2007), varied less by BAM. Their values were more distant from the average than those of the graphs on coarser ontologies, which indicates that their stuctural properties were more idiosyncratic.

**Figure 1:**
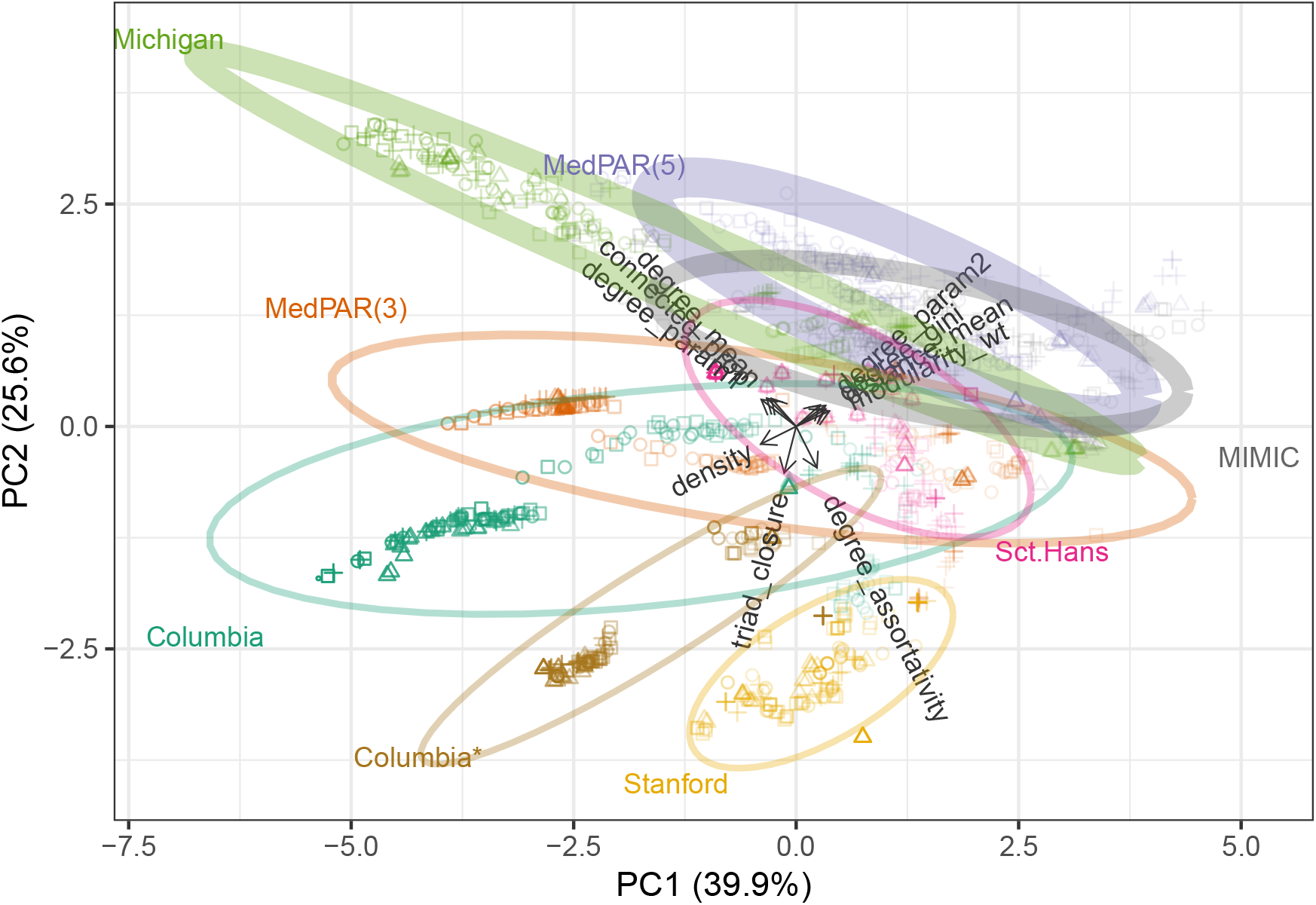
Row-principal PCA biplot for the summary statistics with networks (cases) in principal coordinates and statistics (variables) in standard coordinates. The values for graphs constructed from a common dataset are summarized by 95% confidence ellipses. Symbol corresponds to BAM, color indicates data source, and opacity is proportional to network density. Ellipse thicknesses are proportional to the number of clinical concepts (nodes) in the ontology (graph).

The networks constructed from a common dataset form subsets that emanate outward in clearly different directions, indicating that comorbidity network structure depends crucially on the source of data. These clusters have relatively consistent scores on PC2, which separates the clusters by ontology size. In contrast, they vary widely along PC1, which is in most cases clearly correlated with changes due to the evaluative cutoff. These observations are nearly exhaustive: together PC1 and PC2 account for more than 60% of the variance in the point cloud, with PC3 accounting for less than 15% more.^8^

#### 6.1.4 Centrality rankings of disorders

From correlation biplots, we observed great variation in the disorder rankings within each data source and centrality measure (Figure S9). Note that, for these comparisons, the p-value cutoff determined the discrete network structure while the BAM determined the weighting scheme upon this structure. While rankings were rarely discordant (the relative positions of more pairs of disorders reversed than preserved), frequently two different constructions yielded weak correlations (*r* < .5). The rankings were sensitive to both construction parameters, and no single choice of p-value correction or BAM consistently produced rankings that were robust to the other parameter.

The data sets also range widely in terms of which construction parameter explains more of the variation in rankings. For an example taken at random, rankings of full ICD9 codes based on Michigan data were sensitive to the BAM though robust to the correction (Figure 2). In other cases, rankings were variably more sensitive to the BAM (MedPAR(3), Columbia*) or to the correction (Sct. Hans, MIMIC). This is quantified in terms of the decomposition of inertia (Table 4). Comorbidity networks based on Sct. Hans and MIMIC produced centrality rankings that were highly sensitive to the p-value correction but robust to the BAM, while rankings of those based on MedPAR(3) and Columbia* were more sensitive to the BAM. The underlying ontologies do not explain these differences. However, the results reveal clear differences between the sensitivities of the rankings based on different centrality measures, with degree centrality most sensitive to the p-value correction and closeness centrality most sensitive to the BAM (though only slightly more than betweenness). This is consistent with the theoretical distinctions between the measures, with betweenness and closeness reliant on geodesic paths that vary more with differences in link weights, and may help guide the choice of measure best suited to a particular application.

**Figure 2:**
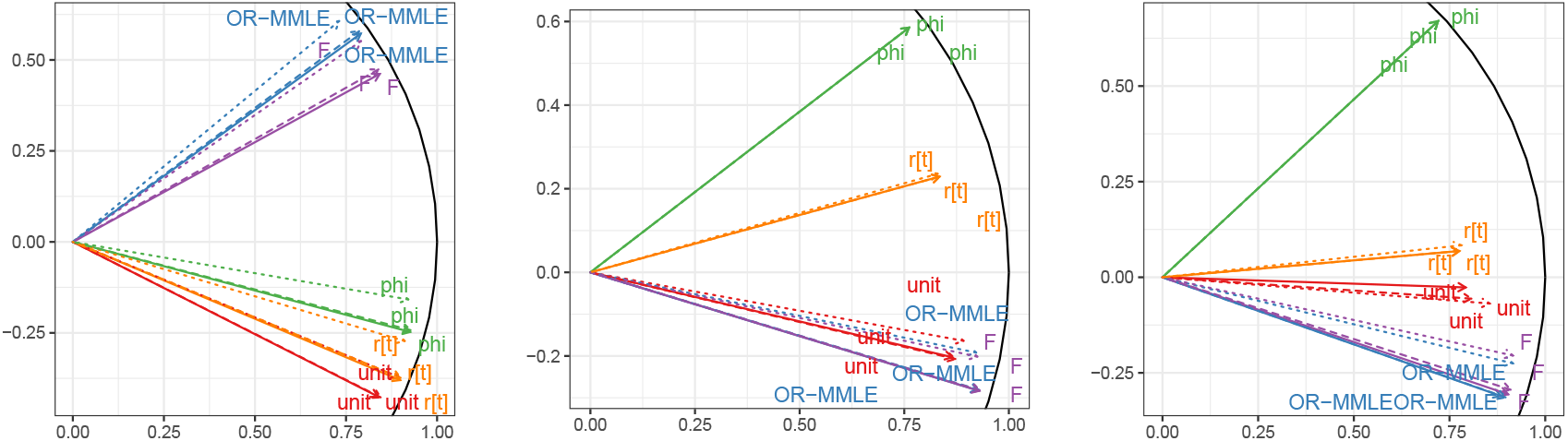
Eigendecomposition biplots for the Kendall correlations among (left to right) degree, betweenness, and closeness centrality rankings of disorders in networks constructed from the Michigan data, using a 5% TWER with each error rate correction and each BAM. The linetype of each arrow indicates the correction (solid for none, dotted for FWER, dashed for FDR) and its color and label indicate the BAM.

**Table 4:** Proportion of variance in rank correlations accounted for by the choice(s) of p-value correction, of binary association measure, and of both. Kendall rank correlations were calculated among all combinations of network construction parameters within each data source and centrality measure.

| Dataset | Ontology | Centrality | Correction | Measure | Both |
| --- | --- | --- | --- | --- | --- |
| Columbia | Rzhetsky | degree | 0.055 | 0.883 | 0.939 |
| Columbia | Rzhetsky | betweenness | 0.435 | 0.469 | 0.903 |
| Columbia | Rzhetsky | closeness | 0.141 | 0.772 | 0.913 |
| MedPAR(3) | ICD9-3 | degree | 0.082 | 0.819 | 0.901 |
| MedPAR(3) | ICD9-3 | betweenness | 0.198 | 0.700 | 0.899 |
| MedPAR(3) | ICD9-3 | closeness | 0.096 | 0.816 | 0.912 |
| MedPAR(5) | ICD9-5 | degree | 0.490 | 0.365 | 0.855 |
| MedPAR(5) | ICD9-5 | betweenness | 0.680 | 0.223 | 0.902 |
| MedPAR(5) | ICD9-5 | closeness | 0.532 | 0.311 | 0.843 |
| Sct.Hans | ICD10-3 | degree | 0.240 | 0.606 | 0.846 |
| Sct.Hans | ICD10-3 | betweenness | 0.870 | 0.092 | 0.962 |
| Sct.Hans | ICD10-3 | closeness | 0.877 | 0.090 | 0.967 |
| Michigan | ICD9-5 | degree | 0.009 | 0.978 | 0.987 |
| Michigan | ICD9-5 | betweenness | 0.010 | 0.978 | 0.988 |
| Michigan | ICD9-5 | closeness | 0.009 | 0.975 | 0.984 |
| Stanford | Rzhetsky | degree | 0.007 | 0.987 | 0.994 |
| Stanford | Rzhetsky | betweenness | 0.434 | 0.496 | 0.930 |
| Stanford | Rzhetsky | closeness | 0.031 | 0.911 | 0.941 |
| Columbia* | Rzhetsky | degree | 0.003 | 0.995 | 0.998 |
| Columbia* | Rzhetsky | betweenness | 0.225 | 0.726 | 0.951 |
| Columbia* | Rzhetsky | closeness | 0.024 | 0.954 | 0.978 |
| MIMIC | ICD9-5 | degree | 0.533 | 0.256 | 0.789 |
| MIMIC | ICD9-5 | betweenness | 0.795 | 0.094 | 0.889 |
| MIMIC | ICD9-5 | closeness | 0.713 | 0.110 | 0.824 |

The betweenness centrality scores, like those for degree but unlike those for closeness, yield clear hubs, dominated by non-specific symptoms and disorders. For the regional EHR data sets, these include epilepsy (Columbia), limb pain, unclassified respiratory problems (Michigan), vitamin deficiency (Stanford), benign neoplasms, and tuberculosis (Columbia*). While the rankings differ, most of the same disorders fill the top slots according to degree centrality. These include gram-negative bacteria, carcinoma in situ, lipid metabolism disorders (Columbia), unclassified respiratory disorders, unspecified chest pain, unspecified pneumonia (Michigan), acidosis, mineral metabolism of calcium and magnesium (Stanford), and Hepititis C (Columbia*).

The more specific populations covered by the other data sets yield their own characteristic hubs: non-specific diagnoses of fluid and electrolyte imbalances, urinary tract disorders, and bacterial infections (MedPAR), which may be associated with increased hospital and nursing home care as well as with aging itself; gait and mobility disorders, which are strongly associated with nervous disorders (Sct. Hans); and acute posthemorrhagic anemia (APHA), a common symptom of injury-induced blood loss (MIMIC). The high betweenness of these disorders can be largely explained by their high degree; they are associated with a variety of comorbidities each. Indeed, in the first two cases (Medicare and psychiatric patients), the same disorders are identified as hubs according to degree centrality. In the third case (intensive care patients), though, despite its more than two-fold lead in betweenness centrality, APHA has lower degree centrality than unspecified acute kidney failure, acute respiratory failure, severe sepsis, unspecified urinary tract infection, acidosis, and unspecified congestive heart failure. This discrepancy may reflect that APHA is a common result of several otherwise ontologically distinct types of injury and trauma.

**Table 5:**
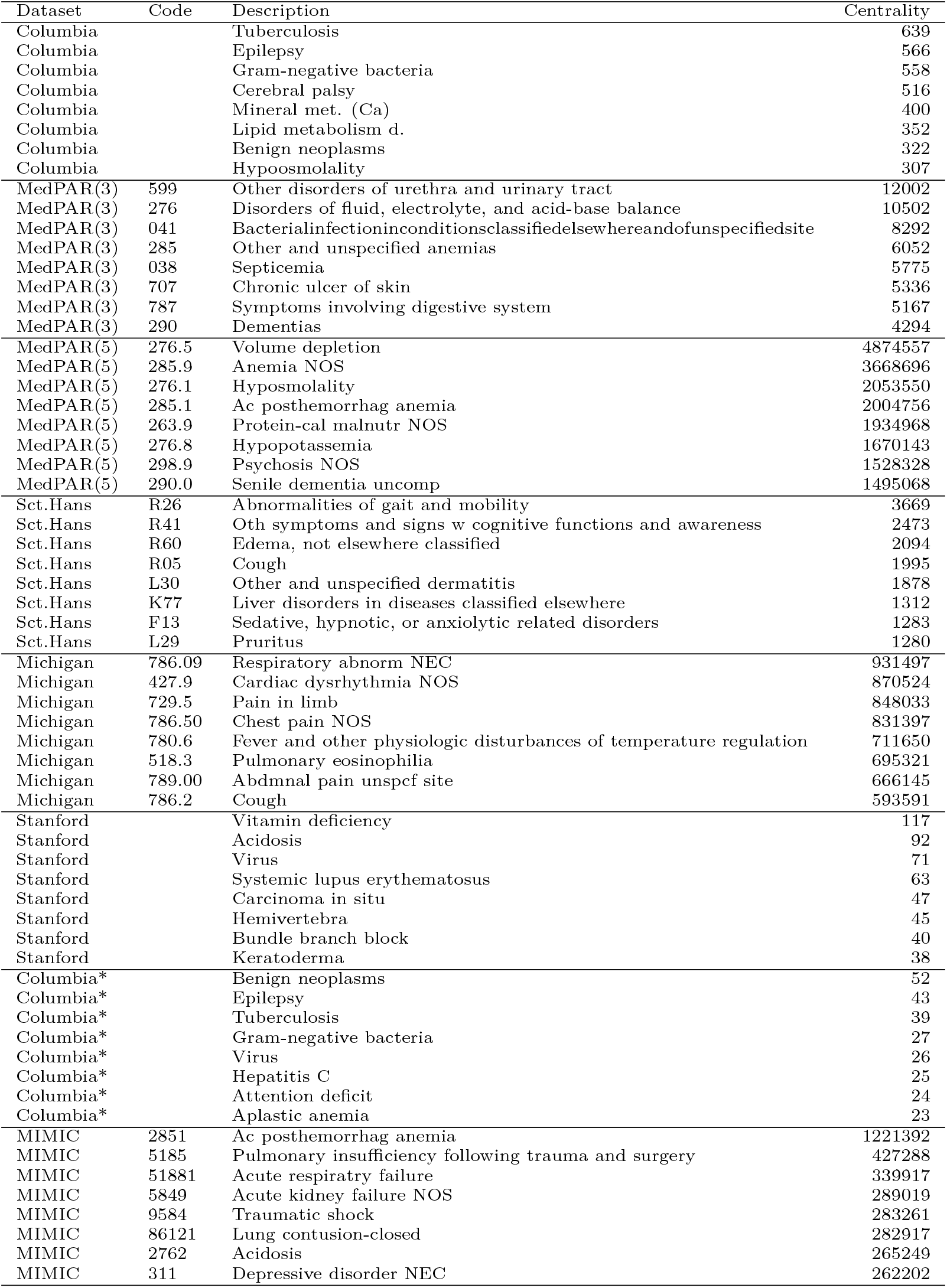
Most betweenness-central disorders in each comorbidity network, subject to Benjamini-Hochberg-corrected 5% FDR.

Notably, centrality rankings of the disorders in a common ontology, but based on node versus group centralities calculated in different network models, were generally not concordant: in some case the concordance was weak, in others negative. The limitations of our data and methodology prevented us from determining how much of this discordance was attributable to the data (i.e. the patient populations and collection protocols) versus to the effect of using group centralities versus crosswalking the incidence data before constructing the network. See the SI Text for more details.

### 6.2 Multiple and multivariate analysis

#### 6.2.1 Correlation structure and link determination for NAMCS encounters

The four correlation matrices *R_t_*, 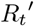, 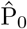, 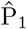 exhibited both clear similarities and clear differences (Figure S18). Most notably, in the pairwise and full partial correlation matrices *R_t_* and 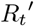, all 13 disorders loaded positively onto the first eigenvector, consistent with most comorbidity (controlling for prevalence) being explained by a one-dimensional spectrum between good and poor health. This spectrum was most closely aligned, under both models, with hypertension (HT), hyperlipidemia (HLD), and ischemic heart disease (IHD). The matrices differed starkly, though, in the proportion of variance captured by this first eigenvector—64% (pairwise) versus 22% (partial). The joint distribution estimates 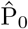 and 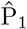 attributed only about 30–40% of variance to the first eigenvector, and this dimension was again most aligned with HT, HLD, and IHD. In contrast, though, these models more clearly oriented some disorders, particularly depression and cancer, in opposition to the majority of others, including diabetes and obesity in addition to the aforementioned cardiac disorders.

Figure 3 presents four networks on the node set of NAMCS chronic diseases, using the conventional, partial correlation, and two JDM constructions. Most of the residual interactions in JDM1 were weaker than in JDM0—for example, the associations of HT with arthritis and with cerebrovascular disease (CVD). From the conventional to the multivariate models, some pairs switched from positive to indiscernible (chronic obstructive pulmonary disease (COPD) with osteoporosis (OP), arthritis with IHD), negative to indiscernible (IHD with obesity), or indiscernible to negative (CVD with obesity). Notably, all three negative associations observed in the pairwise analysis were negative in every analysis, and every positive association observed under the strictest model (JDM1) was positive under the pairwise model. See the SI Text for a quantitative comparison of the correlation estimates from each of the models.

**Figure 3:**
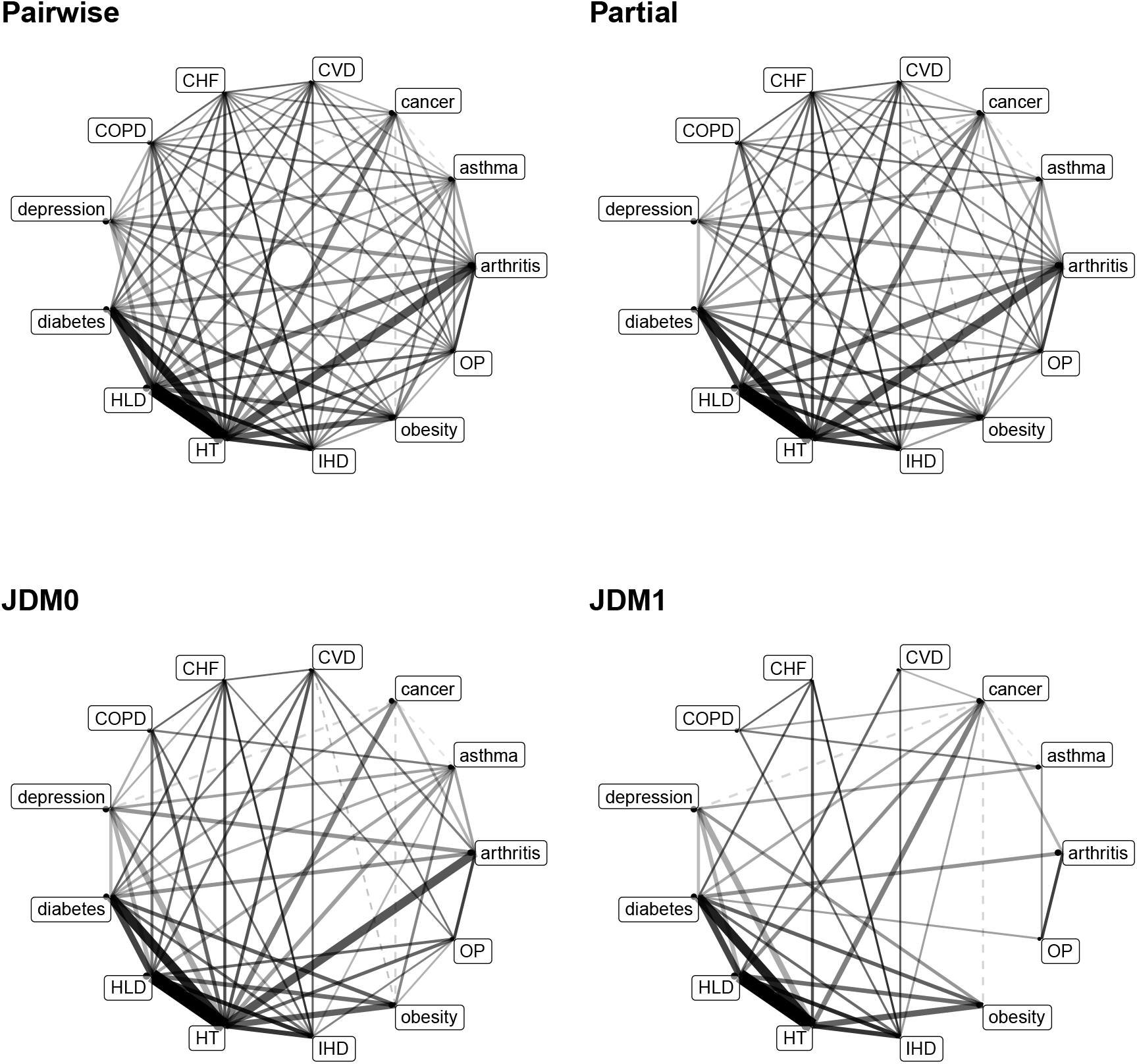
Four comorbidity networks constructed from the NAMCS chronic disease incidence data. From left to right, then top to bottom: conventional comorbidity network with links determined from a 5partial correlation comorbidity network adapted from the conventional network; JDM network controlling only for disease prevalence, with links weighted by 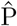; JDM network also controlling for patient-level demographics. Black (respectively, grey) links indicate positive (negative) associations.

#### 6.2.2 Correlation structure and link determination for MIMIC-III units

Results for the care unit populations from MIMIC-III were in several respects dissimilar to those for NAMCS. Network diagrams showed similar behavior across the units but were less informative due to the number of nodes in each; they are included for each unit and each model as supporting figures.

Table 6 reports, for each care unit and each network model, the proportions of positive, negative, and indiscernible links, based on whether the confidence (resp. credible) interval of radius two standard errors (resp. standard deviations) about each correlation estimate contains zero. The clear patterns across most units are that (a) in the pairwise network, vastly more links are positive than negative, and more negative than missing; (b) in the partial correlation network, links are positive, negative, and missing in roughly equal proportions; and (c) in the joint distribution network, vastly more links are missing than positive, and more positive than negative. For the neonatal units (NWARD and NICU), which used smaller ontologies, the majority types (the positive links in the pairwise network and the missing links in the joint distribution network) were still more dominant, and much more of the links in the pairwise correlation network were missing.

**Table 6:** Link determinations for each MIMIC-III care unit using each model, with a consistent evidential threshold of 2 standard deviations.

| Model | Determination | NWARD | MICU | SICU | NICU | CCU | CSRU | TSICU |
| --- | --- | --- | --- | --- | --- | --- | --- | --- |
| Pairwise | Positive | 90.9% | 60.6% | 66.5% | 91.3% | 73.9% | 77.2% | 69.0% |
| Pairwise | Negative | 6.8% | 31.2% | 24.1% | 6.6% | 16.9% | 15.7% | 22.1% |
| Pairwise | Indiscernible | 2.3% | 8.2% | 9.4% | 2.1% | 9.2% | 7.1% | 9.0% |
| Partial | Positive | 17.2% | 39.6% | 35.2% | 26.2% | 36.4% | 37.7% | 33.2% |
| Partial | Negative | 10.0% | 36.2% | 32.2% | 20.3% | 33.4% | 34.1% | 30.2% |
| Partial | Indiscernible | 72.8% | 24.2% | 32.6% | 53.6% | 30.2% | 28.2% | 36.6% |
| JDM | Positive | 0.4% | 40.0% | 20.8% | 0.8% | 13.6% | 15.4% | 15.6% |
| JDM | Negative | 0.4% | 0.0% | 0.0% | 0.4% | 0.2% | 0.4% | 0.0% |
| JDM | Indiscernible | 99.2% | 60.0% | 79.2% | 98.8% | 86.2% | 84.3% | 84.4% |

Scatterplots of correlation estimates from the different models are included as supporting figures. In contrast to NAMCS, in this case the partial estimates were more correlated than the JDM estimates with the pairwise estimates. Though the correlation was strong between estimates in the conventional model and the JDM among the most prevalent disorders, those for the least prevalent were essentially uncorrelated. Meanwhile, the relationship between the pairwise and partial correlations was roughly a scaling one: For this and other non-neonatal units, the relationship was noisy but compatible with the constraint of passing through the origin.

#### 6.2.3 Centrality rankings of disorders

As in our comparisons of pairwise constructions, centrality rankings of disorders within each MIMIC unit varied greatly by network model, and overall patterns were not evident (Figure S29). Pairwise network rankings consistently stood apart from those of the multivariate model–based networks based on the medical intensive care unit (MICU) data; joint distribution network rankings tended to be more distinctive for the neonatal ward (NWARD), full partial network rankings were more distinctive for the neonatal intensive care unit (NICU), and for other units no two models consistently produced similar rankings.

## 7 Acknowledgments

Beverly Setzer and Lauren Geiser contributed code for processing data and constructing graphs as well as helpful methodological conversations. David Hanauer kindly shared the pre-processed contingency table data from the University of Michigan Health System. JCB was supported in part by an NIH T90 training grant (5T90DE021989-07).

## 8 Supporting tables and figures

Most supporting figures and one supporting table are made available as separate files. The remainder appear in the SI Text. LATEXlabels correspond to file paths.

Table S1: Quintiles of binary association measures for pairs in each dataset for different evidential thresholds. “B” indicates Bonferroni correction.

Table S2: Quintiles of binary association measures for pairs in each dataset for different evidential thresholds. “B” indicates Bonferroni correction.

Figure S1: For each data source (row), evaluative cutoff (column), and BAM (abscissa), a box-and-whisker plot of the quantiles at which the links of each comorbidity network were trimmed by the cutoff. The networks were constructed over the range of evidential (p-value) cutoffs 10^−*i*^, *i* = 1,…, 6 and for each p-value correction (none, FWER, FDR).

Figure S2: Network diagram (hairball plot) of the comorbidity network constructed from the Columbia data, using the evidential cutoff *α* < 5% with Bonferroni correction and the evaluative cutoff *OR* ≥ 6. Clusters identified using the Walktrap algorithm are color-coded.

Figure S3: Network diagram (hairball plot) of the comorbidity network constructed from the MedPAR data on level-3 ICD9 codes, using the evidential cutoff *α* < 5% with Bonferroni correction and the evaluative cutoff *OR* ≥ 6. Clusters identified using the Walktrap algorithm are color-coded.

Figure S4: Network diagram (hairball plot) of the comorbidity network constructed from the Sct. Hans data, using the evidential cutoff *α* < 5% with Bonferroni correction and the evaluative cutoff *OR* ≥ 6. Clusters identified using the Walktrap algorithm are color-coded.

Figure S5: Network diagram (hairball plot) of the comorbidity network constructed from the Stanford data, using the evidential cutoff *α* < 5% with Bonferroni correction and the evaluative cutoff *OR* ≥ 6. Clusters identified using the Walktrap algorithm are color-coded.

Figure S6: Network diagram (hairball plot) of the comorbidity network constructed from the Columbia data, using the evidential cutoff *α* < 5% with Bonferroni correction and the evaluative cutoff *OR* ≥ 6. Clusters identified using the Walktrap algorithm are color-coded.

Table S3: Log-likelihood ratios *R* and two-sided p-values *p* from likelihood-ratio tests between four families of models to degree sequence tails. LRTs were performed for graphs constructed from each dataset using two p-value significance thresholds and corrections for family-wise error rate and for false discovery rate. Family 1 is preferred when *R* > 0, Family 2 when *R* < 0.

Figure S7: Diverging-color tilings of log-likelihood ratios *R* from likelihood-ratio tests (LRT) between fits of different families of models to degree sequence tails. LRTs were performed for graphs constructed from each dataset using evidential cutoff *α* < 0.05 and both corrections. Family 1 is preferred when *R* > 0, Family 2 when *R* < 0. The boundary color of each tile indicates whether *R* is positive or negative. The p-value from the LRT is printed on each tile.

Table S4: Tail index (*ξ*) and tail exponent (*γ*) estimators that were successfully estimated on comorbidity networks.

Table S5: Values of several global statistics calculated on comorbidity networks constructed over a range of data sets and parameter settings. See Section 5.1.4 in the main text.

Figure S8: Mean values of several global statistics calculated on comorbidity networks constructed over a range of data sets and parameter settings, plotted against several measures of network size. See Section 5.1.4 in the main text.

Table S6: LMs of network statistics on data source and test-wise error rate.

Table S7: Hierarchical models of network statistics on test-wise error rate, grouped by data source.

Table S8: Hierarchical models of network statistics on test-wise error rate and binary association measure, grouped by data source.

Table S9: Variance decomposition for hierarchical models of network statistics on test-wise error rate, grouped by data source.

Table S10: Variance decomposition for hierarchical models of network statistics on test-wise error rate and binary association measure, grouped by data source.

Table S11: Akaike information criteria for each model fitted to the values taken by each network statistic.

Figure S9: For each data source and centrality measure, a correlation biplot of Kendall correlations between centrality rankings of disorders based on comorbidity networks constructed using the evidential cutoff *α* < 0.05, each of three corrections (none, FWER, FDR), and each of five BAMs (unit, odds ratio, Pearson correlation, Forbes coefficient, and tetrachoric). In this and other biplots, first and second eigenvectors are reversed if necessary so that each centroid lies in the first quadrant.

Table S12: Most degree-central disorders in each comorbidity network, subject to Benjamini-Hochberg-corrected 5% FDR.

Table S13: Most closeness-central disorders in each comorbidity network, subject to Benjamini-Hochberg-corrected 5% FDR.

Table S14: Most prevalent disorders in each comorbidity network.

Figure S10: Scatterplots of prevalences of disorders in the ICD9 ontology in different data sets using this ontology. Note the effect of the restriction, in the Michigan dataset, to disorders that appeared on at least 30 patient records.

Figure S11: Scatterplots of prevalences of disorders in the Rzhetsky ontology in different data sets using this ontology. Note that the disorders included in the Columbia dataset exactly match their prevalences in the Columbia dataset.

Figure S12: For each p-value correction and centrality measure, a correlation biplot of Kendall correlations between centrality rankings of disorders based on comorbidity networks constructed from each data set that uses or could be crosswalked to the Rzhetsky ontology. Centrality measures are node-based for data encoded using this ontology and group-based for data crosswalked to this ontology.

Figure S13: For each p-value correction and centrality measure, a correlation biplot of Kendall correlations between centrality rankings of disorders based on comorbidity networks constructed from each data set that uses or could be crosswalked to the level-3 ICD9 ontology. Centrality measures are node-based for data encoded using this ontology and group-based for data crosswalked to this ontology.

Figure S14: For each p-value correction and centrality measure, a correlation biplot of Kendall correlations between centrality rankings of disorders based on comorbidity networks constructed from each data set that uses the level-5 ICD9 ontology.

Table S15: Most degree-central disorders in comorbidity networks using the ICD9-3 ontology, subject to Benjamini-Hochberg-corrected 5% FDR.

Table S16: Most degree-central disorders in comorbidity networks using the Rzhetsky ontology, subject to Benjamini-Hochberg-corrected 5% FDR.

Table S17: Most closeness-central disorders in comorbidity networks using the ICD9-3 ontology, subject to Benjamini-Hochberg-corrected 5% FDR.

Table S18: Most closeness-central disorders in comorbidity networks using the Rzhetsky ontology, subject to Benjamini-Hochberg-corrected 5% FDR.

Figure S15: Frequency–rank plot for the 13 chronic disorders recorded in the NAMCS sample.

Figure S16: Alluvial diagram of discernible signs of the population-level associations between NAMCS chronic disorders, using each of 4 network models: pairwise correlation, full partial correlation, endogenous joint distribution, and joint distribution controlling for exogenous predictors.

Table S19: Estimated effects of demographic predictors on the incidence of chronic disorders in Model 1. Each value indicates the effect of the predictor on the mean of the normal distribution from which the latent variable is sampled (see the text). Estimates whose 95% credible intervals contain zero are excluded.

Table S20: Histograms of estimates from the joint distribution model with exogenous (patient-level) predictors. Left: Exogenous effects on disorder prevalence. Right: Correlation (epidemiological comorbidity) accounted for by exogenous effects.

Figure S17: Alluvial diagram of discernible signs of the population-level associations between NAMCS chronic disorders, using each of 4 network models: pairwise correlation, full partial correlation, endogenous joint distribution, and joint distribution controlling for exogenous predictors.

Figure S18: For each of four models, a correlation biplot of estimated latent correlations between 13 chronic disorders in the NAMCS sample.

Figure S19: For each critical care unit in the MIMIC-III database, a frequency–rank plot for the recorded diagnoses, crosswalked to CCS codes.

Figure S20: Scatterplots between the pairwise tetrachoric correlations *r_t_*, the full partial correlations 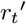, and the correlation estimates 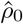 from the endogenous JDM for the CSRU population.

Figure S21: Alluvial diagrams of discernible signs of the population-level associations between MIMIC diagnoses within each admission unit cohort, using each of 3 network models: pairwise correlation, full partial correlation, and endogenous joint distribution.

Figure S22: Three comorbidity networks constructed from the CCU population. From left to right: Sample tetrachoric correlations *r_t_*, partial tetrachoric correlations 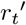, and correlation estimates 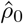 from the endogenous JDM.

Figure S23: Three comorbidity networks constructed from the CSRU population. From left to right: Sample tetrachoric correlations *r_t_*, partial tetrachoric correlations 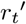, and correlation estimates 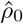 from the endogenous JDM.

Figure S24: Three comorbidity networks constructed from the MICU population. From left to right: Sample tetrachoric correlations *r_t_*, partial tetrachoric correlations 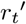, and correlation estimates 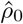 from the endogenous JDM.

Figure S25: Three comorbidity networks constructed from the NICU population. From left to right: Sample tetrachoric correlations *r_t_*, partial tetrachoric correlations 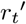, and correlation estimates 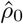 from the endogenous JDM.

Figure S26: Three comorbidity networks constructed from the NWARD population. From left to right: Sample tetrachoric correlations *r_t_*, partial tetrachoric correlations 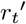, and correlation estimates 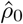 from the endogenous JDM.

Figure S27: Three comorbidity networks constructed from the SICU population. From left to right: Sample tetrachoric correlations *r_t_*, partial tetrachoric correlations 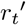, and correlation estimates 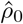 from the endogenous JDM.

Figure S28: Three comorbidity networks constructed from the TSICU population. From left to right: Sample tetrachoric correlations *r_t_*, partial tetrachoric correlations 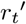, and correlation estimates 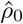 from the endogenous JDM.

Figure S29: For each care unit and centrality measure, a correlation biplot of Kendall correlations between centrality rankings of disorders based on each of four comorbidity network models: pairwise correlations, full partial correlations, and a joint distribution model without exogenous effects.

## Footnotes

1 These concerns apply to the analysis of disease co-occurrence data, not to their collection, with which a different body of work continues to raise and address concerns.

2 Construct a comorbidity network using the Rzhetsky data and ontology with a Bonferroni-corrected .05 evidential cutoff and weighting the links by the Forbes coefficient *F* (and restricting to positive links).

3 To avoid infinities, replace each weight *F* with 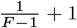.

4 The *χ*^2^ test commonly used in CNAs is an approximation to Fisher’s exact test.

5 The threshold ranges of *θ_m_* for each BAM *m* were chosen so that the corresponding quantiles of pairs in each data set are roughly equal (Figure S1). It should be noted, though, that the Pearson correlation thresholds are more restrictive than those of the other correlation measure *r_t_*.

6 We also fit two families of hierarchical regression models to the same data, grouping by data source and by BAM (Gelman et al. 2012). In terms of the Akaike information criterion (Burnham and Anderson 2004), these models were consistently worse, with one exception, than the corresponding multiple regression models, so we do not report the results here.

7 While a matrix of rank correlations may not be positive-semidefinite, it still admits an eigendecomposition. The correlations between the rankings will equal the cosines between the unit vectors associated with the rankings in the space of the eigenvectors. In most practical settings the first two eigenvalues will be positive, so correlations will be well-represented visually by cosines between the vectors projected to a two-dimensional biplot. So it was in our cases.

8 We repeated the PCA after including the comorbidity network constructed from VAERS. This network does not stand out from the rest, and indeed situates itself among those other networks constructed from larger data sets.

