## Supplemental Text for "Sensitivity and robustness of comorbidity network analysis"

### Supporting information for the paper: Sensitivity and robustness of comorbidity network analysis

#### Methods

##### Detailed discussion of the pairwise models

###### Data sets

As noted in the main text, the data sets from which we obtained pairwise contingency tables vary widely:

- Rzhetsky et al. (2007) used a coarse custom ontology mapped from level-5 ICD9 codes.[1] Their pairwise data are unique among this collection in including contingency tables for *every* pair of disorders.
- The MedPAR data used by Hidalgo et al. (2009) concern the specific subpopulation of Medicare patients over age 65, and were made available both according to full ICD9 and to the level-3 prefixes of these codes. Both data sets are limited to a subset of pairs, though the authors do not indicate how this subsetting was done.
- Sct. Hans is a psychiatric hospital, so this dataset concerns that specific subpopulation. Roque et al. (2011) restricted their attention (and their public dataset[2]) to 802 pairs of codes with Benjamini–Hochberg p-values $p < .01$ and odds ratios $OR > 2$.
- Hanauer and Ramakrishnan (2013) restricted the study dataset to pairs of codes that appeared in at least 30 patient records each and in at least 10 records together. The study focused on temporal relationships between diagnoses but the dataset was not trimmed to exclude pairs that were not temporally related (c.f. Jensen et al. (2014)).
- Bagley et al. (2016) adopted the custom ontology of Rzhetsky et al. (2007). They collected records in 2013 and excluded records before 2008, internally inconsistent records, patients older than 90, conditions with prevalence below 50, and pairs of conditions with co-occurrence below 5. They also included the Columbia data from Rzhetsky et al. (2007), re-processed in this way, which we designate with an asterisk (e.g. "Columbia*" in the densities table in the main text.
- The data in MIMIC-III is pooled only from hospital admissions that included a stay at any of five intensive care units; like the others mentioned, this subpopulation may exhibit peculiar traits.

[1] See SI (http://www.pnas.org/content/suppl/2007/06/21/0704820104.DC1), Appendix 3.
[2] See SI (http://journals.plos.org/ploscompbiol/article?id=10.1371/journal.pcbi.1002141#s5), Dataset S1.

In most cases, $\chi^2$ tests on the contingency tables yielded p-values ranging from 0 to $1-\varepsilon$, $\varepsilon < 10^{-3}$. The exception was the data from Sct. Hans, which included only highly discernible comorbid pairs ($p < 10^{-4}$).

##### Binary association measures

We used four binary association measures (BAMs) to evaluate the strength of association between disorders in the pairwise models:

- The *odds ratio* $OR = \frac{a/b}{c/d}$ is widely used in clinical research for its interpretability, comes recommended as a standard measure of epidemiological comorbidity (Kraemer 1995), and was used in some comorbidity network studies (Hanauer et al. 2009; Hanauer and Ramakrishnan 2013; Kim et al. 2016). When the binary variables are independent, it has expected value 1. In most analyses, in order to avoid infinite link weights, we substitute the modified maximum likelihood estimate of the odds ratio, $\widehat{OR} = \frac{(a+.5)(d+.5)}{(b+.5)(c+.5)}$ (Parzen et al. 2002).
- *Pearson's binary correlation coefficient* $\phi = \frac{ad-bc}{\sqrt{(a+b)(a+c)(b+d)(c+d)}}$, designated $A_{30}$ (and misidentified as the tetrachoric) in Hubalek's (1982) review of BAMs, was used in several exploratory comorbidity network studies (Hidalgo et al. 2009; Folino and Pizzuti 2012; Chen and Xu 2014; Chmiel et al. 2014; Lai 2015). Being a correlation coefficient, its values range from $-1$ to 1, with independent variables having expected value 0.
- *Forbes' coefficient of association* $F = \frac{a/(a+b)}{(a+c)/(a+b+c+d)}$, designated $A_{40}$ by Hubalek (1982), was often used together with $\phi$ and sometimes called "relative risk" (Hidalgo et al. 2009; Chen and Xu 2014; Chmiel et al. 2014)—though it is distinct from that term's two usual referents, the risk ratio formula and $OR$ (Zhang and Yu 1998). Like $OR$, $F$ has expected value 1 when the variables are independent.

We additionally use the *tetrachoric correlation coefficient* $r_t$. To motivate this statistic, consider that one can generate a $2 \times 2$ contingency table by sampling ordered pairs $(Z_1, Z_2) \sim N(\boldsymbol{\mu}, \Sigma)$ from a bivariate standard normal distribution with means $\mu = (\mu_1, \mu_2)^\mathsf{T}$ and covariance matrix

$$\Sigma = \begin{pmatrix} {\sigma_1}^2 & \rho\sigma_1\sigma_2 \\ \rho\sigma_1\sigma_2 & {\sigma_2}^2 \end{pmatrix},$$

and then dichotomize the samples $(z_1, z_2)^\mathsf{T}$ according to sign:

$$y_i = \begin{cases} 0 & z_i \leq 0 \\ 1 & z_i > 0 \end{cases}$$

The marginal probabilities $P(D_i)$ for the table are then determined by the scaled means $\mu_1/\sigma_1$ and $\mu_2/\sigma_2$, while the joint probabilities deviate from the products of the marginals according to the linear correlation $\rho$ between the latent variables. Under this model, $r_t$ is the value of $\rho$ for which the joint probabilities are in proportion to the empirical data (Drasgow 2006). While disease diagnoses are discrete events, the underlying normal distributions can be interpreted as distributions of health and risk, so that a diagnosis occurs when the value exceeds a certain threshold.

#### Detailed discussion of the multivariate models

##### Partial correlation networks

The statistical tests conventionally used to identify comorbidities fail to account for an important source of variation actually contained within the underlying incidence data: associations with disorders beyond the index pair. One well-known mechanism is common causation, when, for instance, two disorders are clinically unrelated but have a common risk factor that results in a strong association between them in the population. Comorbidity network researchers are aware that such "transitive" correlations are likely and have suggested that clustering patterns should be examined with such explanations in mind (Hanauer et al. 2009); though such confounding can also result in real, possibly causal comorbidities being obscured.

Partial correlations provide a way to control for these confounding effects from the correlation matrix calculated from pairwise data. In the case of normally-distributed data $\mathbf{y} \sim N(\mathbf{0}, \Sigma)$ with $\operatorname{diag} \Sigma = ({\sigma_1}^2, \ldots, {\sigma_n}^2)^\intercal$, the *partial correlations*

$$r(y_i, y_j \mid \mathbf{y} \setminus \{y_i, y_j\}) = -\frac{\kappa_{ij}}{\sqrt{\kappa_{ii}\kappa_{jj}}}$$

between $y_i$ and $y_j$, obtained using the precision matrix $\Sigma^{-1} = \mathrm{K} = (\kappa_{ij})$, are the standardized coefficients $\beta_{ij}\sigma_j/\sigma_i = \beta_{ji}\sigma_i/\sigma_j$ obtained by maximum-likelihood estimation for multiple regression models predicting $y_i$ from $\mathbf{y} \setminus \{y_i\}$ and vice-versa. Epskamp and Fried (2017) recommend modeling continuous multivariate data as a network with links weighted by their partial correlations. When multivariate binary data are modeled using latent multivariate normal distributions, this technique can be applied to pairwise tetrachoric correlations, yielding *partial tetrachoric correlations* ${r_t}'$. We take this approach to our data sets to construct comorbidity networks based on partial latent correlations.

The multiple regression framework controls for confounding of the effect on $y_i$ of $y_j$ by treating the remaining $y_k$ as independent covariates. In reality, the correlation between two disorders may depend on interaction effects between other disorders as well as on their raw incidence. Thus the partial correlation network is only a first-order correction to confounding among the response variables. We next consider a multivariate model that encodes the complete set of interactions as a covariance matrix and takes a Bayesian approach to estimate them simultaneously.

##### Joint distribution networks

In addition to endogenous confounding by other disorders, spurious effects can also arise when exogenous factors, i.e. patient-level covariates, are ignored: If specific demographic subgroups (e.g. elderly or infirm patients) are at heightened risk of many otherwise physiologically unrelated disorders, failure to stratify the analysis by these demographics will result in artificially inflated correlations among these disorders. Several comorbidity network studies incorporated demographic predictors into their binary association tests (Rzhetsky et al. 2007; Feldman et al. 2016; Glicksberg et al. 2016), though this information was unavailable for our sensitivity analysis.

Comorbidity network researchers have also acknowledged the problem of multiple comparisons, which becomes acute in this setting due to the combinatorial explosion in the number of subsets of an already large number of variables. Even the coarse ontology of Rzhetsky et al. (2007) presents 12,880 pairings among its 161 concepts, which would require a $p = 4 \times 10^{-6}$ threshold on each pairwise test to obtain a family-wise error rate of .05. Comorbidity network researchers have differed between adopting a classical (Hanauer and Ramakrishnan 2013) versus a corrected cutoff (Bagley et al. 2016). Even the corrections considered in the previous phase do not resolve the problem of dependency: the tests rely on the same underlying population and the same response variables, as do the partial correlation networks described previously.

The multiple comparisons problem can be obviated by adopting a hierarchical modeling framework, in which grouping effects are treated not as independent but as sampled from some higher-order distribution with its own central tendency and spread (Gelman et al. 2012). Variation between groups is then partially attributed to this spread, so that estimated group averages "shrink" toward the population average. Meanwhile, the appropriate framework for modeling multiple potentially correlated (or anti-correlated) response variables is multivariate regression, which encodes the interactions between the responses as a covariance matrix $\Sigma$. Once such a model is fitted to incidence data, a network model can be inferred from the estimated covariance matrix $\hat{\Sigma}$ by normalizing $\hat{\Sigma}$ to an estimated correlation matrix $\hat{\mathrm{P}}$ (Rho), dropping statistically weak entries, and interpreting the result as a weighted adjacency matrix. In addition to incorporating endogenous effects among these responses, this framework admits exogenous predictors in the same manner as classical regression.

We adapted for these purposes the *joint distribution model* (JDM) proposed by Pollock et al. (2014) to allow ecologists to incorporate both species–environment relationships (exogenous effects) and species–species interactions (endogenous effects). The model includes a matrix $B$ of exogenous effects interpretable in the traditional manner, a matrix $\mathbf{P}$ of contributions from the exogenous factors to the observed co-occurrence rates, and a matrix $\mathrm{P}$ of latent correlations interpretable as comorbidities in the same way as tetrachoric correlations. Indeed, this model generalizes the latent bivariate normal model underlying the tetrachoric correlation coefficient, so that the off-diagonal entries of the residual correlation matrix $\hat{\mathrm{P}}$ can be directly compared with the $r_t$.

### Results

#### Additional results from the pairwise analysis

##### Link determination

In addition to network densities, we also calculated the quintiles of each BAM for each dataset, restricted to associations remaining at each TWER, in order to compare the distributions of the BAMs over the comorbid associations. The quintiles are reported in Tables S1 and S2. The BAM distributions are not highly sensitive to the TWER, and the direction of change (whether quintiles increase or decrease as the TWER is tightened) is inconsistent from dataset to dataset. This indicates that, in general, the amount of evidence for a comorbid association is not predictive of its strength. Some network properties may therefore be more sensitive to an evaluative cutoff than to an evidential cutoff. Once a fixed TWER has been imposed, BAM quintiles typically differ by no more than a factor of 10.

Figure S1: For each data source (row), evaluative cutoff (column), and BAM (abscissa), a box-and-whisker plot of the quantiles at which the links of each comorbidity network were trimmed by the cutoff. The networks were constructed over the range of evidential (p-value) cutoffs $10^{-i},\ i = 1, \ldots, 6$ and for each p-value correction (none, FWER, FDR).

Figure S2: Network diagram (hairball plot) of the comorbidity network constructed from the Columbia data, using the evidential cutoff $\alpha < 5\%$ with Bonferroni correction and the evaluative cutoff $OR \geq 6$. Clusters identified using the Walktrap algorithm are color-coded.

Figure S3: Network diagram (hairball plot) of the comorbidity network constructed from the MedPAR data on level-3 ICD9 codes, using the evidential cutoff $\alpha < 5\%$ with Bonferroni correction and the evaluative cutoff $OR \geq 6$. Clusters identified using the Walktrap algorithm are color-coded.

Figure S4: Network diagram (hairball plot) of the comorbidity network constructed from the Sct. Hans data, using the evidential cutoff $\alpha < 5\%$ with Bonferroni correction and the evaluative cutoff $OR \geq 6$. Clusters identified using the Walktrap algorithm are color-coded.

Figure S5: Network diagram (hairball plot) of the comorbidity network constructed from the Stanford data, using the evidential cutoff $\alpha < 5\%$ with Bonferroni correction and the evaluative cutoff $OR \geq 6$. Clusters identified using the Walktrap algorithm are color-coded.

Figure S6: Network diagram (hairball plot) of the comorbidity network constructed from the Columbia data, using the evidential cutoff $\alpha < 5\%$ with Bonferroni correction and the evaluative cutoff $OR \geq 6$. Clusters identified using the Walktrap algorithm are color-coded.

**Degree sequence distributions**

Table S3 reports the results of the LRTs performed between different families of distributions on the tails of the comorbidity network degree sequences. (A diverging-color tiling visualization is provided in Figure S7.) In general, the distributions are preferred in the following order:

$$\text{log-normal} \ > \ \text{exponential} \ > \ \text{power law} \ > \ \text{Poisson}$$

Comorbidity networks have veritably heavy tails that are poorly modeled by Poisson distributions. Only in one case (MedPAR(3)) was the power-law fit clearly preferred over the exponential fit, and in no cases was it preferred over the log-normal fit. As has been observed of many other real-world networks using this approach (Broido and Clauset 2018), comorbidity networks tend to be better-modeled by log-normal distributions.

There is no consistent relationship, controlling for the dataset, between the choice of evidential threshold and the strength of evidence for one family over another. For example, the Bonferroni correction to the MedPAR(3) network construction reversed the observations that node degrees were better modeled by a power law or a log-normal than an exponential, whereas the conclusions for the MedPAR(5) network essentially did not change. In general, though, tightening the evidential threshold had the effect of weakening the preferred order described in the previous paragraph.

Table S4 reports the estimators obtained under the hypothesis of regular variation. The method often failed to produce estimators; in cases where more than one estimator was calculated, usually at least one was negative (hence uninterpretable). In the 14 cases (out of 48 attempts) in which at least two estimators were positive, which were based on degree sequences obtained for MedPAR(5), Michigan, and MIMIC, they took dissimilar values and were fitted to the degree sequence tail at different scales. Moreover, there was no evident relationship between the estimators obtained using the three implemented methods (Adjusted Hill, moments, and kernel-type) across the networks.

Table S1: Quintiles of binary association measures for pairs in each dataset for different evidential thresholds. "B" indicates Bonferroni correction.

| TWER | BAM | quant. | Columbia | MedPAR(3) | MedPAR(5) | SctHans | Michigan | Stanford | Columbia* | MIMIC |
|---|---|---|---|---|---|---|---|---|---|---|
| 1 | $\widehat{OR}$ | 0% | 1.002 | 1.000 | 1.000 | 2.530 | 1.000 | 1.160 | 1.209 | 1.000 |
| 1 | $\widehat{OR}$ | 20% | 2.377 | 1.274 | 1.568 | 3.414 | 2.470 | 2.404 | 2.509 | 2.462 |
| 1 | $\widehat{OR}$ | 40% | 4.539 | 1.621 | 2.450 | 4.258 | 3.784 | 4.141 | 3.651 | 4.673 |
| 1 | $\widehat{OR}$ | 60% | 11.215 | 2.227 | 4.530 | 5.962 | 5.673 | 7.308 | 5.038 | 10.181 |
| 1 | $\widehat{OR}$ | 80% | 52.045 | 4.065 | 13.359 | 13.238 | 10.424 | 11.895 | 9.393 | 34.174 |
| .05 | $\widehat{OR}$ | 0% | 1.089 | 1.008 | 1.011 | 2.530 | 1.028 | 1.160 | 1.209 | 1.098 |
| .05 | $\widehat{OR}$ | 20% | 2.513 | 1.450 | 2.239 | 3.414 | 2.800 | 2.404 | 2.509 | 4.540 |
| .05 | $\widehat{OR}$ | 40% | 3.675 | 1.845 | 4.172 | 4.258 | 4.089 | 4.141 | 3.651 | 11.734 |
| .05 | $\widehat{OR}$ | 60% | 5.628 | 2.552 | 11.801 | 5.962 | 6.030 | 7.308 | 5.038 | 44.221 |
| .05 | $\widehat{OR}$ | 80% | 12.350 | 4.938 | 63.885 | 13.238 | 11.088 | 11.895 | 9.393 | 132.922 |
| .001 | $\widehat{OR}$ | 0% | 1.135 | 1.008 | 1.012 | 2.530 | 1.070 | 1.160 | 1.209 | 1.132 |
| .001 | $\widehat{OR}$ | 20% | 2.512 | 1.457 | 1.945 | 3.414 | 3.275 | 2.432 | 2.510 | 3.380 |
| .001 | $\widehat{OR}$ | 40% | 3.590 | 1.818 | 2.896 | 4.258 | 4.576 | 4.373 | 3.650 | 5.937 |
| .001 | $\widehat{OR}$ | 60% | 5.214 | 2.419 | 5.392 | 5.962 | 6.630 | 7.631 | 5.042 | 14.343 |
| .001 | $\widehat{OR}$ | 80% | 10.376 | 4.273 | 22.679 | 13.238 | 12.239 | 12.270 | 9.508 | 81.545 |
| B-.05 | $\widehat{OR}$ | 0% | 1.135 | 1.008 | 1.022 | 2.530 | 1.075 | 1.160 | 1.209 | 1.185 |
| B-.05 | $\widehat{OR}$ | 20% | 2.512 | 1.467 | 1.898 | 3.414 | 3.669 | 2.432 | 2.516 | 3.083 |
| B-.05 | $\widehat{OR}$ | 40% | 3.555 | 1.826 | 2.694 | 4.258 | 5.275 | 4.421 | 3.655 | 4.752 |
| B-.05 | $\widehat{OR}$ | 60% | 5.118 | 2.422 | 4.523 | 5.962 | 7.704 | 7.848 | 5.007 | 8.942 |
| B-.05 | $\widehat{OR}$ | 80% | 10.083 | 4.247 | 13.869 | 13.238 | 14.332 | 12.324 | 9.745 | 28.055 |
| 1 | $F$ | 0% | 1.002 | 1.000 | 1.000 | 2.013 | 1.000 | 1.106 | 1.172 | 1.000 |
| 1 | $F$ | 20% | 1.766 | 1.245 | 1.479 | 2.323 | 2.370 | 2.218 | 2.335 | 2.087 |
| 1 | $F$ | 40% | 2.681 | 1.552 | 2.204 | 2.732 | 3.564 | 3.707 | 3.328 | 3.680 |
| 1 | $F$ | 60% | 3.987 | 2.071 | 3.829 | 3.629 | 5.238 | 5.837 | 4.608 | 7.419 |
| 1 | $F$ | 80% | 7.676 | 3.563 | 10.457 | 6.785 | 9.355 | 9.824 | 8.295 | 22.561 |
| .05 | $F$ | 0% | 1.080 | 1.005 | 1.009 | 2.013 | 1.024 | 1.106 | 1.172 | 1.069 |
| .05 | $F$ | 20% | 2.330 | 1.419 | 2.149 | 2.323 | 2.665 | 2.218 | 2.335 | 3.818 |
| .05 | $F$ | 40% | 3.343 | 1.783 | 3.806 | 2.732 | 3.828 | 3.707 | 3.328 | 8.913 |
| .05 | $F$ | 60% | 4.957 | 2.418 | 9.899 | 3.629 | 5.542 | 5.837 | 4.608 | 27.462 |
| .05 | $F$ | 80% | 10.400 | 4.474 | 42.308 | 6.785 | 9.895 | 9.824 | 8.295 | 75.519 |
| .001 | $F$ | 0% | 1.114 | 1.005 | 1.010 | 2.013 | 1.063 | 1.106 | 1.172 | 1.092 |
| .001 | $F$ | 20% | 2.333 | 1.424 | 1.890 | 2.323 | 3.100 | 2.265 | 2.335 | 2.928 |
| .001 | $F$ | 40% | 3.296 | 1.758 | 2.780 | 2.732 | 4.262 | 3.893 | 3.334 | 4.970 |
| .001 | $F$ | 60% | 4.679 | 2.308 | 5.058 | 3.629 | 6.066 | 6.154 | 4.609 | 11.204 |
| .001 | $F$ | 80% | 8.876 | 3.985 | 20.020 | 6.785 | 10.869 | 10.039 | 8.318 | 52.744 |
| B-.05 | $F$ | 0% | 1.126 | 1.005 | 1.013 | 2.013 | 1.066 | 1.106 | 1.172 | 1.099 |
| B-.05 | $F$ | 20% | 2.331 | 1.431 | 1.843 | 2.323 | 3.456 | 2.265 | 2.339 | 2.642 |
| B-.05 | $F$ | 40% | 3.279 | 1.763 | 2.596 | 2.732 | 4.879 | 3.927 | 3.343 | 3.976 |
| B-.05 | $F$ | 60% | 4.617 | 2.306 | 4.308 | 3.629 | 6.997 | 6.346 | 4.601 | 7.136 |
| B-.05 | $F$ | 80% | 8.474 | 3.970 | 12.858 | 6.785 | 12.628 | 10.080 | 8.451 | 20.368 |

Table S2: Quintiles of binary association measures for pairs in each dataset for different evidential thresholds. "B" indicates Bonferroni correction.

| TWER | BAM | quant. | Columbia | MedPAR(3) | MedPAR(5) | SctHans | Michigan | Stanford | Columbia* | MIMIC |
|---|---|---|---|---|---|---|---|---|---|---|
| 1 | $\phi$ | 0% | 0.000 | 0.000 | 0.000 | 0.093 | 0.000 | 0.001 | 0.004 | 0.000 |
| 1 | $\phi$ | 20% | 0.001 | 0.000 | 0.000 | 0.110 | 0.004 | 0.012 | 0.007 | 0.006 |
| 1 | $\phi$ | 40% | 0.003 | 0.001 | 0.000 | 0.124 | 0.006 | 0.019 | 0.011 | 0.010 |
| 1 | $\phi$ | 60% | 0.006 | 0.001 | 0.001 | 0.146 | 0.009 | 0.032 | 0.019 | 0.016 |
| 1 | $\phi$ | 80% | 0.013 | 0.003 | 0.002 | 0.192 | 0.017 | 0.066 | 0.039 | 0.029 |
| .05 | $\phi$ | 0% | 0.002 | 0.000 | 0.000 | 0.093 | 0.000 | 0.001 | 0.004 | 0.000 |
| .05 | $\phi$ | 20% | 0.004 | 0.001 | 0.001 | 0.110 | 0.004 | 0.012 | 0.007 | 0.018 |
| .05 | $\phi$ | 40% | 0.006 | 0.002 | 0.001 | 0.124 | 0.006 | 0.019 | 0.011 | 0.024 |
| .05 | $\phi$ | 60% | 0.010 | 0.003 | 0.002 | 0.146 | 0.010 | 0.032 | 0.019 | 0.032 |
| .05 | $\phi$ | 80% | 0.021 | 0.007 | 0.003 | 0.192 | 0.018 | 0.066 | 0.039 | 0.050 |
| .001 | $\phi$ | 0% | 0.003 | 0.000 | 0.000 | 0.093 | 0.000 | 0.001 | 0.004 | 0.000 |
| .001 | $\phi$ | 20% | 0.006 | 0.002 | 0.001 | 0.110 | 0.005 | 0.012 | 0.007 | 0.026 |
| .001 | $\phi$ | 40% | 0.010 | 0.002 | 0.002 | 0.124 | 0.007 | 0.020 | 0.011 | 0.033 |
| .001 | $\phi$ | 60% | 0.015 | 0.004 | 0.003 | 0.146 | 0.011 | 0.034 | 0.020 | 0.046 |
| .001 | $\phi$ | 80% | 0.029 | 0.010 | 0.007 | 0.192 | 0.020 | 0.068 | 0.040 | 0.081 |
| B-.05 | $\phi$ | 0% | 0.004 | 0.000 | 0.000 | 0.093 | 0.000 | 0.001 | 0.004 | 0.007 |
| B-.05 | $\phi$ | 20% | 0.007 | 0.002 | 0.002 | 0.110 | 0.006 | 0.013 | 0.008 | 0.034 |
| B-.05 | $\phi$ | 40% | 0.010 | 0.003 | 0.003 | 0.124 | 0.009 | 0.020 | 0.012 | 0.042 |
| B-.05 | $\phi$ | 60% | 0.016 | 0.005 | 0.004 | 0.146 | 0.013 | 0.035 | 0.020 | 0.054 |
| B-.05 | $\phi$ | 80% | 0.030 | 0.011 | 0.008 | 0.192 | 0.022 | 0.069 | 0.040 | 0.080 |
| 1 | | 0% | 0.000 | 0.000 | 0.000 | 0.233 | 0.000 | 0.034 | 0.038 | 0.000 |
| 1 | | 20% | 0.090 | 0.024 | 0.035 | 0.307 | 0.105 | 0.136 | 0.135 | 0.106 |
| 1 | | 40% | 0.157 | 0.048 | 0.070 | 0.344 | 0.158 | 0.212 | 0.178 | 0.183 |
| 1 | | 60% | 0.232 | 0.078 | 0.115 | 0.408 | 0.210 | 0.304 | 0.225 | 0.269 |
| 1 | | 80% | 0.348 | 0.130 | 0.195 | 0.526 | 0.286 | 0.399 | 0.317 | 0.404 |
| .05 | | 0% | 0.016 | 0.002 | 0.003 | 0.233 | 0.006 | 0.034 | 0.038 | 0.026 |
| .05 | | 20% | 0.119 | 0.046 | 0.079 | 0.307 | 0.120 | 0.136 | 0.135 | 0.215 |
| .05 | | 40% | 0.163 | 0.071 | 0.127 | 0.344 | 0.168 | 0.212 | 0.178 | 0.318 |
| .05 | | 60% | 0.210 | 0.103 | 0.205 | 0.408 | 0.217 | 0.304 | 0.225 | 0.423 |
| .05 | | 80% | 0.295 | 0.163 | 0.308 | 0.526 | 0.293 | 0.399 | 0.317 | 0.546 |
| .001 | | 0% | 0.023 | 0.003 | 0.003 | 0.233 | 0.014 | 0.034 | 0.038 | 0.038 |
| .001 | | 20% | 0.131 | 0.050 | 0.074 | 0.307 | 0.140 | 0.138 | 0.135 | 0.206 |
| .001 | | 40% | 0.173 | 0.075 | 0.111 | 0.344 | 0.182 | 0.228 | 0.179 | 0.277 |
| .001 | | 60% | 0.221 | 0.107 | 0.165 | 0.408 | 0.230 | 0.308 | 0.225 | 0.377 |
| .001 | | 80% | 0.304 | 0.166 | 0.280 | 0.526 | 0.305 | 0.401 | 0.318 | 0.556 |
| B-.05 | | 0% | 0.023 | 0.003 | 0.007 | 0.233 | 0.016 | 0.038 | 0.038 | 0.052 |
| B-.05 | | 20% | 0.133 | 0.052 | 0.077 | 0.307 | 0.161 | 0.140 | 0.137 | 0.210 |
| B-.05 | | 40% | 0.174 | 0.078 | 0.112 | 0.344 | 0.203 | 0.230 | 0.179 | 0.270 |
| B-.05 | | 60% | 0.222 | 0.111 | 0.161 | 0.408 | 0.249 | 0.314 | 0.227 | 0.349 |
| B-.05 | | 80% | 0.305 | 0.171 | 0.258 | 0.526 | 0.324 | 0.403 | 0.319 | 0.485 |

Table S3: Log-likelihood ratios $R$ and two-sided p-values $p$ from likelihood-ratio tests between four families of models to degree sequence tails. LRTs were performed for graphs constructed from each dataset using two p-value significance thresholds and corrections for family-wise error rate and for false discovery rate. Family 1 is preferred when $R > 0$, Family 2 when $R < 0$.

| Dataset | Family 1 | Family 2 | $R$ ($10^{-2}$) | $p$ ($10^{-2}$) | $R$ ($10^{-2}$, B) | $p$ ($10^{-2}$, B) | $R$ ($10^{-2}$, B-H) | $p$ ($10^{-2}$, B-H) | $R$ ($10^{-5}$) | $p$ ($10^{-5}$) | $R$ ($10^{-5}$, B) | $p$ ($10^{-5}$, B) | $R$ ($10^{-5}$, B-H) | $p$ ($10^{-5}$, B-H) |
|---|---|---|---|---|---|---|---|---|---|---|---|---|---|---|
| Columbia | Poisson | power law | -11.3 | 0.000 | -0.4 | 0.684 | -11.8 | 0.000 | -10.5 | 0.000 | -0.8 | 0.433 | -9.7 | 0.000 |
| Columbia | Poisson | exponential | -13.1 | 0.000 | -0.9 | 0.357 | -13.4 | 0.000 | -11.9 | 0.000 | -1.5 | 0.142 | -11.1 | 0.000 |
| Columbia | Poisson | log-normal | -13.1 | 0.000 | -1.9 | 0.060 | -13.2 | 0.000 | -11.6 | 0.000 | -2.6 | 0.010 | -10.9 | 0.000 |
| Columbia | power law | exponential | -15.8 | 0.000 | -4.9 | 0.000 | -9.7 | 0.000 | -5.4 | 0.000 | -6.6 | 0.000 | -6.8 | 0.000 |
| Columbia | power law | log-normal | -12.7 | 0.000 | -2.5 | 0.011 | -9.7 | 0.000 | -5.3 | 0.000 | -3.4 | 0.001 | -6.2 | 0.000 |
| Columbia | exponential | log-normal | 2.3 | 0.021 | -1.7 | 0.092 | 3.7 | 0.000 | 4.0 | 0.000 | -2.1 | 0.035 | 4.6 | 0.000 |
| MedPAR(3) | Poisson | power law | -28.3 | 0.000 | -23.9 | 0.000 | -27.6 | 0.000 | -24.5 | 0.000 | -23.4 | 0.000 | -25.0 | 0.000 |
| MedPAR(3) | Poisson | exponential | -29.1 | 0.000 | -24.4 | 0.000 | -28.2 | 0.000 | -25.0 | 0.000 | -23.9 | 0.000 | -25.6 | 0.000 |
| MedPAR(3) | Poisson | log-normal | -29.0 | 0.000 | -24.4 | 0.000 | -28.1 | 0.000 | -25.0 | 0.000 | -23.9 | 0.000 | -25.6 | 0.000 |
| MedPAR(3) | power law | exponential | -36.3 | 0.000 | -25.6 | 0.000 | -22.9 | 0.000 | -26.3 | 0.000 | -24.5 | 0.000 | -25.8 | 0.000 |
| MedPAR(3) | power law | log-normal | -29.7 | 0.000 | -21.9 | 0.000 | -19.3 | 0.000 | -20.7 | 0.000 | -22.2 | 0.000 | -21.3 | 0.000 |
| MedPAR(3) | exponential | log-normal | 14.3 | 0.000 | 6.1 | 0.000 | 17.1 | 0.000 | 5.5 | 0.000 | 6.4 | 0.000 | 9.8 | 0.000 |
| MedPAR(5) | Poisson | power law | -7.6 | 0.000 | -9.9 | 0.000 | -10.1 | 0.000 | -10.5 | 0.000 | -6.5 | 0.000 | -10.0 | 0.000 |
| MedPAR(5) | Poisson | exponential | -7.6 | 0.000 | -9.9 | 0.000 | -9.5 | 0.000 | -9.8 | 0.000 | -6.5 | 0.000 | -10.1 | 0.000 |
| MedPAR(5) | Poisson | log-normal | -7.6 | 0.000 | -9.9 | 0.000 | -10.1 | 0.000 | -10.5 | 0.000 | -6.5 | 0.000 | -10.1 | 0.000 |
| MedPAR(5) | power law | exponential | -1.6 | 0.107 | 2.6 | 0.008 | 12.8 | 0.000 | 13.2 | 0.000 | 0.0 | 0.965 | 1.1 | 0.253 |
| MedPAR(5) | power law | log-normal | -5.0 | 0.000 | -10.7 | 0.000 | -11.7 | 0.000 | -11.1 | 0.000 | -5.3 | 0.000 | -9.2 | 0.000 |
| MedPAR(5) | exponential | log-normal | -2.3 | 0.021 | -7.7 | 0.000 | -12.9 | 0.000 | -13.3 | 0.000 | -2.9 | 0.003 | -6.5 | 0.000 |
| Sct.Hans | Poisson | power law | -4.3 | 0.000 | -5.3 | 0.000 | -4.3 | 0.000 | -3.8 | 0.000 | -2.2 | 0.028 | -4.9 | 0.000 |
| Sct.Hans | Poisson | exponential | -5.2 | 0.000 | -6.4 | 0.000 | -5.2 | 0.000 | -5.0 | 0.000 | -4.2 | 0.000 | -6.1 | 0.000 |
| Sct.Hans | Poisson | log-normal | -5.1 | 0.000 | -6.1 | 0.000 | -5.1 | 0.000 | -4.9 | 0.000 | -3.7 | 0.000 | -5.8 | 0.000 |
| Sct.Hans | power law | exponential | -3.8 | 0.000 | -1.2 | 0.228 | -3.8 | 0.000 | -3.8 | 0.000 | -2.5 | 0.013 | -1.2 | 0.238 |
| Sct.Hans | power law | log-normal | -3.5 | 0.000 | -3.6 | 0.000 | -3.5 | 0.000 | -3.7 | 0.000 | -3.2 | 0.001 | -3.4 | 0.001 |
| Sct.Hans | exponential | log-normal | 2.7 | 0.007 | -1.4 | 0.174 | 2.7 | 0.007 | 1.9 | 0.060 | 0.6 | 0.548 | -1.2 | 0.230 |
| Michigan | Poisson | power law | -14.6 | 0.000 | -13.0 | 0.000 | -16.8 | 0.000 | -14.1 | 0.000 | -10.5 | 0.000 | -17.7 | 0.000 |
| Michigan | Poisson | exponential | -14.7 | 0.000 | -13.0 | 0.000 | -16.8 | 0.000 | -14.1 | 0.000 | -10.6 | 0.000 | -17.8 | 0.000 |
| Michigan | Poisson | log-normal | -14.7 | 0.000 | -13.0 | 0.000 | -16.8 | 0.000 | -14.1 | 0.000 | -10.6 | 0.000 | -17.8 | 0.000 |
| Michigan | power law | exponential | -5.2 | 0.000 | -3.6 | 0.000 | -6.0 | 0.000 | -3.3 | 0.001 | -5.4 | 0.000 | -6.0 | 0.000 |
| Michigan | power law | log-normal | -7.2 | 0.000 | -6.8 | 0.000 | -10.0 | 0.000 | -8.3 | 0.000 | -4.9 | 0.000 | -12.3 | 0.000 |
| Michigan | exponential | log-normal | -0.9 | 0.372 | -2.2 | 0.029 | -3.0 | 0.003 | -3.3 | 0.001 | 2.1 | 0.040 | -3.8 | 0.000 |
| Stanford | Poisson | power law | -1.1 | 0.255 | -2.7 | 0.008 | -1.0 | 0.325 | -2.6 | 0.009 | -3.7 | 0.000 | -2.5 | 0.013 |
| Stanford | Poisson | exponential | -3.4 | 0.001 | -4.8 | 0.000 | -3.3 | 0.001 | -4.9 | 0.000 | -5.1 | 0.000 | -4.6 | 0.000 |
| Stanford | Poisson | log-normal | -3.9 | 0.000 | -4.5 | 0.000 | -3.9 | 0.000 | -4.6 | 0.000 | -4.7 | 0.000 | -4.4 | 0.000 |
| Stanford | power law | exponential | -6.1 | 0.000 | -3.3 | 0.001 | -6.3 | 0.000 | -3.5 | 0.000 | -1.1 | 0.293 | -3.4 | 0.001 |
| Stanford | power law | log-normal | -4.7 | 0.000 | -3.7 | 0.000 | -4.8 | 0.000 | -3.7 | 0.000 | -2.6 | 0.009 | -3.8 | 0.000 |
| Stanford | exponential | log-normal | -0.3 | 0.728 | 1.2 | 0.228 | -0.6 | 0.553 | 1.8 | 0.073 | -0.9 | 0.359 | 1.1 | 0.267 |
| Columbia* | Poisson | power law | 1.0 | 0.328 | 1.5 | 0.132 | 1.8 | 0.069 | 2.6 | 0.009 | 3.7 | 0.000 | 3.4 | 0.001 |
| Columbia* | Poisson | exponential | -0.7 | 0.510 | 2.1 | 0.034 | 1.0 | 0.337 | 1.7 | 0.086 | 1.7 | 0.091 | 1.7 | 0.091 |
| Columbia* | Poisson | log-normal | -1.9 | 0.061 | -1.2 | 0.237 | 0.1 | 0.939 | 0.9 | 0.392 | -0.6 | 0.575 | -0.1 | 0.888 |
| Columbia* | power law | exponential | -6.9 | 0.000 | -1.0 | 0.332 | -4.4 | 0.000 | -5.2 | 0.000 | -10.4 | 0.000 | -9.2 | 0.000 |
| Columbia* | power law | log-normal | -3.7 | 0.000 | -1.4 | 0.176 | -2.4 | 0.017 | -2.8 | 0.006 | -5.3 | 0.000 | -4.6 | 0.000 |
| Columbia* | exponential | log-normal | -1.3 | 0.179 | -1.4 | 0.175 | -1.4 | 0.164 | -1.8 | 0.073 | -2.9 | 0.004 | -2.6 | 0.010 |
| MIMIC | Poisson | power law | -13.7 | 0.000 | -9.5 | 0.000 | -14.2 | 0.000 | -9.7 | 0.000 | -10.8 | 0.000 | -10.7 | 0.000 |
| MIMIC | Poisson | exponential | -13.8 | 0.000 | -9.6 | 0.000 | -14.1 | 0.000 | -9.8 | 0.000 | -10.9 | 0.000 | -10.9 | 0.000 |
| MIMIC | Poisson | log-normal | -13.8 | 0.000 | -9.6 | 0.000 | -14.2 | 0.000 | -9.8 | 0.000 | -10.9 | 0.000 | -10.8 | 0.000 |
| MIMIC | power law | exponential | 1.5 | 0.142 | -1.3 | 0.211 | 12.8 | 0.000 | -2.6 | 0.009 | -1.2 | 0.227 | -1.7 | 0.090 |
| MIMIC | power law | log-normal | -11.7 | 0.000 | -6.3 | 0.000 | -11.6 | 0.000 | -5.8 | 0.000 | -7.9 | 0.000 | -7.2 | 0.000 |
| MIMIC | exponential | log-normal | -8.7 | 0.000 | -3.4 | 0.001 | -17.9 | 0.000 | -2.3 | 0.020 | -4.4 | 0.000 | -3.6 | 0.000 |

Figure S7: Diverging-color tilings of log-likelihood ratios $R$ from likelihood-ratio tests (LRT) between fits of different families of models to degree sequence tails. LRTs were performed for graphs constructed from each dataset using evidential cutoff $\alpha < 0.05$ and both corrections. Family 1 is preferred when $R > 0$, Family 2 when $R < 0$. The boundary color of each tile indicates whether $R$ is positive or negative. The p-value from the LRT is printed on each tile.

Table S4: Tail index ($\xi$) and tail exponent ($\gamma$) estimators that were successfully estimated on comorbidity networks.

| Dataset | p-value | Correction | $\xi_{\text{Hill}}$ | $\xi_{\text{moments}}$ | $\xi_{\text{kernel}}$ | $\gamma_{\text{Hill}}$ | $\gamma_{\text{moments}}$ | $\gamma_{\text{kernel}}$ |
|---|---|---|---|---|---|---|---|---|
| MedPAR(3) | 0.01 | FDR | 0.125 | -0.275 | -1.172 | 9.000 | -2.636 | 0.147 |
| MedPAR(3) | 0.00001 | FDR | 0.156 | -0.394 | -0.325 | 7.410 | -1.538 | -2.077 |
| MedPAR(3) | 0.01 | FWER | 0.114 | -0.358 | -0.452 | 9.772 | -1.793 | -1.212 |
| MedPAR(3) | 0.00001 | FWER | 0.158 | -0.157 | -0.310 | 7.329 | -5.369 | -2.226 |
| MedPAR(3) | 0.01 | – | 0.130 | -0.411 | -0.479 | 8.692 | -1.433 | -1.088 |
| MedPAR(3) | 0.00001 | – | 0.143 | -0.416 | -0.441 | 7.993 | -1.404 | -1.268 |
| MedPAR(5) | 0.01 | FDR | 0.239 | 0.014 | 0.482 | 5.184 | 72.429 | 3.075 |
| MedPAR(5) | 0.00001 | FDR | 0.266 | 0.194 | 0.413 | 4.759 | 6.155 | 3.421 |
| MedPAR(5) | 0.01 | FWER | 0.284 | 0.071 | 0.398 | 4.521 | 15.085 | 3.513 |
| MedPAR(5) | 0.00001 | FWER | 0.254 | 0.134 | 0.425 | 4.937 | 8.463 | 3.353 |
| MedPAR(5) | 0.01 | – | 0.204 | -121.754 | 0.407 | 5.902 | 0.992 | 3.457 |
| MedPAR(5) | 0.00001 | – | 0.299 | 0.014 | 0.487 | 4.344 | 72.429 | 3.053 |
| Michigan | 0.01 | FDR | 0.020 | -0.093 | -0.161 | 51.000 | -9.753 | -5.211 |
| Michigan | 0.00001 | FDR | 0.023 | -0.094 | 0.527 | 44.478 | -9.638 | 2.898 |
| Michigan | 0.01 | FWER | 0.020 | -0.137 | 0.580 | 51.000 | -6.299 | 2.724 |
| Michigan | 0.00001 | FWER | 0.013 | -0.168 | 0.087 | 77.923 | -4.952 | 12.494 |
| Michigan | 0.01 | – | 0.017 | -0.102 | -0.162 | 59.824 | -8.804 | -5.173 |
| Michigan | 0.00001 | – | 0.021 | -0.073 | -0.533 | 48.619 | -12.699 | -0.876 |
| MIMIC | 0.01 | FDR | 0.176 | 0.031 | 0.262 | 6.682 | 33.258 | 4.817 |
| MIMIC | 0.00001 | FDR | 0.112 | -0.249 | 0.350 | 9.929 | -3.016 | 3.857 |
| MIMIC | 0.01 | FWER | 0.171 | -0.220 | 0.378 | 6.848 | -3.545 | 3.646 |
| MIMIC | 0.00001 | FWER | 0.223 | -0.112 | 0.400 | 5.484 | -7.929 | 3.500 |
| MIMIC | 0.01 | – | 0.082 | 0.149 | 0.589 | 13.195 | 7.711 | 2.698 |
| MIMIC | 0.00001 | – | 0.172 | -0.003 | 0.260 | 6.814 | -332.333 | 4.846 |

**Single-valued summary statistics**

A priori of regression modeling, several global network statistics are clearly dependent on the choice of TWER ($\alpha$). After some basic transformations to increase linearity, their values at each graph we constructed are plotted against $\alpha$ in Figure S8. These plots suggest that the effect of $\alpha$ on many global statistics varies by dataset. For example, the density increases more dramatically in the MedPAR(5) and MIMIC-III data sets as the evidential threshold is weakened, with corresponding changes in other statistics including the proportion of nodes in the largest component and the skewness of the degree sequence as measured by the Gini index and the $\mu$ parameter of the log-normal fit. However, this difference may be an artifact of the limited data provided for the other networks; co-occurrence data for most pairs was only available for MedPAR(3) and MedPAR(5) and for all pairs for MIMIC and Columbia, and indeed the patterns for Columbia and MedPAR(3) are less pronounced but similar to those for MedPAR(5) and MIMIC. Different populations seem to exhibit different comorbidity network structure in terms of the raw values of several global statistics, but our observations are consistent with the choice of TWER having a broadly similar effect on this structure regardless of the source.

Table S5: Values of several global statistics calculated on comorbidity networks constructed over a range of data sets and parameter settings. See the section on global properties in the main text.

Figure S8: Mean values of several global statistics calculated on comorbidity networks constructed over a range of data sets and parameter settings, plotted against several measures of network size. See the section on global properties in the main text.

Table S6: LMs of network statistics on data source and test-wise error rate.

| | *Dependent variable:* | | | | | | | | |
|---|---|---|---|---|---|---|---|---|---|
| | LCP | $r$ | $G$ | $\bar{k}$ | $\hat{\mu}$ | $\hat{\sigma}$ | $\bar{\ell}$ | $Q$ | $C$ |
| | (1) | (2) | (3) | (4) | (5) | (6) | (7) | (8) | (9) |
| Columbia | $0.40^{***}$ | $-0.13^{***}$ | $-0.20^{***}$ | $-27.40^{*}$ | $0.62^{**}$ | $-0.51^{***}$ | $-0.66^{***}$ | $-0.16^{***}$ | $0.13^{***}$ |
| | (0.05) | (0.02) | (0.02) | (15.89) | (0.22) | (0.11) | (0.11) | (0.01) | (0.01) |
| MedPAR(3) | $0.40^{***}$ | $-0.14^{***}$ | $-0.20^{***}$ | $110.30^{***}$ | $2.34^{***}$ | $-0.47^{***}$ | $-0.74^{***}$ | $-0.13^{***}$ | $0.05^{***}$ |
| | (0.05) | (0.02) | (0.02) | (15.89) | (0.22) | (0.11) | (0.11) | (0.01) | (0.01) |
| MedPAR(5) | $0.19^{***}$ | $-0.22^{***}$ | $0.15^{***}$ | 14.99 | $1.16^{***}$ | $0.42^{***}$ | 0.15 | $-0.11^{***}$ | $-0.27^{***}$ |
| | (0.05) | (0.02) | (0.02) | (15.89) | (0.22) | (0.11) | (0.11) | (0.01) | (0.01) |
| Sct.Hans | $0.27^{***}$ | $0.05^{**}$ | $-0.08^{***}$ | $-52.96^{**}$ | $-1.46^{***}$ | 0.04 | $0.75^{***}$ | $0.10^{***}$ | $-0.21^{***}$ |
| | (0.05) | (0.02) | (0.02) | (15.89) | (0.22) | (0.11) | (0.11) | (0.01) | (0.01) |
| Michigan | $0.45^{***}$ | $-0.41^{***}$ | $-0.02$ | $495.20^{***}$ | $2.95^{***}$ | $0.20^{*}$ | $-0.66^{***}$ | $-0.18^{***}$ | $-0.08^{***}$ |
| | (0.05) | (0.02) | (0.02) | (15.89) | (0.22) | (0.11) | (0.11) | (0.01) | (0.01) |
| Stanford | $-0.29^{***}$ | $0.65^{***}$ | $-0.18^{***}$ | $-53.99^{**}$ | $-1.87^{***}$ | $-0.03$ | $-0.46^{***}$ | $0.09^{***}$ | $0.21^{***}$ |
| | (0.05) | (0.02) | (0.02) | (15.89) | (0.22) | (0.11) | (0.11) | (0.01) | (0.01) |
| Columbia* | $-0.26^{***}$ | $0.11^{***}$ | $-0.28^{***}$ | $-49.04^{**}$ | $-0.74^{**}$ | $-0.48^{***}$ | $-1.06^{***}$ | $-0.16^{***}$ | $0.24^{***}$ |
| | (0.05) | (0.02) | (0.02) | (15.89) | (0.22) | (0.11) | (0.11) | (0.01) | (0.01) |
| MIMIC | $0.19^{**}$ | 0.02 | $0.08^{***}$ | $-11.55$ | 0.18 | $0.43^{***}$ | $0.70^{***}$ | $-0.01$ | $-0.18^{***}$ |
| | (0.05) | (0.02) | (0.02) | (15.89) | (0.22) | (0.11) | (0.11) | (0.01) | (0.01) |
| $\log(p)$ | $0.02^{***}$ | $-0.001$ | $-0.01^{***}$ | $6.60^{***}$ | $0.05^{**}$ | $-0.01$ | $-0.03^{***}$ | $-0.01^{***}$ | $-0.002^{*}$ |
| | (0.004) | (0.001) | (0.001) | (1.16) | (0.02) | (0.01) | (0.01) | (0.001) | (0.001) |
| Observations | 48 | 48 | 48 | 48 | 48 | 48 | 48 | 48 | 48 |
| Adjusted $R^2$ | 0.87 | 0.99 | 0.94 | 0.97 | 0.93 | 0.73 | 0.90 | 0.94 | 0.99 |

*Note:* $^{*}p<0.1$; $^{**}p<0.01$; $^{***}p<0.001$

A plausible alternative modeling approach would have been to treat the data source as a random effect, sampled from an underlying distribution, on the values of the network statistics. Though only few data sets were used, we fit hierarchical regression models to the same data as a robustness check. The results were consistent with those for the classical models (Tables S7, S9, S8, and S10). The Akaike Information Criterion (Burnham and Anderson 2004), which penalizes increased model complexity as well as un-accounted for variance as a corrective to overfitting, favored the classical models with respect to every network statistic (Table S11).

Table S7: Hierarchical models of network statistics on test-wise error rate, grouped by data source

| | *Dependent variable:* | | | | | | | | |
|---|---|---|---|---|---|---|---|---|---|
| | LCP | $r$ | $G$ | $\bar{k}$ | $\hat{\mu}$ | $\hat{\sigma}$ | $\bar{\ell}$ | $Q$ | $C$ |
| | (1) | (2) | (3) | (4) | (5) | (6) | (7) | (8) | (9) |
| $\log(p)$ | $0.02^{*}$ | 0.003 | $-0.01^{***}$ | $6.20^{***}$ | $-0.03$ | 0.03 | $-0.05^{*}$ | $-0.01^{***}$ | $-0.003^{*}$ |
| | (0.01) | (0.003) | (0.003) | (1.63) | (0.05) | (0.02) | (0.02) | (0.002) | (0.001) |
| Const. | $0.91^{***}$ | 0.01 | $0.50^{***}$ | $154.63^{*}$ | $3.82^{***}$ | $0.86^{***}$ | $2.11^{***}$ | $0.12^{**}$ | $0.44^{***}$ |
| | (0.12) | (0.11) | (0.06) | (69.61) | (0.68) | (0.20) | (0.29) | (0.04) | (0.07) |
| Observations | 24 | 24 | 24 | 24 | 24 | 24 | 24 | 24 | 24 |
| Log Likelihood | 7.94 | 22.38 | 26.58 | -116.37 | -29.78 | -8.04 | -11.24 | 38.11 | 39.92 |
| Akaike Inf. Crit. | -7.87 | -36.76 | -45.16 | 240.73 | 67.56 | 24.09 | 30.48 | -68.21 | -71.84 |
| Bayesian Inf. Crit. | -3.16 | -32.04 | -40.44 | 245.44 | 72.27 | 28.80 | 35.19 | -63.50 | -67.13 |
| *Note:* | | | | | | | | | $^{*}$p<0.1; $^{**}$p<0.01; $^{***}$p<0.001 |

Table S8: Hierarchical models of network statistics on test-wise error rate and binary association measure, grouped by data source

| | *Dependent variable:* | | | | | | | | |
|---|---|---|---|---|---|---|---|---|---|
| | LCP | $r$ | $G$ | $\bar{k}$ | $\hat{\mu}$ | $\hat{\sigma}$ | $\bar{\ell}$ | $Q$ | $C$ |
| | (1) | (2) | (3) | (4) | (5) | (6) | (7) | (8) | (9) |
| $\log(p)$ | $0.02^{***}$ | $-0.003$ | $-0.01^{***}$ | $1.66^{*}$ | $0.05^{***}$ | $-0.01^{**}$ | $-0.06^{**}$ | $-0.01^{***}$ | $-0.01^{***}$ |
| | (0.002) | (0.002) | (0.001) | (0.79) | (0.01) | (0.003) | (0.02) | (0.002) | (0.001) |
| $F \times \theta_F$ | $-0.01^{***}$ | $0.003^{***}$ | $0.002^{***}$ | $-1.12^{***}$ | $-0.03^{***}$ | $-0.003^{**}$ | $0.02^{***}$ | $0.003^{***}$ | $-0.003^{***}$ |
| | (0.001) | (0.001) | (0.0003) | (0.20) | (0.004) | (0.001) | (0.005) | (0.001) | (0.0003) |
| $\widehat{OR} \times \theta_{\widehat{OR}}$ | $-3.23^{***}$ | $2.20^{***}$ | $1.47^{***}$ | $-415.08^{***}$ | $-19.28^{***}$ | 0.49 | $3.88^{**}$ | $0.59^{***}$ | $0.52^{***}$ |
| | (0.15) | (0.17) | (0.09) | (59.38) | (1.08) | (0.30) | (1.38) | (0.16) | (0.11) |
| $\phi \times \theta_\phi$ | $-0.01^{***}$ | $0.001^{*}$ | $0.002^{***}$ | $-1.15^{***}$ | $-0.04^{***}$ | $-0.002^{*}$ | $0.02^{***}$ | $0.002^{***}$ | $-0.003^{***}$ |
| | (0.001) | (0.001) | (0.0003) | (0.20) | (0.004) | (0.001) | (0.005) | (0.001) | (0.0003) |
| $r_t \times \theta_{r_t}$ | $-0.62^{***}$ | $0.20^{***}$ | $0.19^{***}$ | $-122.67^{***}$ | $-4.47^{***}$ | $-0.04$ | $2.71^{***}$ | $0.35^{***}$ | $-0.21^{***}$ |
| | (0.05) | (0.05) | (0.03) | (17.67) | (0.30) | (0.08) | (0.41) | (0.05) | (0.03) |
| Observations | 576 | 568 | 576 | 576 | 504 | 504 | 576 | 576 | 568 |
| Log Likelihood | 71.57 | 65.09 | 395.39 | -3,316.68 | -800.80 | -162.98 | -1,167.92 | 79.83 | 313.28 |
| Akaike Inf. Crit. | -129.13 | -116.17 | -776.79 | 6,647.37 | 1,615.61 | 339.96 | 2,349.83 | -145.67 | -612.57 |
| Bayesian Inf. Crit. | -98.64 | -85.78 | -746.30 | 6,677.86 | 1,645.16 | 369.52 | 2,380.33 | -115.18 | -582.17 |
| *Note:* | | | | | | | | | $^{*}$p<0.1; $^{**}$p<0.01; $^{***}$p<0.001 |

Table S9: Variance decomposition for hierarchical models of network statistics on test-wise error rate, grouped by data source

| | LCP | $\bar{k}$ | $G$ | $\hat{\mu}$ | $\hat{\sigma}$ | $r$ | $C$ | $\bar{\ell}$ | $Q$ |
|---|---|---|---|---|---|---|---|---|---|
| $\sigma_1$(source) | 0.302 | 194.102 | 0.146 | 1.662 | 0.354 | 0.315 | 0.198 | 0.702 | 0.105 |
| $\sigma$(resid.) | 0.077 | 14.988 | 0.031 | 0.428 | 0.203 | 0.029 | 0.011 | 0.186 | 0.017 |

Table S10: Variance decomposition for hierarchical models of network statistics on test-wise error rate and binary association measure, grouped by data source

| | LCP | $\bar{k}$ | $G$ | $\hat{\mu}$ | $\hat{\sigma}$ | $r$ | $C$ | $\bar{\ell}$ | $Q$ |
|---|---|---|---|---|---|---|---|---|---|
| $\sigma_1$(source) | 0.912 | 110.686 | 0.497 | 3.874 | 0.778 | 0.162 | 0.377 | 2.582 | 0.245 |
| $\sigma$(resid.) | 0.196 | 75.361 | 0.111 | 1.103 | 0.309 | 0.203 | 0.128 | 1.749 | 0.197 |

Table S11: Akaike information criteria for each model fitted to the values taken by each network statistic

| | Model 1 | Model 2 | Model 3 | Model 4 |
|---|---|---|---|---|
| LCP | −69 | −226 | −8 | −129 |
| $\bar{k}$ | 478 | 6629 | 241 | 6647 |
| $G$ | −167 | −878 | −45 | −777 |
| $\hat{\mu}$ | 68 | 1544 | 68 | 1616 |
| $\hat{\sigma}$ | −1 | 261 | 24 | 340 |
| $r$ | −174 | −184 | −37 | −116 |
| $C$ | −247 | −706 | −72 | −613 |
| $\bar{\ell}$ | 1 | 2293 | 30 | 2350 |
| $Q$ | −195 | −221 | −68 | −146 |

#### Centrality rankings of disorders

Figure S9: For each data source and centrality measure, a correlation biplot of Kendall correlations between centrality rankings of disorders based on comorbidity networks constructed using the evidential cutoff $\alpha < 0.05$, each of three corrections (none, FWER, FDR), and each of five BAMs (unit, odds ratio, Pearson correlation, Forbes coefficient, and tetrachoric). In this and other biplots, first and second eigenvectors are reversed if necessary so that each centroid lies in the first quadrant.

A closer inspection of the effect of the TWER and of its corrections on node centralities (not shown) reveals that the largest inconsistencies in rank, at least in terms of degree and betweenness, tend to occur among the least central disorders. Thus, the highly central "hub" disorders are robust to the TWER. In contrast, the rankings suggest very different central cores in networks constructed from different data sources. The main text provides lists of the 8 most central disorders in each dataset, under betweenness centrality. Tables S12 and S13 provide analogous information based on degree and closeness centrality. Distance-based measures are more discriminating when graphs are more modular; because the stricter evidential cutoffs produced more modular graphs, we calculated centralities for these tables from the comorbidity networks based on 5% TWERs with FDR correction.

Table S12: Most degree-central disorders in each comorbidity network, subject to Benjamini-Hochberg-corrected 5% FDR.

| Dataset | Code | Description | Centrality |
|---|---|---|---|
| Columbia | | Tuberculosis | 111 |
| Columbia | | Gram-negative bacteria | 98 |
| Columbia | | Diabetes type 2 | 94 |
| Columbia | | Hypoosmolality | 93 |
| Columbia | | Benign neoplasms | 92 |
| Columbia | | Lipid metabolism d. | 92 |
| Columbia | | Carcinoma in situ | 89 |
| Columbia | | Hepatitis C | 89 |
| MedPAR(3) | 599 | Other disorders of urethra and urinary tract | 755 |
| MedPAR(3) | 276 | Disorders of fluid, electrolyte, and acid-base balance | 753 |
| MedPAR(3) | 041 | Bacterialinfectioninconditionsclassifiedelsewhereandofunspecifiedsite | 738 |
| MedPAR(3) | 285 | Other and unspecified anemias | 693 |
| MedPAR(3) | 038 | Septicemia | 680 |
| MedPAR(3) | 780 | General symptoms | 657 |
| MedPAR(3) | 263 | Other and unspecified protein-calorie malnutrition | 653 |
| MedPAR(3) | 787 | Symptoms involving digestive system | 648 |
| MedPAR(5) | 276.5 | Volume depletion | 3365 |
| MedPAR(5) | 285.9 | Anemia NOS | 3331 |
| MedPAR(5) | 276.1 | Hyposmolality | 2881 |
| MedPAR(5) | 276.8 | Hypopotassemia | 2790 |
| MedPAR(5) | 263.9 | Protein-cal malnutr NOS | 2574 |
| MedPAR(5) | 311 | Depressive disorder NEC | 2358 |
| MedPAR(5) | 298.9 | Psychosis NOS | 2112 |
| MedPAR(5) | 038.1 | Staphylococcal septicemia | 2110 |
| Sct.Hans | L30 | Other and unspecified dermatitis | 43 |
| Sct.Hans | R05 | Cough | 39 |
| Sct.Hans | R26 | Abnormalities of gait and mobility | 36 |
| Sct.Hans | R53 | Malaise and fatigue | 36 |
| Sct.Hans | R60 | Edema, not elsewhere classified | 35 |
| Sct.Hans | L29 | Pruritus | 34 |
| Sct.Hans | B17 | Other acute viral hepatitis | 30 |
| Sct.Hans | K77 | Liver disorders in diseases classified elsewhere | 28 |
| Michigan | 786.09 | Respiratory abnorm NEC | 7022 |
| Michigan | 729.5 | Pain in limb | 6845 |
| Michigan | 786.50 | Chest pain NOS | 6808 |
| Michigan | 789.00 | Abdmnal pain unspcf site | 6785 |
| Michigan | 786.2 | Cough | 6683 |
| Michigan | 486 | Pneumonia, organism NOS | 6661 |
| Michigan | 780.6 | Fever and other physiologic disturbances of temperature regulation | 6636 |
| Michigan | 427.9 | Cardiac dysrhythmia NOS | 6524 |
| Stanford | | Acidosis | 20 |
| Stanford | | Mineral met. (Ca) | 18 |
| Stanford | | Mineral met. (Mg) | 18 |
| Stanford | | Hyperosmolality | 17 |
| Stanford | | Gram-negative bacteria | 16 |
| Stanford | | Hypoosmolality | 16 |
| Stanford | | Urea cycle met. | 16 |
| Stanford | | Staphilococcus | 15 |
| Columbia* | | Tuberculosis | 26 |
| Columbia* | | Hepatitis C | 25 |
| Columbia* | | Benign neoplasms | 24 |
| Columbia* | | Diabetes type 2 | 24 |
| Columbia* | | Aplastic anemia | 22 |
| Columbia* | | Carcinoma in situ | 22 |
| Columbia* | | Cardiomyopathy | 22 |
| Columbia* | | Lipid metabolism d. | 22 |
| Columbia* | | Rheumatoid arthritis | 22 |
| Columbia* | | Virus | 22 |
| MIMIC | 5849 | Acute kidney failure NOS | 763 |
| MIMIC | 51881 | Acute respiratry failure | 668 |
| MIMIC | 99592 | Severe sepsis | 659 |
| MIMIC | 5990 | Urin tract infection NOS | 636 |
| MIMIC | 2762 | Acidosis | 624 |
| MIMIC | 2851 | Ac posthemorrhag anemia | 546 |
| MIMIC | 4280 | CHF NOS | 541 |
| MIMIC | 78552 | Septic shock | 533 |

Table S13: Most closeness-central disorders in each comorbidity network, subject to Benjamini-Hochberg-corrected 5% FDR.

| Dataset | Code | Description | Centrality |
|---|---|---|---|
| Columbia | | Tuberculosis | 0.434 |
| Columbia | | Gram-negative bacteria | 0.418 |
| Columbia | | Hypoosmolality | 0.414 |
| Columbia | | Diabetes type 2 | 0.413 |
| Columbia | | Benign neoplasms | 0.412 |
| Columbia | | Lipid metabolism d. | 0.412 |
| Columbia | | Carcinoma in situ | 0.409 |
| Columbia | | Hepatitis C | 0.409 |
| MedPAR(3) | 599 | Other disorders of urethra and urinary tract | 0.309 |
| MedPAR(3) | 276 | Disorders of fluid, electrolyte, and acid-base balance | 0.309 |
| MedPAR(3) | 041 | Bacterialinfectioninconditionsclassifiedelsewhereandofunspecifiedsite | 0.307 |
| MedPAR(3) | 285 | Other and unspecified anemias | 0.303 |
| MedPAR(3) | 038 | Septicemia | 0.302 |
| MedPAR(3) | 780 | General symptoms | 0.299 |
| MedPAR(3) | 263 | Other and unspecified protein-calorie malnutrition | 0.299 |
| MedPAR(3) | 787 | Symptoms involving digestive system | 0.299 |
| MedPAR(5) | 276.5 | Volume depletion | 0.000221 |
| MedPAR(5) | 285.9 | Anemia NOS | 0.000221 |
| MedPAR(5) | 276.1 | Hyposmolality | 0.000221 |
| MedPAR(5) | 276.8 | Hypopotassemia | 0.000221 |
| MedPAR(5) | 263.9 | Protein-cal malnutr NOS | 0.000221 |
| MedPAR(5) | 311 | Depressive disorder NEC | 0.000221 |
| MedPAR(5) | 298.9 | Psychosis NOS | 0.000221 |
| MedPAR(5) | 038.1 | Staphylococcal septicemia | 0.000221 |
| Sct.Hans | R60 | Edema, not elsewhere classified | 0.0452 |
| Sct.Hans | R53 | Malaise and fatigue | 0.0451 |
| Sct.Hans | R26 | Abnormalities of gait and mobility | 0.0451 |
| Sct.Hans | L30 | Other and unspecified dermatitis | 0.045 |
| Sct.Hans | R12 | Heartburn | 0.0449 |
| Sct.Hans | N30 | Cystitis | 0.0449 |
| Sct.Hans | B17 | Other acute viral hepatitis | 0.0449 |
| Sct.Hans | R50 | Fever of other and unknown origin | 0.0448 |
| Michigan | 786.09 | Respiratory abnorm NEC | 0.323 |
| Michigan | 729.5 | Pain in limb | 0.321 |
| Michigan | 786.50 | Chest pain NOS | 0.321 |
| Michigan | 789.00 | Abdmnal pain unspcf site | 0.32 |
| Michigan | 786.2 | Cough | 0.319 |
| Michigan | 486 | Pneumonia, organism NOS | 0.319 |
| Michigan | 780.6 | Fever and other physiologic disturbances of temperature regulation | 0.319 |
| Michigan | 427.9 | Cardiac dysrhythmia NOS | 0.317 |
| Stanford | | Vitamin deficiency | 0.0137 |
| Stanford | | Bundle branch block | 0.0137 |
| Stanford | | Lipid metabolism d. | 0.0137 |
| Stanford | | Giant cell arteritis | 0.0137 |
| Stanford | | Diabetes type 2 | 0.0137 |
| Stanford | | Keratoderma | 0.0137 |
| Stanford | | Rheumatoid arthritis | 0.0136 |
| Stanford | | Spondylolisthesis | 0.0136 |
| Columbia* | | Tuberculosis | 0.0143 |
| Columbia* | | Benign neoplasms | 0.0143 |
| Columbia* | | Aplastic anemia | 0.0143 |
| Columbia* | | Virus | 0.0143 |
| Columbia* | | Hepatitis B | 0.0143 |
| Columbia* | | Systemic lupus erythematosus | 0.0143 |
| Columbia* | | Depression | 0.0143 |
| Columbia* | | Helicobacter pilori | 0.0143 |
| MIMIC | 5849 | Acute kidney failure NOS | 0.000499 |
| MIMIC | 51881 | Acute respiratry failure | 0.000499 |
| MIMIC | 2762 | Acidosis | 0.000499 |
| MIMIC | 2851 | Ac posthemorrhag anemia | 0.000499 |
| MIMIC | 5990 | Urin tract infection NOS | 0.000499 |
| MIMIC | 99592 | Severe sepsis | 0.000499 |
| MIMIC | 5845 | Ac kidny fail, tubr necr | 0.000499 |
| MIMIC | 486 | Pneumonia, organism NOS | 0.000499 |

A legitimate concern with centrality analysis is that high centrality is only possible via many links, which may only be discernible due to high prevalence. That is, if two disorders have the same centrality in a "ground truth" network of all population-level comorbidities, the evidential cutoff makes the one with higher prevalence more likely to be centrally located in the comorbidity network (Table S14). While we found a clear positive relationship between prevalence and centrality, and a certain minimum prevalence is required for a certain centrality, neither is strongly predictive of the other (not shown).

Table S14: Most prevalent disorders in each comorbidity network.

| Dataset | Code | Description | Prevalence |
|---|---|---|---|
| Columbia | | Virus | 135833 |
| Columbia | | Prion | 78783 |
| Columbia | | Benign neoplasms | 77272 |
| Columbia | | Tuberculosis | 66569 |
| Columbia | | Diabetes type 2 | 60815 |
| Columbia | | Lipid metabolism d. | 57308 |
| Columbia | | Carcinoma in situ | 31151 |
| Columbia | | Streptococcus | 27682 |
| MedPAR(3) | 401 | Essential hypertension | 4417651 |
| MedPAR(3) | 276 | Disorders of fluid, electrolyte, and acid-base balance | 3542548 |
| MedPAR(3) | 427 | Cardiac dysrhythmias | 3138602 |
| MedPAR(3) | 414 | Other forms of chronic ischemic heart disease | 3110840 |
| MedPAR(3) | 599 | Other disorders of urethra and urinary tract | 2404493 |
| MedPAR(3) | 428 | Heart failure | 2397390 |
| MedPAR(3) | 250 | Diabetes mellitus | 2127097 |
| MedPAR(3) | 285 | Other and unspecified anemias | 1936774 |
| MedPAR(5) | 401.9 | Hypertension NOS | 4163381 |
| MedPAR(5) | 414.0 | Coronary atherosclerosis | 2673957 |
| MedPAR(5) | 428.0 | CHF NOS | 2343516 |
| MedPAR(5) | 599.0 | Urin tract infection NOS | 2087236 |
| MedPAR(5) | 496 | Chr airway obstruct NEC | 1878976 |
| MedPAR(5) | 427.31 | Atrial fibrillation | 1806969 |
| MedPAR(5) | 276.5 | Volume depletion | 1533833 |
| MedPAR(5) | 276.8 | Hypopotassemia | 1448586 |
| Sct.Hans | F20 | Schizophrenia | 1414 |
| Sct.Hans | F10 | Alcohol related disorders | 1149 |
| Sct.Hans | G47 | Sleep disorders | 990 |
| Sct.Hans | R44 | Oth symptoms and signs w general sensations and perceptions | 901 |
| Sct.Hans | R05 | Cough | 852 |
| Sct.Hans | J10 | Influenza due to other identified influenza virus | 756 |
| Sct.Hans | F12 | Cannabis related disorders | 665 |
| Sct.Hans | M54 | Dorsalgia | 654 |
| Michigan | 465.9 | Acute uri NOS | 171406 |
| Michigan | V70.0 | Routine medical exam | 149380 |
| Michigan | 427.9 | Cardiac dysrhythmia NOS | 148237 |
| Michigan | 786.50 | Chest pain NOS | 143866 |
| Michigan | V07.8 | Prophyl or tx meas NEC | 135571 |
| Michigan | 789.00 | Abdmnal pain unspcf site | 120539 |
| Michigan | 401.9 | Hypertension NOS | 117392 |
| Michigan | V20.2 | Routin child health exam | 114429 |
| Stanford | | Lipid metabolism d. | 79104 |
| Stanford | | Benign neoplasms | 59938 |
| Stanford | | Diabetes type 2 | 40176 |
| Stanford | | Carcinoma in situ | 27861 |
| Stanford | | Allergic rhinitis | 22523 |
| Stanford | | Bundle branch block | 13641 |
| Stanford | | Migraine | 12593 |
| Stanford | | Hypoosmolality | 12588 |
| Columbia* | | Virus | 135833 |
| Columbia* | | Benign neoplasms | 77272 |
| Columbia* | | Tuberculosis | 66569 |
| Columbia* | | Diabetes type 2 | 60815 |
| Columbia* | | Lipid metabolism d. | 57308 |
| Columbia* | | Carcinoma in situ | 31151 |
| Columbia* | | Streptococcus | 27682 |
| Columbia* | | Alcoholism | 27638 |
| MIMIC | 4019 | Hypertension NOS | 17613 |
| MIMIC | 41401 | Crnry athrscl natve vssl | 10775 |
| MIMIC | 42731 | Atrial fibrillation | 10271 |
| MIMIC | 4280 | CHF NOS | 9843 |
| MIMIC | 5849 | Acute kidney failure NOS | 7687 |
| MIMIC | 2724 | Hyperlipidemia NEC/NOS | 7465 |
| MIMIC | 25000 | DMII wo cmp nt st uncntr | 7370 |
| MIMIC | 51881 | Acute respiratry failure | 6719 |

Figure S10: Scatterplots of prevalences of disorders in the ICD9 ontology in different data sets using this ontology. Note the effect of the restriction, in the Michigan dataset, to disorders that appeared on at least 30 patient records.

Figure S11: Scatterplots of prevalences of disorders in the Rzhetsky ontology in different data sets using this ontology. Note that the disorders included in the Columbia dataset exactly match their prevalences in the Columbia dataset.

Among networks constructed from the three data sets that used the full level-5 ICD9 ontology, centrality rankings were discordant (Figures S12, S13, and S14): While those of MedPAR(5) and Michigan were moderately, though positively, correlated, that of MIMIC-III was anti-correlated with both, and highly so with MedPAR(5). The underlying patient populations were quite different in these cases, though (Medicare beneficiaries, a regional adult population, and ICU patients, respectively), which may entirely account for these differences. The node centrality rankings from the three data sets that used the Rzhetsky ontology (Columbia, Stanford, and Columbia*) were also moderately concordant by each of the three centrality measures.

Based on the mappings to level-3 ICD9 codes and to the Rzhetsky ontology, the group centrality rankings for these data sets become more concordant: more correlated between MedPAR(5) and Michigan and less anti-correlated between both and MIMIC. We see only very weak concordance between any of the group centrality rankings and the node centrality rankings, under either mapping. We include tables of most-central disorders for each ontology in Tables S16 through S17.

Figure S12: For each p-value correction and centrality measure, a correlation biplot of Kendall correlations between centrality rankings of disorders based on comorbidity networks constructed from each data set that uses or could be crosswalked to the Rzhetsky ontology. Centrality measures are node-based for data encoded using this ontology and group-based for data crosswalked to this ontology.

Figure S13: For each p-value correction and centrality measure, a correlation biplot of Kendall correlations between centrality rankings of disorders based on comorbidity networks constructed from each data set that uses or could be crosswalked to the level-3 ICD9 ontology. Centrality measures are node-based for data encoded using this ontology and group-based for data crosswalked to this ontology.

Figure S14: For each p-value correction and centrality measure, a correlation biplot of Kendall correlations between centrality rankings of disorders based on comorbidity networks constructed from each data set that uses the level-5 ICD9 ontology.

Table S15: Most degree-central disorders in comorbidity networks using the Rzhetsky ontology, subject to Benjamini-Hochberg-corrected 5% FDR.

| Dataset | Description | Centrality |
|---|---|---|
| Columbia | Tuberculosis | 111 |
| Columbia | Gram-negative bacteria | 98 |
| Columbia | Diabetes type 2 | 94 |
| Columbia | Hypoosmolality | 93 |
| Columbia | Benign neoplasms | 92 |
| Columbia | Lipid metabolism d. | 92 |
| Columbia | Carcinoma in situ | 89 |
| Columbia | Hepatitis C | 89 |
| Columbia | Streptococcus | 88 |
| Columbia | Diabetes type 1 | 87 |
| Columbia | Helicobacter pilori | 87 |
| MedPAR(5) | Hypoosmolality | 2881 |
| MedPAR(5) | Gram-negative bacteria (infection) | 2863 |
| MedPAR(5) | Diabetes mellitus type 2 | 2251 |
| MedPAR(5) | Benign neoplasms | 2250 |
| MedPAR(5) | Diabetes mellitus type 1 | 2178 |
| MedPAR(5) | Carcinoma in situ | 2102 |
| MedPAR(5) | Alzheimer's disease | 1879 |
| MedPAR(5) | Staphilococcus | 1746 |
| MedPAR(5) | Streptococcus | 1682 |
| MedPAR(5) | Alcoholism | 1615 |
| Michigan | Benign neoplasms | 5251 |
| Michigan | Carcinoma in situ | 4856 |
| Michigan | Lipid metabolism disorders | 4803 |
| Michigan | Diabetes mellitus type 2 | 4501 |
| Michigan | Allergic rhinitis | 4296 |
| Michigan | Cholelithiasis | 3638 |
| Michigan | Migraine | 3517 |
| Michigan | Diabetes mellitus type 1 | 3429 |
| Michigan | Virus | 3419 |
| Michigan | Depression | 3409 |
| Stanford | Acidosis | 20 |
| Stanford | Mineral met. (Ca) | 18 |
| Stanford | Mineral met. (Mg) | 18 |
| Stanford | Hyperosmolality | 17 |
| Stanford | Gram-negative bacteria | 16 |
| Stanford | Hypoosmolality | 16 |
| Stanford | Urea cycle met. | 16 |
| Stanford | Staphilococcus | 15 |
| Stanford | Alkalosis | 14 |
| Stanford | Epilepsy | 14 |
| Columbia* | Tuberculosis | 26 |
| Columbia* | Hepatitis C | 25 |
| Columbia* | Benign neoplasms | 24 |
| Columbia* | Diabetes type 2 | 24 |
| Columbia* | Aplastic anemia | 22 |
| Columbia* | Carcinoma in situ | 22 |
| Columbia* | Cardiomyopathy | 22 |
| Columbia* | Lipid metabolism d. | 22 |
| Columbia* | Rheumatoid arthritis | 22 |
| Columbia* | Virus | 22 |
| MIMIC | HIV | 102 |
| MIMIC | Multiple sclerosis | 37 |
| MIMIC | Depression | 25 |
| MIMIC | Benign neoplasms | 8 |
| MIMIC | Virus | 5 |
| MIMIC | Polyarteritis nodosa | 2 |

Table S16: Most degree-central disorders in comorbidity networks using the ICD9-3 ontology, subject to Benjamini-Hochberg-corrected 5% FDR.

| Dataset | Code | Description | Centrality |
|---|---|---|---|
| MedPAR(3) | 599 | Other disorders of urethra and urinary tract | 755 |
| MedPAR(3) | 276 | Disorders of fluid, electrolyte, and acid-base balance | 753 |
| MedPAR(3) | 041 | Bacterialinfectioninconditionsclassifiedelsewhereandofunspecifiedsite | 738 |
| MedPAR(3) | 285 | Other and unspecified anemias | 693 |
| MedPAR(3) | 038 | Septicemia | 680 |
| MedPAR(3) | 780 | General symptoms | 657 |
| MedPAR(3) | 263 | Other and unspecified protein-calorie malnutrition | 653 |
| MedPAR(3) | 787 | Symptoms involving digestive system | 648 |
| MedPAR(3) | 112 | Candidiasis | 645 |
| MedPAR(3) | 707 | Chronic ulcer of skin | 634 |
| MedPAR(3) | 486 | Pneumonia, organism NOS | 629 |
| MedPAR(3) | 518 | Other diseases of lung | 628 |
| MedPAR(5) | 311 | Depressive disorder NEC | 2358 |
| MedPAR(5) | 486 | Pneumonia, organism NOS | 1106 |
| MedPAR(5) | 261 | Nutritional marasmus | 1030 |
| MedPAR(5) | 496 | Chr airway obstruct NEC | 920 |
| MedPAR(5) | 262 | Oth severe malnutrition | 736 |
| MedPAR(5) | 260 | Kwashiorkor | 733 |
| MedPAR(5) | 438 | Late effects of cerebrovascular disease | 708 |
| MedPAR(5) | 436 | Cva | 650 |
| MedPAR(5) | 585 | Chronic kidney disease (CKD) | 564 |
| MedPAR(5) | 586 | Renal failure NOS | 485 |
| MedPAR(5) | 515 | Postinflam pulm fibrosis | 467 |
| MedPAR(5) | 319 | Intellect disability NOS | 463 |
| Michigan | 486 | Pneumonia, organism NOS | 6661 |
| Michigan | 311 | Depressive disorder NEC | 5729 |
| Michigan | 462 | Acute pharyngitis | 3932 |
| Michigan | 591 | Hydronephrosis | 3780 |
| Michigan | 496 | Chr airway obstruct NEC | 3702 |
| Michigan | 514 | Pulm congest/hypostasis | 3471 |
| Michigan | 490 | Bronchitis NOS | 3458 |
| Michigan | 586 | Renal failure NOS | 3436 |
| Michigan | 436 | Cva | 3344 |
| Michigan | 920 | Contusion face/scalp/nck | 3025 |
| Michigan | 217 | Benign neoplasm breast | 2837 |
| Michigan | 585 | Chronic kidney disease (CKD) | 2713 |
| MIMIC | 276 | Disorders of fluid, electrolyte, and acid-base balance | 990 |
| MIMIC | 285 | Other and unspecified anemias | 959 |
| MIMIC | 518 | Other diseases of lung | 904 |
| MIMIC | E87 | | 903 |
| MIMIC | 584 | Acute kidney failure | 849 |
| MIMIC | V58 | Encounter for other and unspecified procedures and aftercare | 807 |
| MIMIC | 995 | Certain adverse effects not elsewhere classified | 768 |
| MIMIC | 996 | Complications peculiar to certain specified procedures | 754 |
| MIMIC | E93 | | 734 |
| MIMIC | V45 | Other postprocedural states | 720 |
| MIMIC | 785 | Symptoms involving cardiovascular system | 698 |
| MIMIC | 041 | Bacterialinfectioninconditionsclassifiedelsewhereandofunspecifiedsite | 691 |

Table S17: Most closeness-central disorders in comorbidity networks using the Rzhetsky ontology, subject to Benjamini-Hochberg-corrected 5% FDR.

| Dataset | Description | Centrality |
|---|---|---:|
| Columbia | Tuberculosis | 0.434 |
| Columbia | Gram-negative bacteria | 0.418 |
| Columbia | Hypoosmolality | 0.414 |
| Columbia | Diabetes type 2 | 0.413 |
| Columbia | Benign neoplasms | 0.412 |
| Columbia | Lipid metabolism d. | 0.412 |
| Columbia | Carcinoma in situ | 0.409 |
| Columbia | Hepatitis C | 0.409 |
| Columbia | Streptococcus | 0.407 |
| Columbia | Helicobacter pilori | 0.407 |
| MedPAR(5) | Hypoosmolality | 0.000221 |
| MedPAR(5) | Hyperosmolality | 0.000221 |
| MedPAR(5) | Alzheimer's disease | 0.000221 |
| MedPAR(5) | Acidosis | 0.000221 |
| MedPAR(5) | Alkalosis | 0.000221 |
| MedPAR(5) | Gram-negative bacteria (infection) | 0.000221 |
| MedPAR(5) | Mineral metabolism disorders (Ca) | 0.000221 |
| MedPAR(5) | Mineral metabolism disorders (Mg) | 0.000221 |
| MedPAR(5) | Depression | 0.000221 |
| MedPAR(5) | Parkinson's disease | 0.000221 |
| Michigan | Hypoosmolality | 0.283 |
| Michigan | Cardiomyopathy, primary | 0.28 |
| Michigan | Acidosis | 0.278 |
| Michigan | Alkalosis | 0.276 |
| Michigan | Systemic lupus erythematosus | 0.274 |
| Michigan | Depression | 0.272 |
| Michigan | Allergic rhinitis | 0.272 |
| Michigan | Keratoderma | 0.271 |
| Michigan | Psoriasis | 0.27 |
| Michigan | Hyperosmolality | 0.27 |
| Stanford | Vitamin deficiency | 0.0137 |
| Stanford | Bundle branch block | 0.0137 |
| Stanford | Lipid metabolism d. | 0.0137 |
| Stanford | Giant cell arteritis | 0.0137 |
| Stanford | Diabetes type 2 | 0.0137 |
| Stanford | Keratoderma | 0.0137 |
| Stanford | Rheumatoid arthritis | 0.0136 |
| Stanford | Spondylolisthesis | 0.0136 |
| Stanford | Alzheimer's | 0.0136 |
| Stanford | Cholelithiasis | 0.0136 |
| Columbia* | Tuberculosis | 0.0143 |
| Columbia* | Benign neoplasms | 0.0143 |
| Columbia* | Aplastic anemia | 0.0143 |
| Columbia* | Virus | 0.0143 |
| Columbia* | Hepatitis B | 0.0143 |
| Columbia* | Systemic lupus erythematosus | 0.0143 |
| Columbia* | Depression | 0.0143 |
| Columbia* | Helicobacter pilori | 0.0143 |
| Columbia* | Alopecia | 0.0143 |
| Columbia* | Bipolar disorder | 0.0143 |
| Columbia* | Psoriasis | 0.0143 |
| MIMIC | HIV | 0.000499 |
| MIMIC | Multiple sclerosis | 0.000499 |
| MIMIC | Polyarteritis nodosa | 0.000499 |
| MIMIC | Depression | 0.000314 |
| MIMIC | Benign neoplasms | 0.000296 |
| MIMIC | Virus | 0.000245 |
| MIMIC | Streptococcus | 0.000163 |

Table S18: Most closeness-central disorders in comorbidity networks using the ICD9-3 ontology, subject to Benjamini-Hochberg-corrected 5% FDR.

| Dataset | Code | Description | Centrality |
|---|---|---|---|
| MedPAR(3) | 599 | Other disorders of urethra and urinary tract | 0.309 |
| MedPAR(3) | 276 | Disorders of fluid, electrolyte, and acid-base balance | 0.309 |
| MedPAR(3) | 041 | Bacterialinfectioninconditionsclassifiedelsewhereandofunspecifiedsite | 0.307 |
| MedPAR(3) | 285 | Other and unspecified anemias | 0.303 |
| MedPAR(3) | 038 | Septicemia | 0.302 |
| MedPAR(3) | 780 | General symptoms | 0.299 |
| MedPAR(3) | 263 | Other and unspecified protein-calorie malnutrition | 0.299 |
| MedPAR(3) | 787 | Symptoms involving digestive system | 0.299 |
| MedPAR(3) | 112 | Candidiasis | 0.298 |
| MedPAR(3) | 707 | Chronic ulcer of skin | 0.297 |
| MedPAR(3) | 486 | Pneumonia, organism NOS | 0.297 |
| MedPAR(3) | 518 | Other diseases of lung | 0.297 |
| MedPAR(5) | 311 | Depressive disorder NEC | 0.000221 |
| MedPAR(5) | 486 | Pneumonia, organism NOS | 0.000221 |
| MedPAR(5) | 438 | Late effects of cerebrovascular disease | 0.000221 |
| MedPAR(5) | 496 | Chr airway obstruct NEC | 0.000221 |
| MedPAR(5) | 436 | Cva | 0.000221 |
| MedPAR(5) | 261 | Nutritional marasmus | 0.000221 |
| MedPAR(5) | 585 | Chronic kidney disease (CKD) | 0.000221 |
| MedPAR(5) | 586 | Renal failure NOS | 0.000221 |
| MedPAR(5) | 260 | Kwashiorkor | 0.000221 |
| MedPAR(5) | 262 | Oth severe malnutrition | 0.000221 |
| MedPAR(5) | 490 | Bronchitis NOS | 0.000221 |
| MedPAR(5) | 515 | Postinflam pulm fibrosis | 0.000221 |
| Michigan | 486 | Pneumonia, organism NOS | 0.319 |
| Michigan | 311 | Depressive disorder NEC | 0.308 |
| Michigan | 462 | Acute pharyngitis | 0.29 |
| Michigan | 591 | Hydronephrosis | 0.289 |
| Michigan | 496 | Chr airway obstruct NEC | 0.288 |
| Michigan | 514 | Pulm congest/hypostasis | 0.286 |
| Michigan | 490 | Bronchitis NOS | 0.286 |
| Michigan | 586 | Renal failure NOS | 0.286 |
| Michigan | 436 | Cva | 0.285 |
| Michigan | 920 | Contusion face/scalp/nck | 0.282 |
| Michigan | 217 | Benign neoplasm breast | 0.28 |
| Michigan | 412 | Old myocardial infarct | 0.279 |
| MIMIC | 486 | Pneumonia, organism NOS | 0.000499 |
| MIMIC | 311 | Depressive disorder NEC | 0.000499 |
| MIMIC | 496 | Chr airway obstruct NEC | 0.000499 |
| MIMIC | 276 | Disorders of fluid, electrolyte, and acid-base balance | 0.000499 |
| MIMIC | 570 | Acute necrosis of liver | 0.000499 |
| MIMIC | 412 | Old myocardial infarct | 0.000499 |
| MIMIC | 578 | Gastrointestinal hemorrhage | 0.000499 |
| MIMIC | 285 | Other and unspecified anemias | 0.000499 |
| MIMIC | 261 | Nutritional marasmus | 0.000499 |
| MIMIC | 591 | Hydronephrosis | 0.000499 |
| MIMIC | 401 | Essential hypertension | 0.000499 |
| MIMIC | 403 | Hypertensive chronic kidney disease | 0.000499 |

#### Additional results from the multivariate analysis

##### Correlation structure and link determination for NAMCS encounters

Figure S16 compares the estimated disease correlations under the four models. The switch from the pairwise tetrachoric scheme to JDM0 shifted the correlation estimates negatively by a consistent magnitude, with no evident scaling. A linear model with fixed slope 1 estimated the shift at $-0.174$. The relationship between the pairwise and partial correlations is noisier, and is not clearly captured by a linear model with fixed intercept 0 or by one with fixed slope 1—that is, by shifting or by

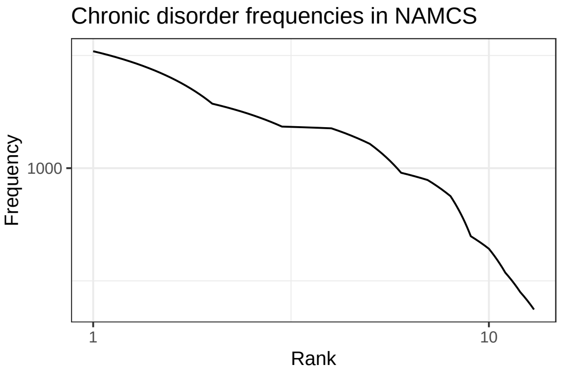


Figure S15: Frequency–rank plot for the 13 chronic disorders recorded in the NAMCS sample.

scaling. The correlation estimates in JDM1 are offset according as the exogenous effects were positive or negative and possibly scaled by a non-unit factor. A linear model with a binary term for the sign of the exogenous effect estimated the slope at .89.

The $18 \times 13 = 234$ effect estimates of the exogenous covariates $\hat{\mathrm{B}}$ on the prevalence of each chronic disease are reported in Table S19, and Figure S17 summarizes these estimates, together with the component of each of the 78 pairwise correlations accounted for by these exogenous effects. The effects on prevalence were distributed unimodally around zero with a standard deviation near $\frac{3}{5}$. The effects on co-occurrence $\hat{\mathbf{P}}$ were substantial, relative to the residual correlations, cleanly split between positive (49) and negative (29). The positive and negative effects were centered about the median values .86 and $-.55$, and the positive effects were themselves bimodally distributed.

Differences between the pairwise network on one hand, and the partial correlation and endogenous-only JDM networks on the other, were due to the loss of information on how the co-occurrences of disorder pairs were concentrated in certain subsets of patients when moving from the patient–disorder occurrence data to the disorder–disorder contingency tables (endogenous covariance). Meanwhile, the discrepancy between the residual networks of the two JDMs, primarily the weakening or disappearance of both positive and negative correlations, indicates that these correlations were due in part to differences in co-occurrence across specific demographic groups (exogenous covariance). Each systems model (partial correlation or JDM) tended to reduce comorbidity estimates, relative to the conventional model, but there were rich disagreements among these three, especially between the partial correlation network and the JDMs on whether several disease pairs were positively versus negatively associated (Figure S18).

##### Correlation structure and link determination for MIMIC-III units

The scatterplots in Figure S21 compare correlation estimates for each pair of disorders in the CSRU data, which are representative of the others (excluding the neonatal units). As with NAMCS, and consistent with expectations, the correlation estimates on average weakened (or turned negative) from the pairwise model to the other models. However, this trend was much more modest than with the chronic disorders recorded in NAMCS.

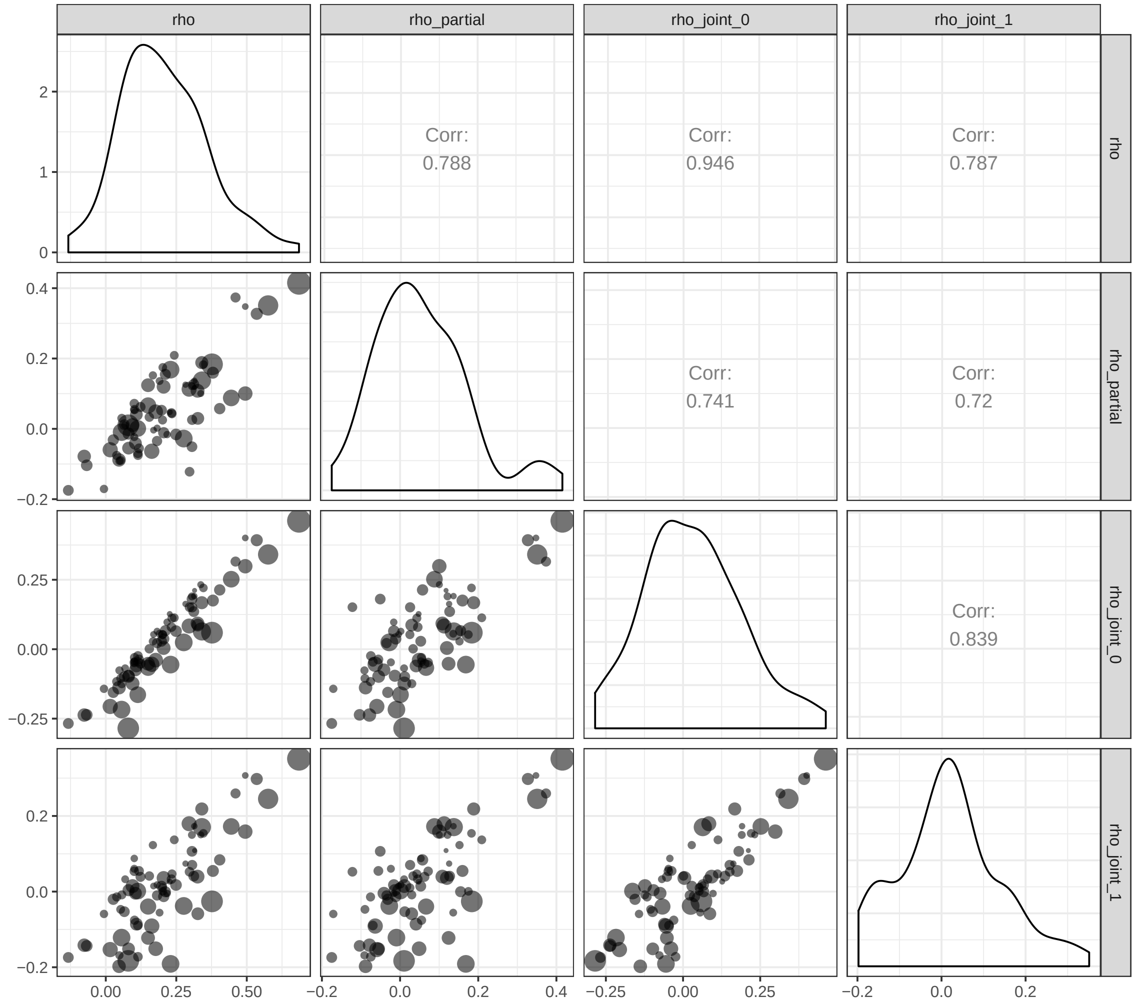


Figure S16: Scatterplots between the pairwise tetrachoric correlations $r_t$, the full partial correlations $r_t'$, and the correlation point estimates $\hat{\rho}_0$ and $\hat{\rho}_1$ from the JDMs for the NAMCS population. The area of each point is in proportion to the product $n_1 n_2$ of the prevalences of the two disorders—equivalently, to their expected co-occurrence under the null hypothesis of no comorbidity.

Bagley SC, Sirota M, Chen R et al (2016) Constraints on Biological Mechanism from Disease Comorbidity Using Electronic Medical Records and Database of Genetic Variants. PLoS Comput Biol 12:e1004885. doi: 10.1371/journal.pcbi.1004885

Broido AD, Clauset A (2018) Scale-free networks are rare

Burnham KP, Anderson DR (2004) Multimodel Inference. Sociological Methods & Research 33:261–304. doi: 10.1177/0049124104268644

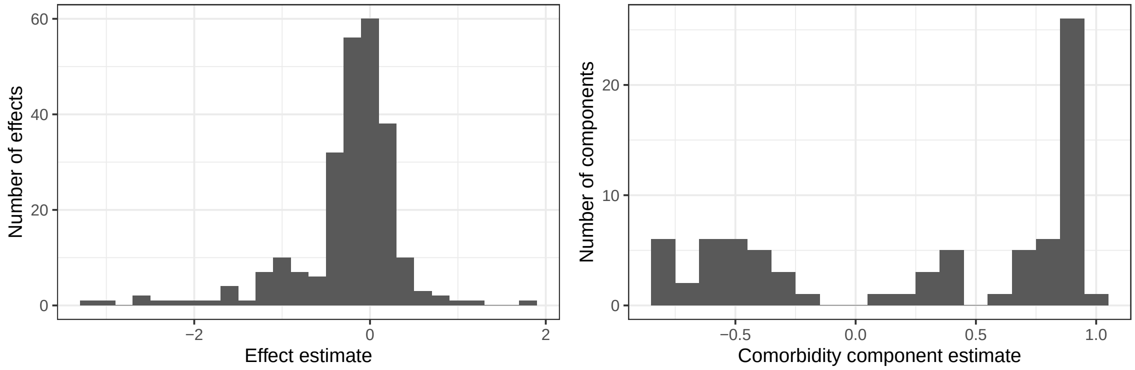


Figure S17: Histograms of estimates from the joint distribution model with exogenous (patient-level) predictors. Left: Exogenous effects on disorder prevalence. Right: Correlation (epidemiological comorbidity) accounted for by exogenous effects.

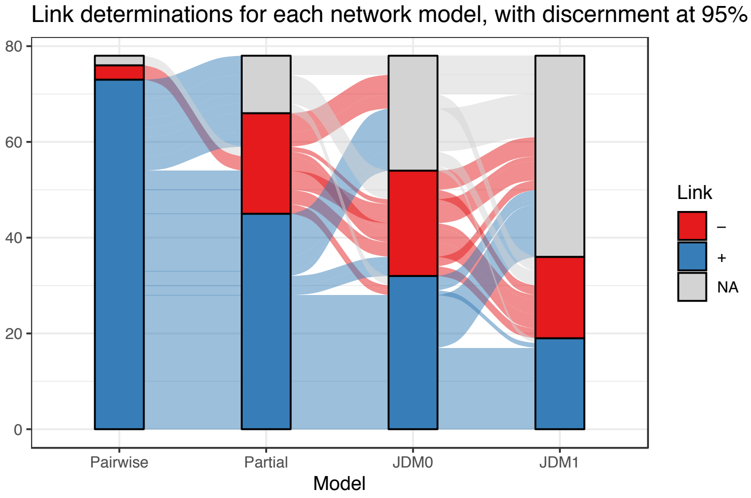


Figure S18: Alluvial diagram of discernible signs of the population-level associations between NAMCS chronic disorders, using each of 4 network models: pairwise correlation, full partial correlation, endogenous joint distribution, and joint distribution controlling for exogenous predictors.

| predictor | arthritis | asthma | cancer | CVD | CHF | COPD | depression | diabetes | HLD | HT | IHD | obesity | OP |
|---|---|---|---|---|---|---|---|---|---|---|---|---|---|
| Const. | – | -1.09 | -1.13 | -1.62 | -1.05 | -0.98 | -0.99 | – | – | 0.94 | -1.17 | -1.03 | -0.72 |
| 0-14 | -1.11 | 1.77 | -1.51 | -3.06 | -2.64 | – | – | -1.47 | -1.94 | -2.56 | -3.22 | 0.45 | -1.02 |
| 15-24 | -0.92 | 0.68 | -1.11 | -1.00 | -2.44 | -0.77 | 1.22 | -0.96 | -1.52 | -1.80 | -1.53 | 0.70 | -2.23 |
| 25-44 | -0.31 | 0.35 | -0.86 | -0.77 | -1.14 | -0.41 | 0.90 | -0.39 | -0.78 | -0.96 | -1.18 | 0.71 | -1.19 |
| 45-64 | – | 0.15 | -0.36 | -0.42 | -0.70 | – | 0.54 | – | -0.21 | -0.36 | -0.38 | 0.45 | -0.54 |
| 65-74 | – | – | – | -0.15 | -0.55 | – | – | – | – | – | -0.16 | 0.25 | – |
| M | -0.21 | -0.24 | 0.16 | – | 0.21 | – | -0.27 | 0.21 | 0.18 | 0.16 | 0.34 | -0.16 | -0.75 |
| Asian | -0.33 | -0.33 | -0.31 | – | -0.48 | – | -0.52 | – | – | – | -0.37 | -0.49 | – |
| Black | – | – | -0.35 | – | – | -0.51 | -0.48 | – | – | – | -0.34 | – | -0.47 |
| White | – | -0.33 | – | -0.39 | -0.51 | -0.38 | – | -0.55 | -0.36 | -0.47 | -0.34 | -0.35 | – |
| Hispanic | – | – | -0.29 | – | – | – | -0.16 | 0.44 | 0.18 | 0.16 | – | 0.15 | – |
| Private | – | 0.18 | 0.16 | 0.34 | – | – | -0.43 | – | 0.19 | 0.19 | 0.19 | – | – |
| Medicare | 0.15 | – | 0.15 | 0.39 | – | 0.26 | -0.18 | 0.16 | 0.20 | 0.24 | 0.32 | – | 0.25 |
| Medicaid | – | 0.14 | – | – | – | 0.28 | – | – | 0.13 | 0.18 | – | 0.21 | – |
| Midwest | -0.12 | – | – | – | – | – | – | – | -0.10 | -0.23 | -0.20 | -0.17 | -0.36 |
| South | -0.15 | -0.12 | -0.15 | – | – | – | 0.16 | – | -0.20 | -0.15 | -0.48 | -0.18 | -0.32 |
| West | – | – | – | -0.27 | – | – | – | – | – | -0.26 | -0.24 | – | – |
| Metro | – | – | 0.36 | 0.33 | – | -0.19 | 0.13 | -0.15 | – | -0.22 | 0.26 | – | – |

Table S19: Estimated effects of demographic predictors on the incidence of chronic disorders in Model 1. Each value indicates the effect of the predictor on the mean of the normal distribution from which the latent variable is sampled (see the text). Estimates whose 95% credible intervals contain zero are excluded.

Figure S19: For each of four models, a correlation biplot of estimated latent correlations between 13 chronic disorders in the NAMCS sample.

Figure S20: For each critical care unit in the MIMIC-III database, a frequency–rank plot for the recorded diagnoses, crosswalked to CCS codes.


Chen Y, Xu R (2014) Mining Cancer-Specific Disease Comorbidities from a Large Observational Health Database. Cancer Informatics 37–44. doi: 10.4137/CIN.S13893

Chmiel A, Klimek P, Thurner S (2014) Spreading of diseases through comorbidity networks across life and gender. New Journal of Physics 16:115013. doi: 10.1088/1367-2630/16/11/115013

Drasgow F (2006) Polychoric and Polyserial Correlations. In: Encyclopedia of statistical sciences. John Wiley & Sons, Inc., Hoboken, NJ, USA

Epskamp S, Fried EI (2017) A Tutorial on Regularized Partial Correlation Networks

Feldman K, Stiglic G, Dasgupta D et al (2016) Insights into Population Health Management Through Disease Diagnoses Networks. Sci Rep 6:30465. doi: 10.1038/srep30465

Folino F, Pizzuti C (2012) Link Prediction Approaches for Disease Networks. In: Böhm C, Khuri S, Lhotská L, Renda ME (eds) Information technology in bio- and medical informatics: Third international conference. Springer Berlin Heidelberg, Berlin, Heidelberg, pp 99–108

Gelman A, Hill J, Yajima M (2012) Why We (Usually) Don't Have to Worry About Multiple Comparisons. Journal of Research on Educational Effectiveness 5:189–211. doi: 10.1080/19345747.2011.618213

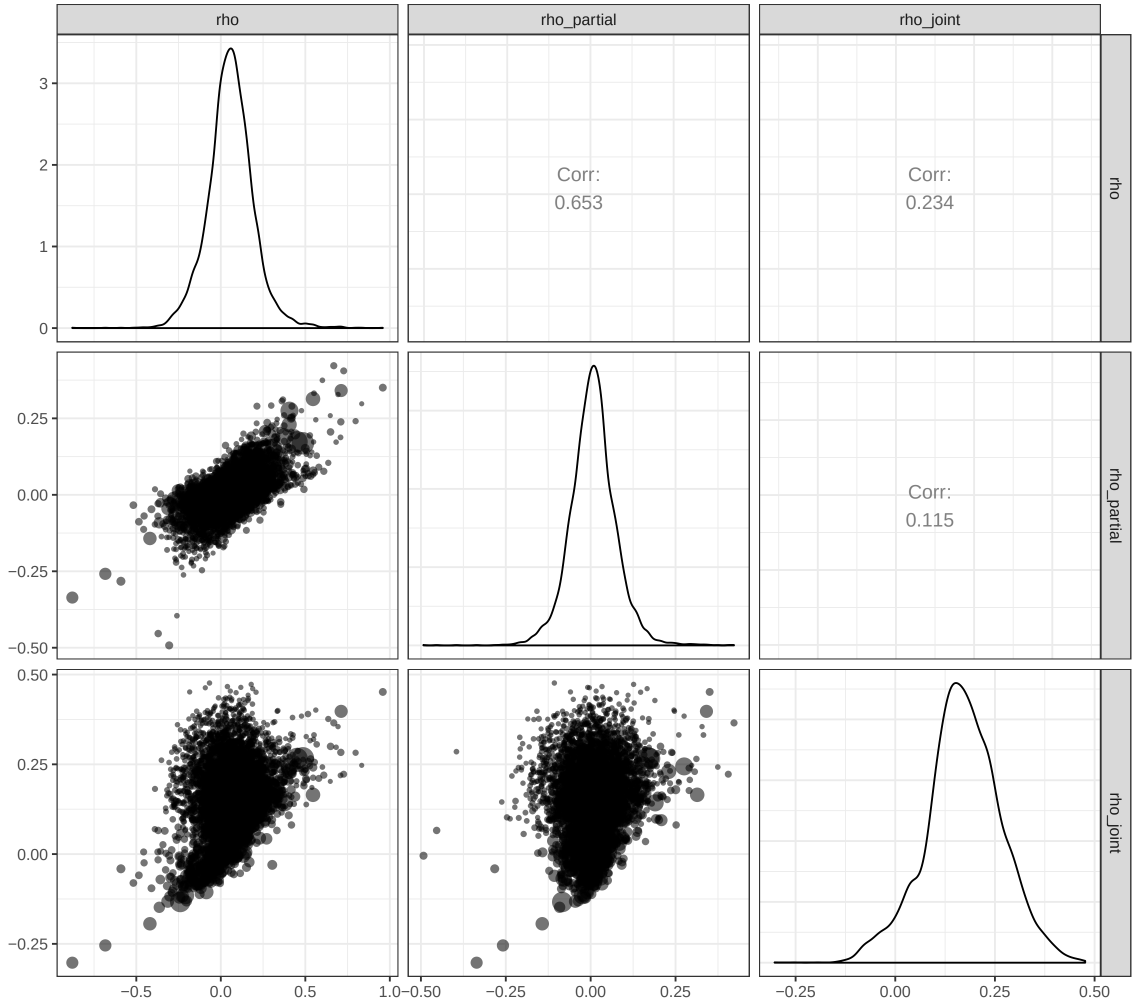


Figure S21: Scatterplots between the pairwise tetrachoric correlations $r_t$, the full partial correlations $r_t'$, and the correlation estimates $\hat{\rho}_0$ from the endogenous JDM for the CSRU population.

Glicksberg BS, Li L-2, Badgeley MA et al (2016) Comparative analyses of population-scale phenomic data in electronic medical records reveal race-specific disease networks. Bioinformatics 32:i101–i110. doi: 10.1093/bioinformatics/btw282

Hanauer DA, Ramakrishnan N (2013) Modeling temporal relationships in large scale clinical associations. Journal of the American Medical Informatics Association 20:332–341. doi: 10.1136/amiajnl-2012-001117

Hanauer DA, Rhodes DR, Chinnaiyan AM (2009) Exploring Clinical Associations Using '-Omics' Based Enrichment Analyses. PLoS One 4:e5203. doi: 10.1371/journal.pone.0005203

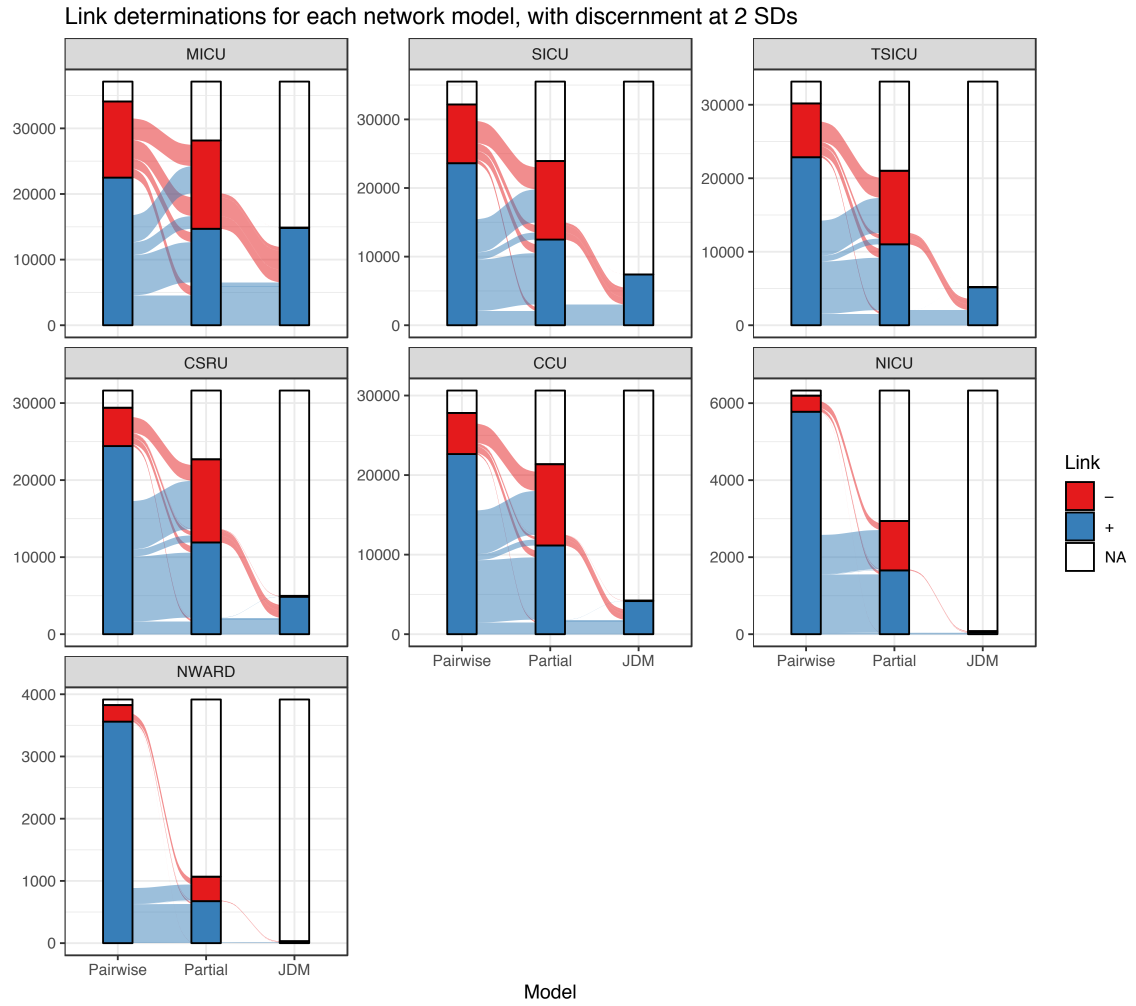


Figure S22: Alluvial diagrams of discernible signs of the population-level associations between MIMIC diagnoses within each admission unit cohort, using each of 3 network models: pairwise correlation, full partial correlation, and endogenous joint distribution.

Figure S23: Three comorbidity networks constructed from the CCU population. From left to right: Sample tetrachoric correlations $r_t$, partial tetrachoric correlations $r_t{}'$, and correlation estimates $\hat{\rho}_0$ from the endogenous JDM.

Hidalgo CA, Blumm N, Barabasi A-L, Christakis NA (2009) A Dynamic Network Approach for the Study of Human Phenotypes. PLoS Comput Biol 5:e1000353. doi: 10.1371/journal.pcbi.1000353

Hubalek Z (1982) Coefficients of association and similarity, based on binary (presence-absence) data:

Figure S24: Three comorbidity networks constructed from the CSRU population. From left to right: Sample tetrachoric correlations $r_t$, partial tetrachoric correlations ${r_t}'$, and correlation estimates $\hat{\rho}_0$ from the endogenous JDM.

Figure S25: Three comorbidity networks constructed from the MICU population. From left to right: Sample tetrachoric correlations $r_t$, partial tetrachoric correlations ${r_t}'$, and correlation estimates $\hat{\rho}_0$ from the endogenous JDM.

Figure S26: Three comorbidity networks constructed from the NICU population. From left to right: Sample tetrachoric correlations $r_t$, partial tetrachoric correlations ${r_t}'$, and correlation estimates $\hat{\rho}_0$ from the endogenous JDM.

Figure S27: Three comorbidity networks constructed from the NWARD population. From left to right: Sample tetrachoric correlations $r_t$, partial tetrachoric correlations ${r_t}'$, and correlation estimates $\hat{\rho}_0$ from the endogenous JDM.

Figure S28: Three comorbidity networks constructed from the SICU population. From left to right: Sample tetrachoric correlations $r_t$, partial tetrachoric correlations ${r_t}'$, and correlation estimates $\hat{\rho}_0$ from the endogenous JDM.

Figure S29: Three comorbidity networks constructed from the TSICU population. From left to right: Sample tetrachoric correlations $r_t$, partial tetrachoric correlations ${r_t}'$, and correlation estimates $\hat{\rho}_0$ from the endogenous JDM.

Figure S30: For each care unit and centrality measure, a correlation biplot of Kendall correlations between centrality rankings of disorders based on each of four comorbidity network models: pairwise correlations, full partial correlations, and a joint distribution model without exogenous effects.

an evaluation. Biological Reviews 57:669–689. doi: 10.1111/j.1469-185X.1982.tb00376.x

Jensen AB, Moseley PL, Oprea TI et al (2014) Temporal disease trajectories condensed from population-wide registry data covering 6.2 million patients. Nat Commun 5:4022. doi: 10.1038/ncomms5022

Kim JH, Son KY, Shin DW et al (2016) Network analysis of human diseases using Korean nationwide claims data. J Biomed Inform 61:276–282. doi: 10.1016/j.jbi.2016.05.002

Kraemer HC (1995) Statistical issues in assessing comorbidity. Statistics in Medicine 14:721–733. doi: 10.1002/sim.4780140803

Lai Y-H (2015) Network Analysis of Comorbidities: Case Study of HIV/AIDS in Taiwan. 2:174–186

Parzen M, Lipsitz S, Ibrahim J, Klar N (2002) An Estimate of the Odds Ratio That Always Exists. Journal of Computational and Graphical Statistics 11:420–436. doi: 10.1198/106186002760180590

Pollock LJ, Tingley R, Morris WK et al (2014) Understanding co-occurrence by modelling species simultaneously with a Joint Species Distribution Model (JSDM). Methods in Ecology and Evolution 5:397–406. doi: 10.1111/2041-210X.12180

Roque FS, Jensen PB, Schmock H et al (2011) Using electronic patient records to discover disease correlations and stratify patient cohorts. PLoS Comput Biol 7:e1002141. doi: 10.1371/journal.pcbi.1002141

Rzhetsky A, Wajngurt D, Park N, Zheng T (2007) Probing genetic overlap among complex human phenotypes. Proceedings of the National Academy of Sciences 104:11694–11699. doi: 10.1073/pnas.0704820104

Zhang J, Yu KF (1998) What's the Relative Risk? JAMA 280:1690. doi: 10.1001/jama.280.19.1690
