## Supplementary figures and images for "Sensitivity and robustness of comorbidity network analysis"

### Figure S1

Quantiles at which selected evaluative cutoffs prune comorbidity networks

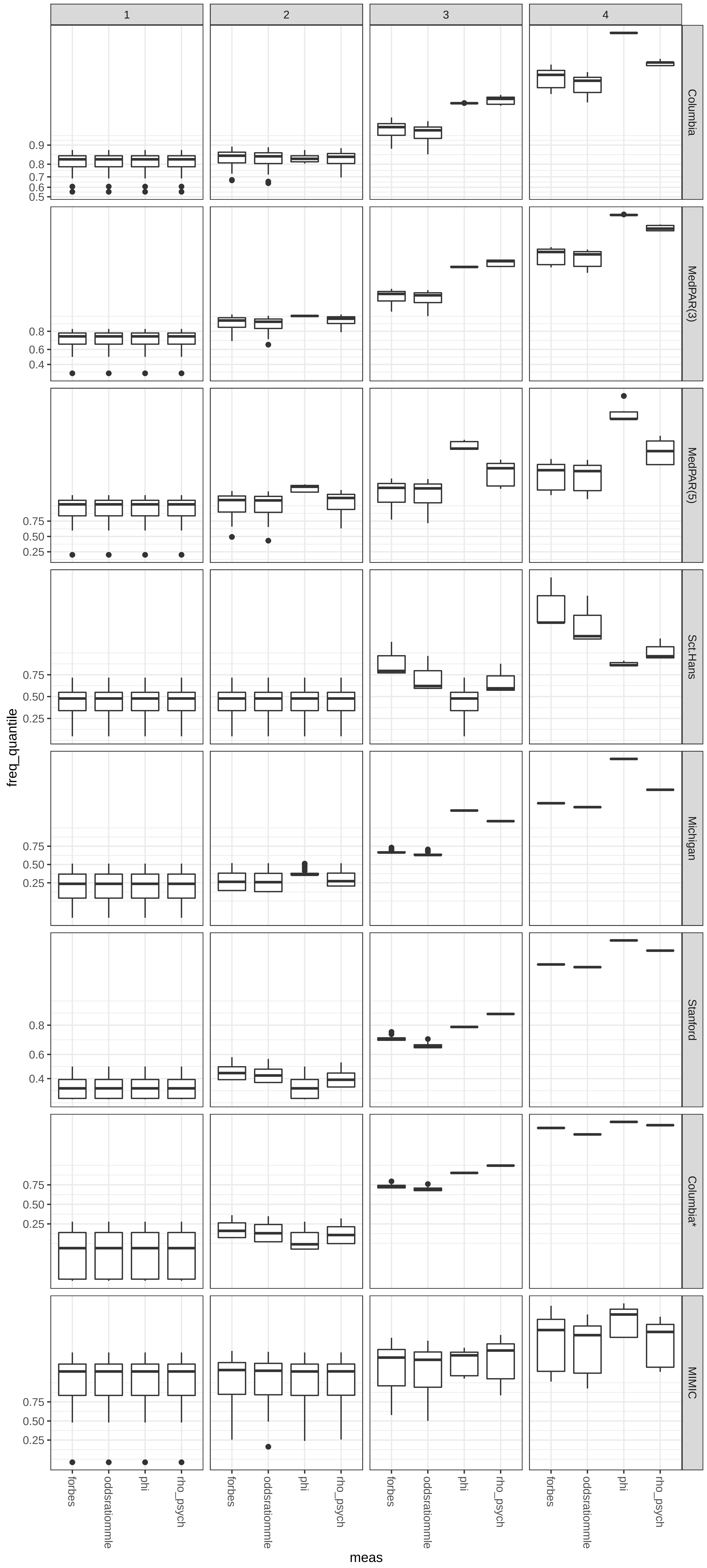

### Figure S2

#### Columbia network, pruned at FDR−corrected 5%

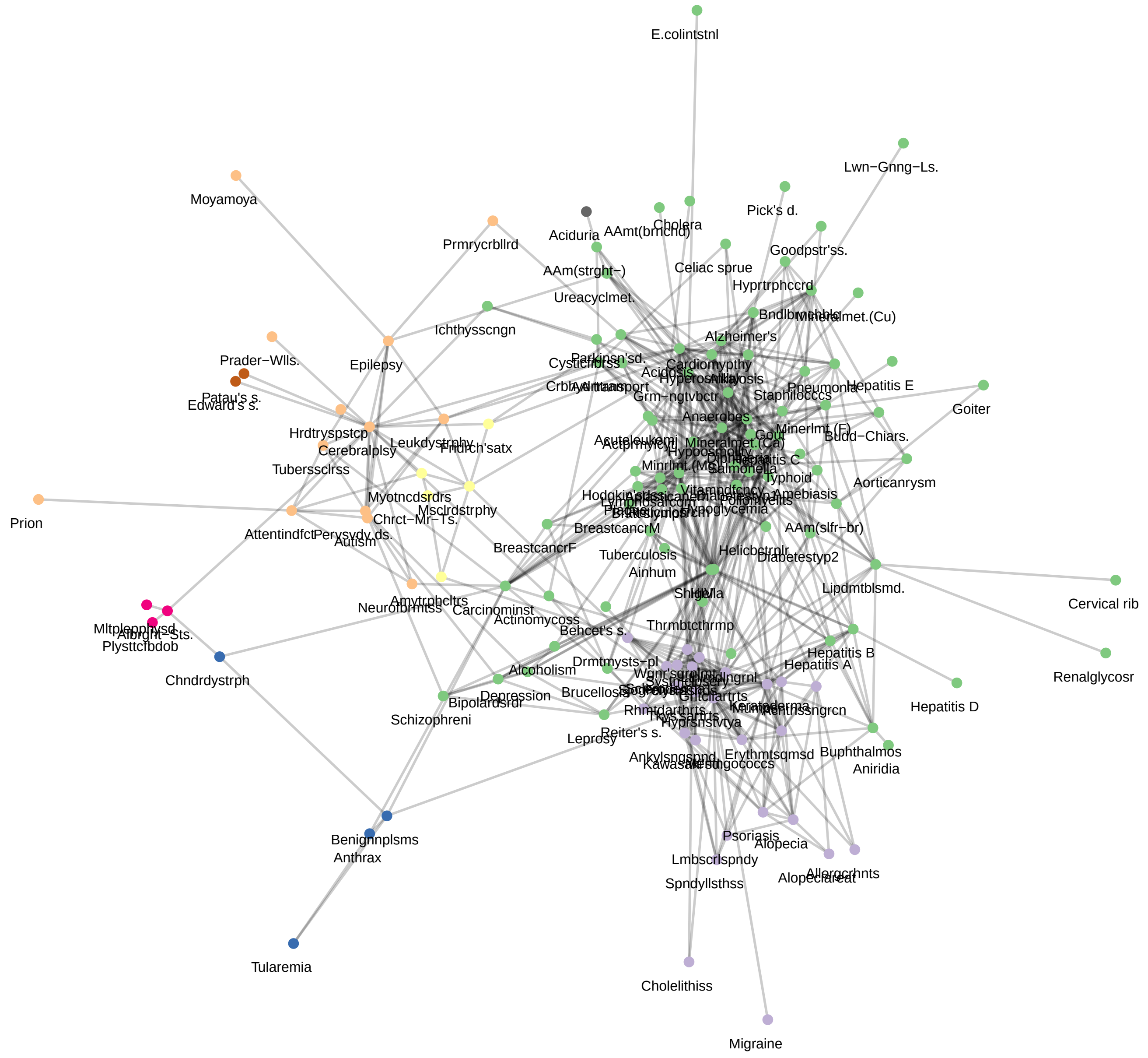

### Figure S3

#### MedPAR(3) network, pruned at FDR−corrected 5%

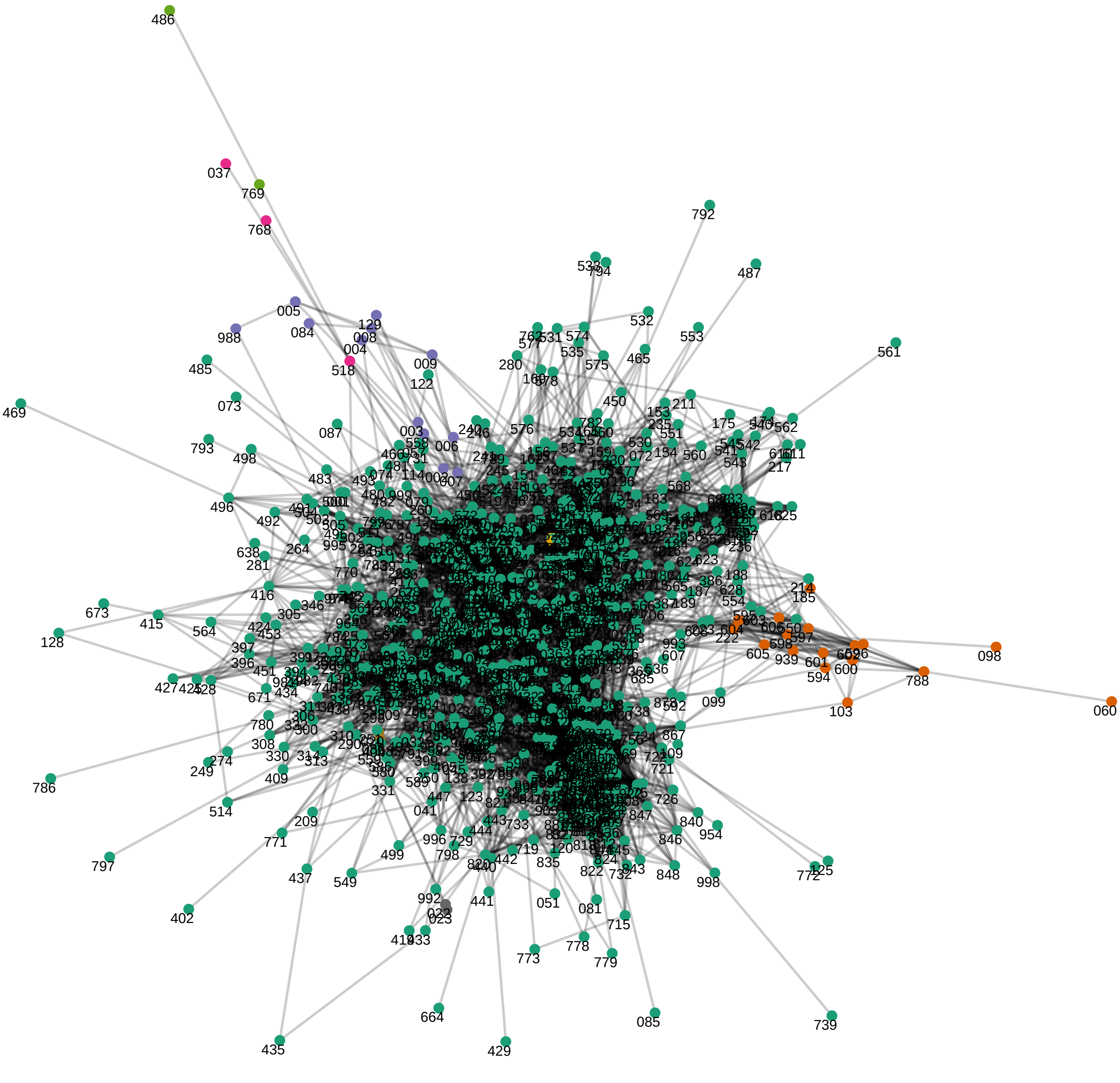

### Figure S4

#### Sct.Hans network, pruned at FDR−corrected 5%

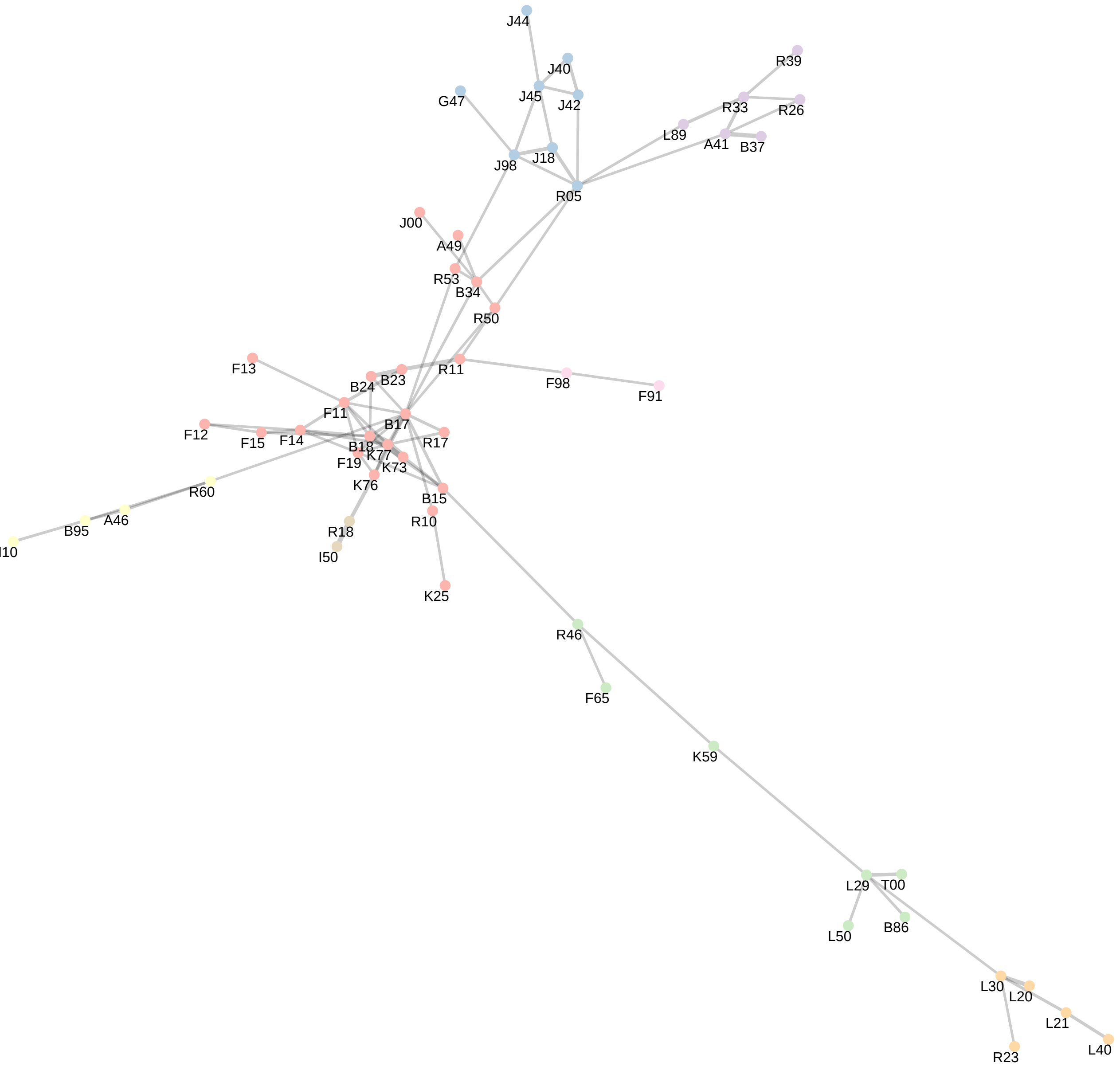

### Figure S5

#### Stanford network, pruned at FDR−corrected 5%

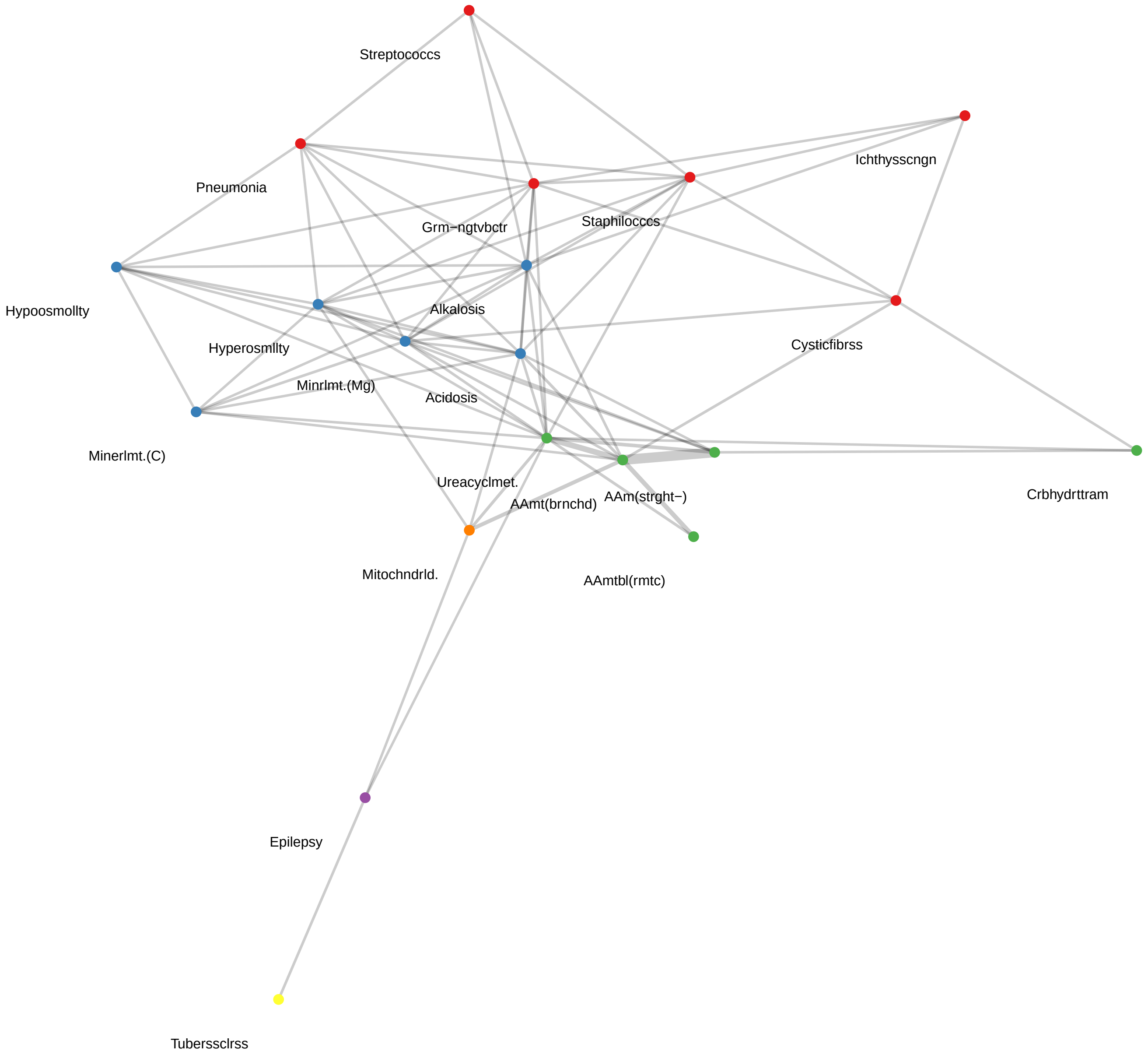

### Figure S6

#### Columbia* network, pruned at FDR−corrected 5%

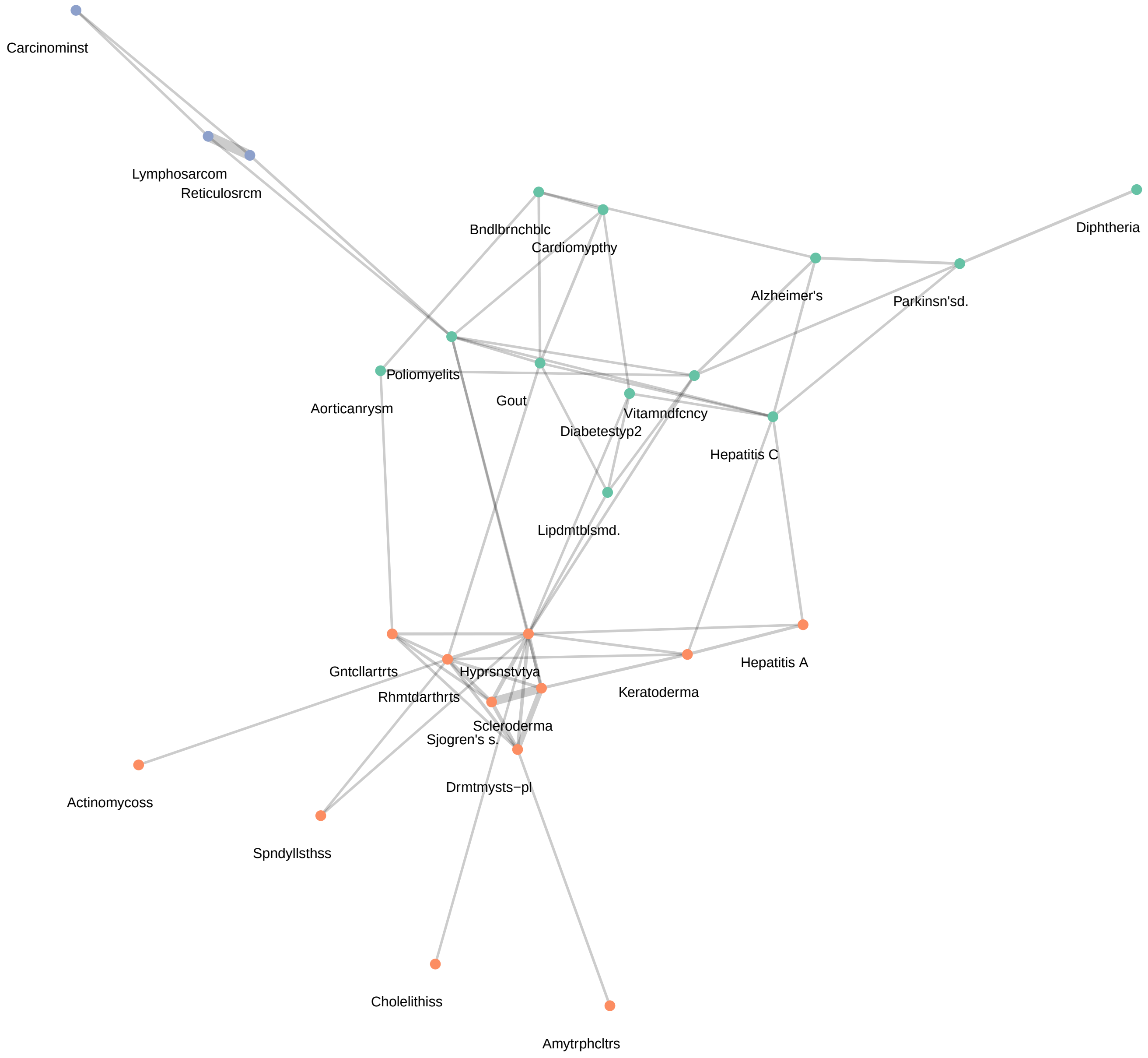

### Figure S7

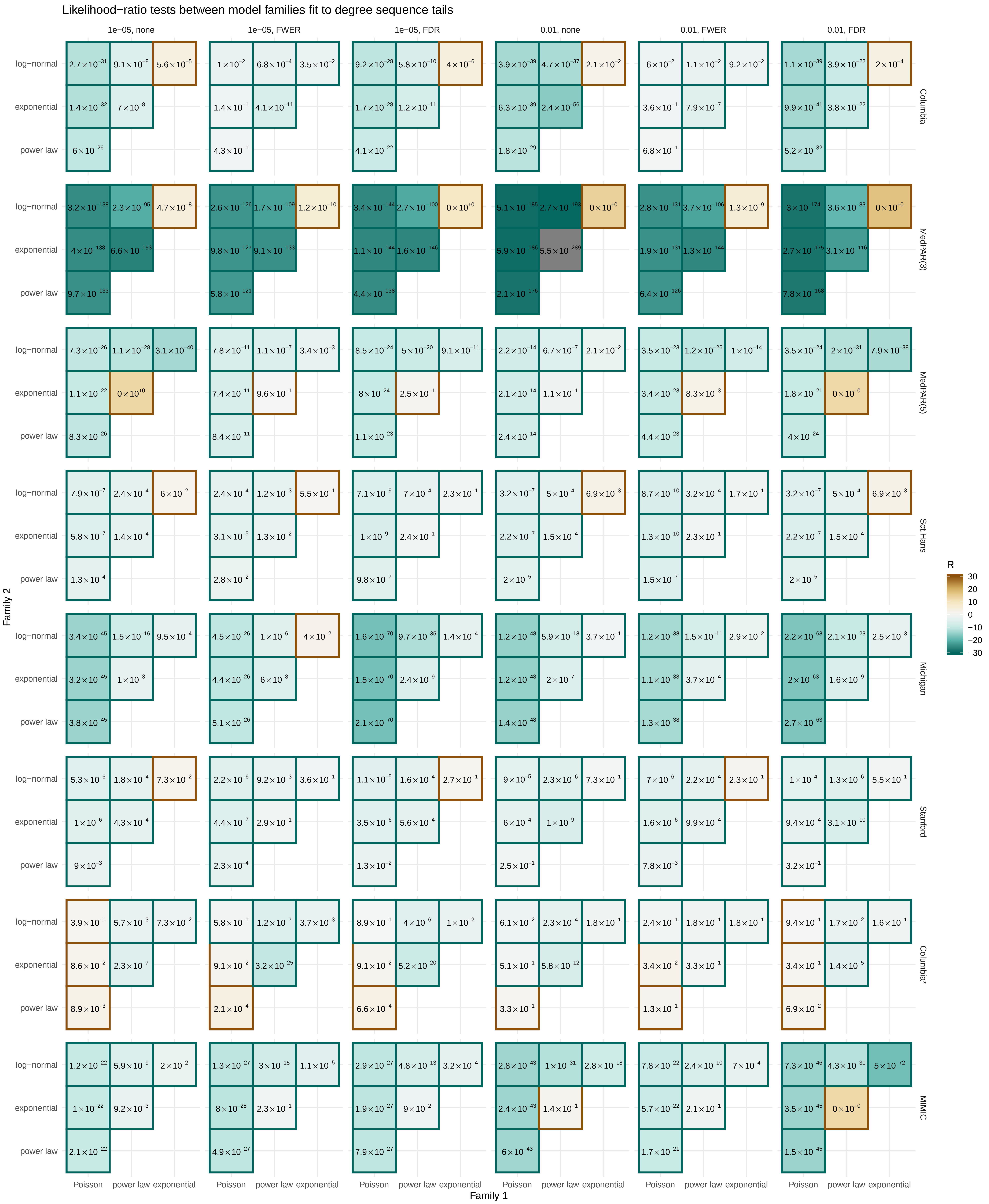

### Figure S8

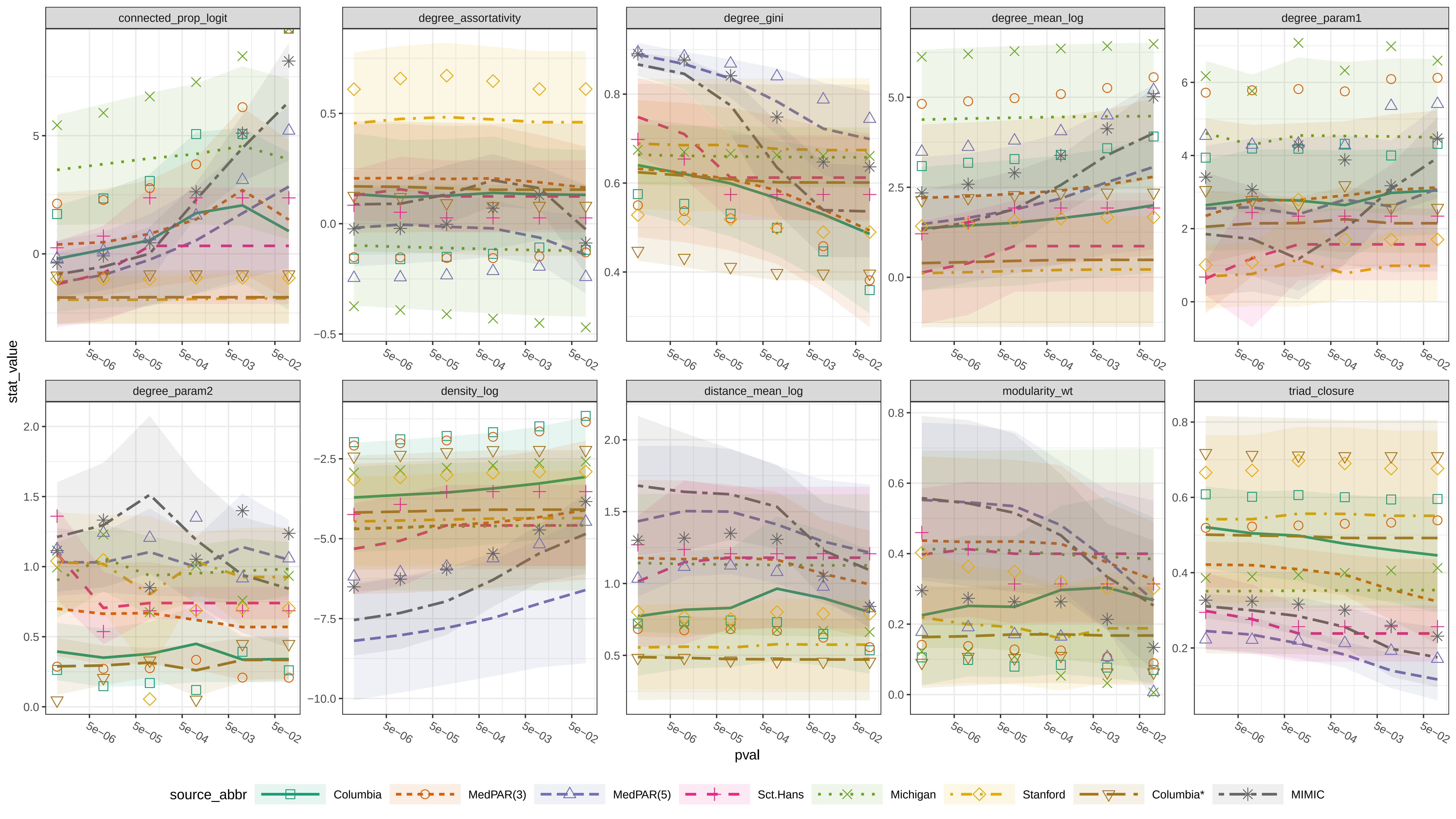

### Figure S9

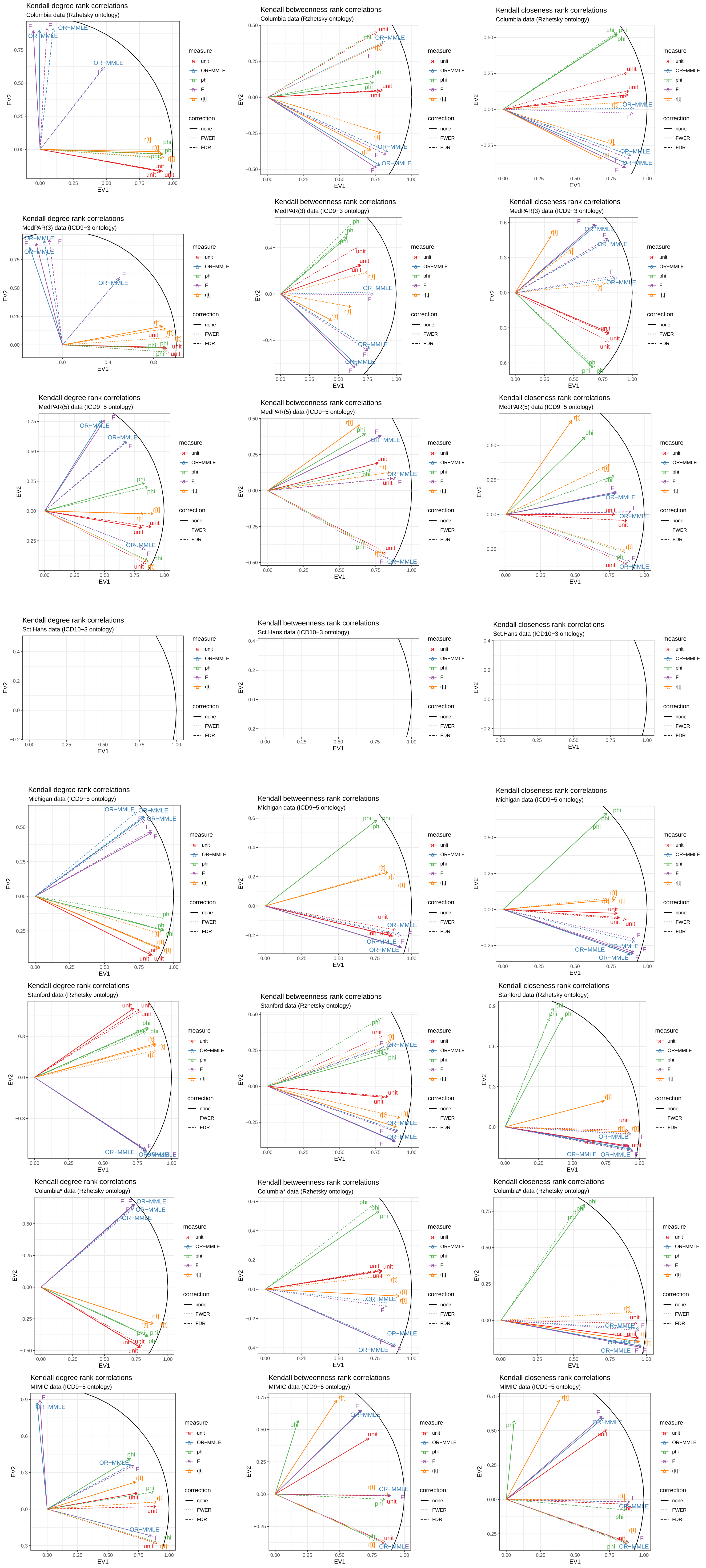

### Figure S10

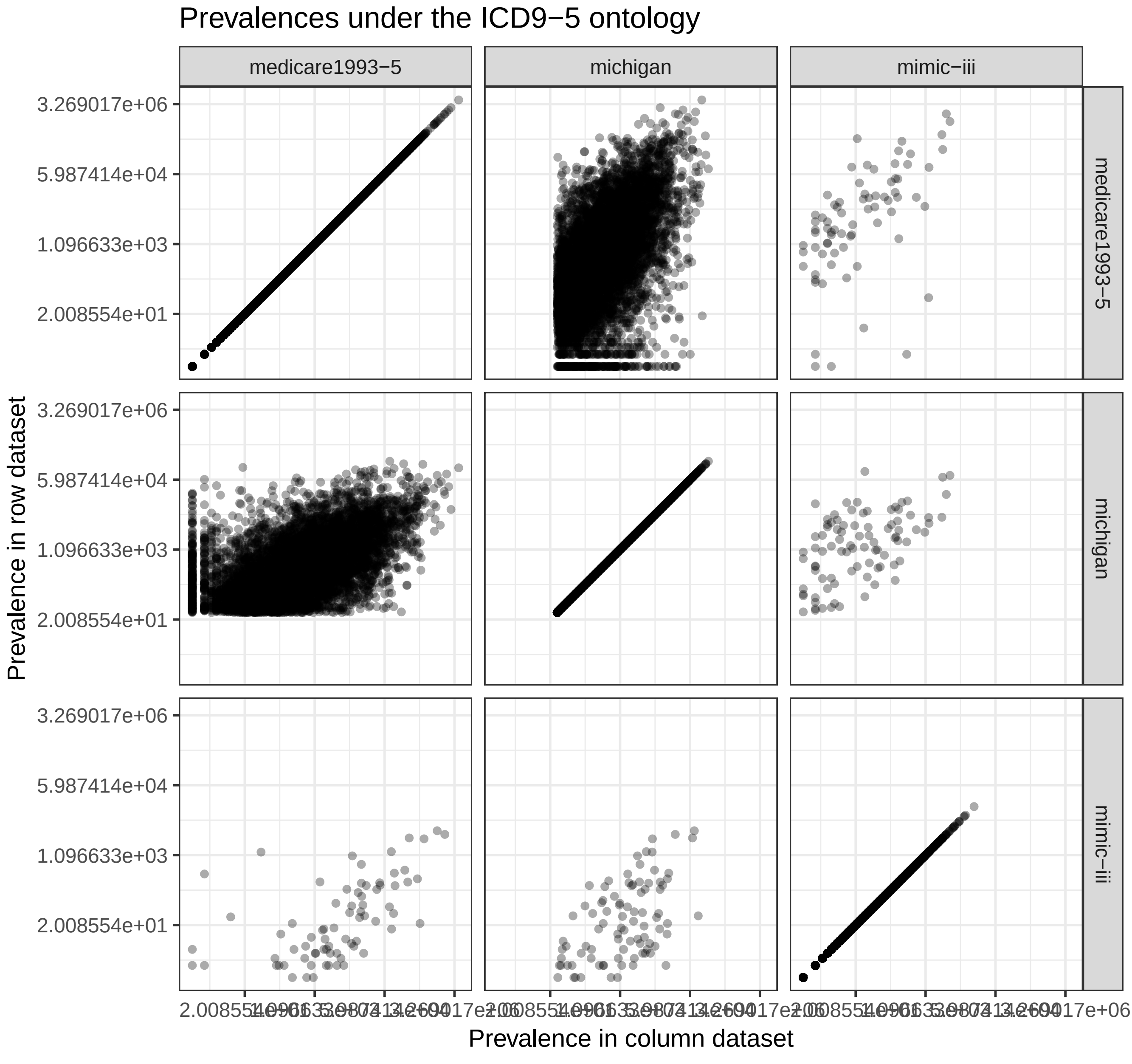

### Figure S11

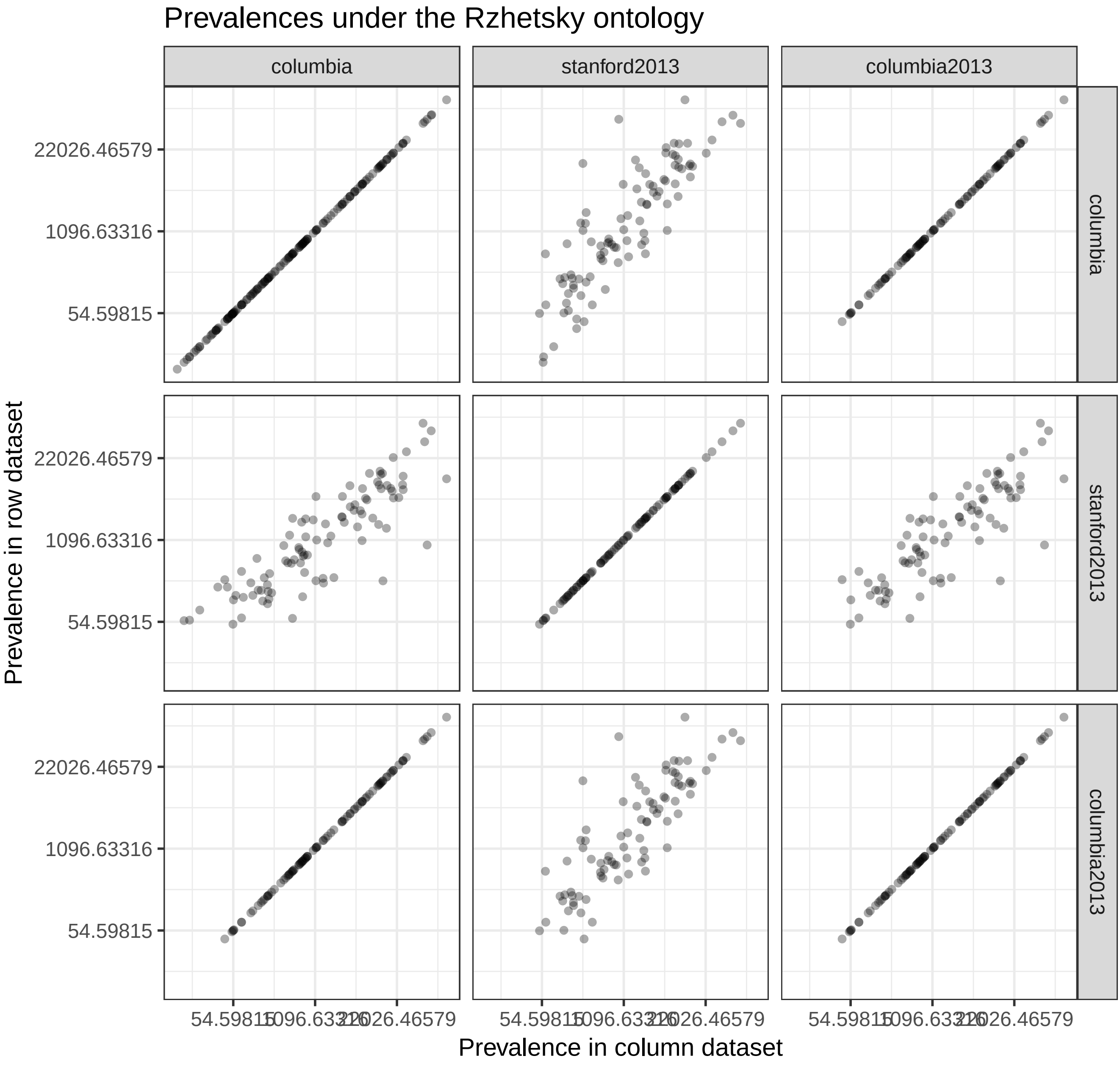

### Figure S12

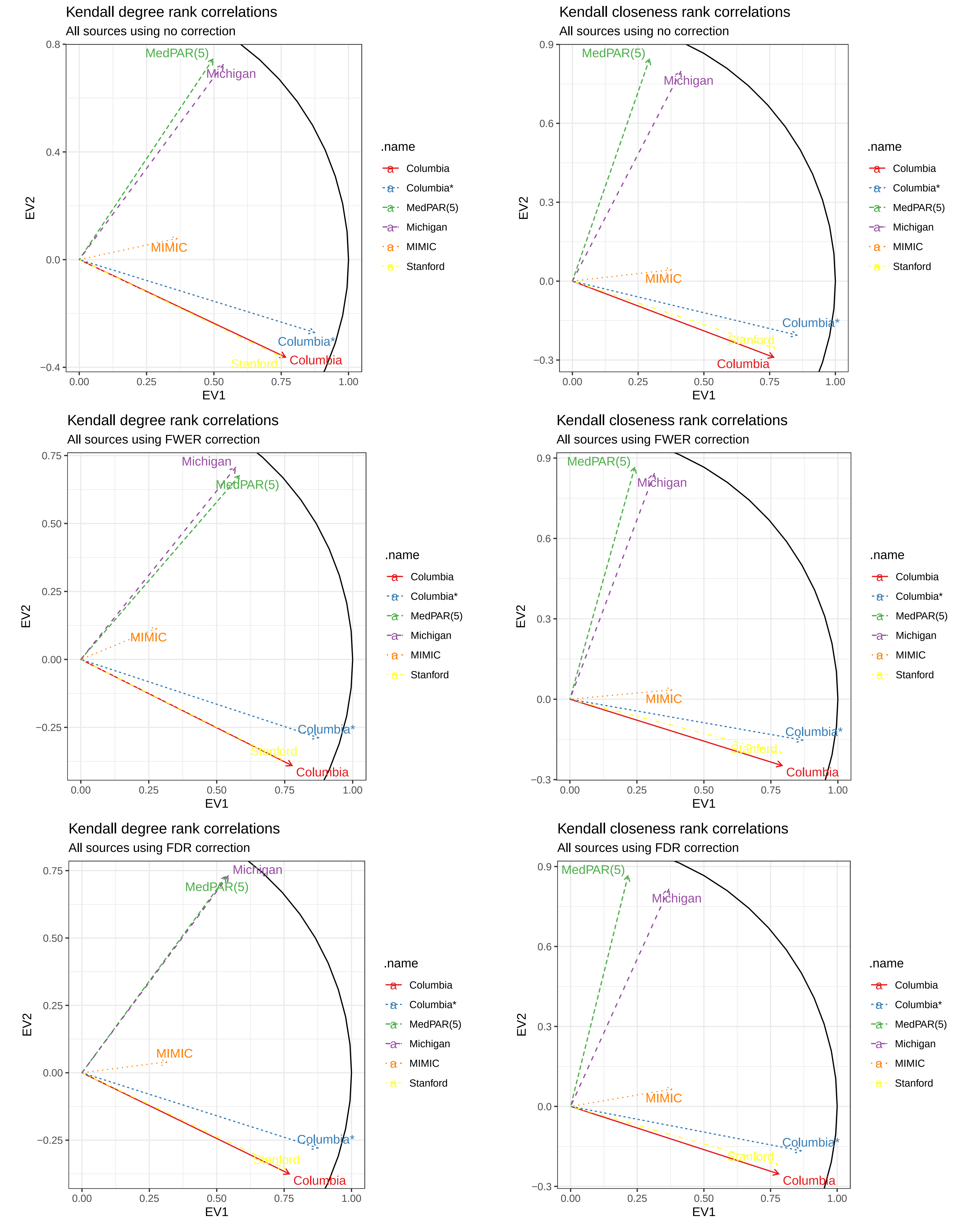

### Figure S14

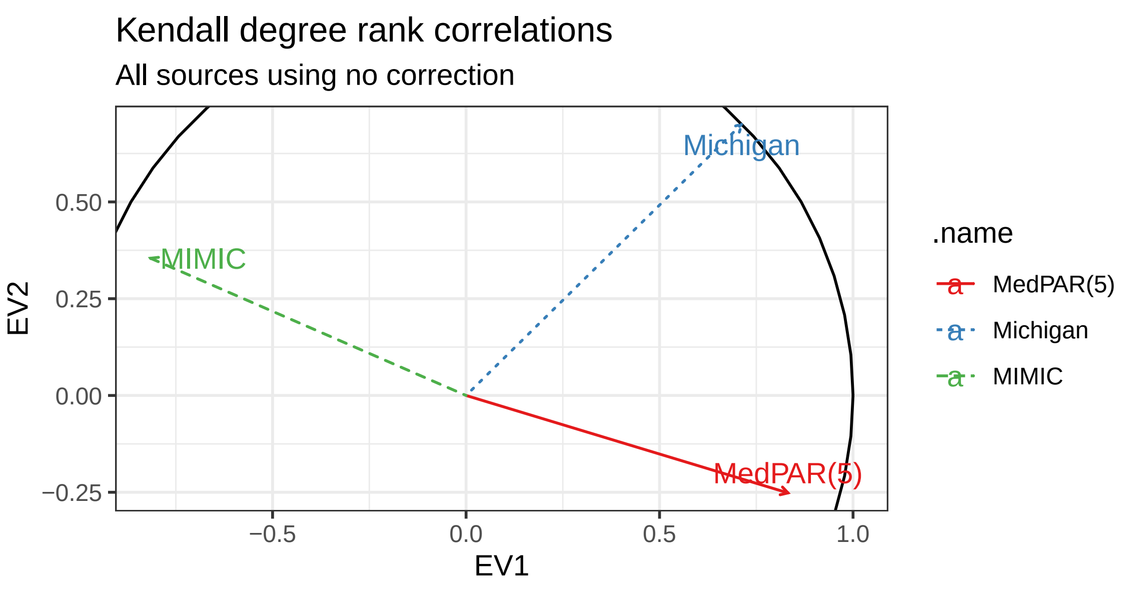

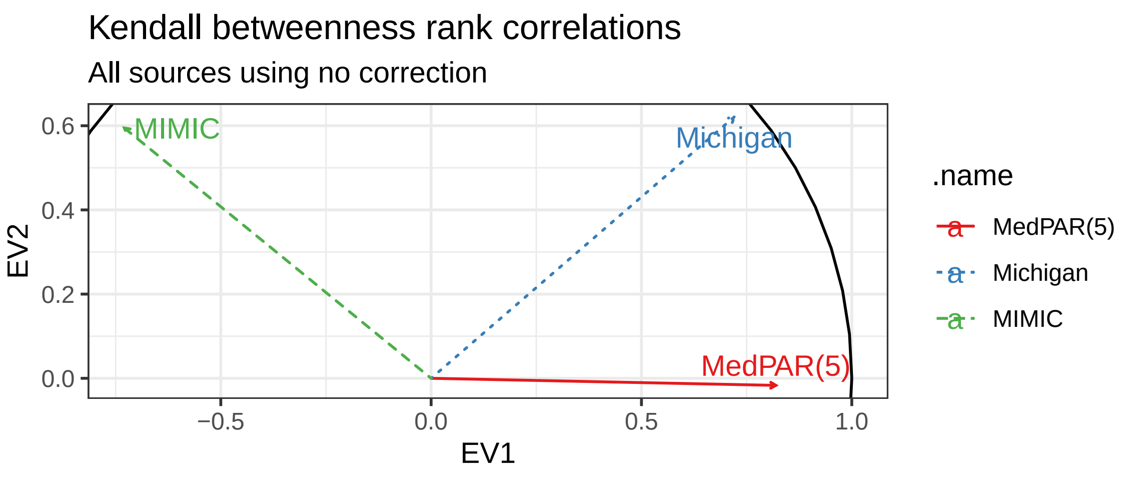

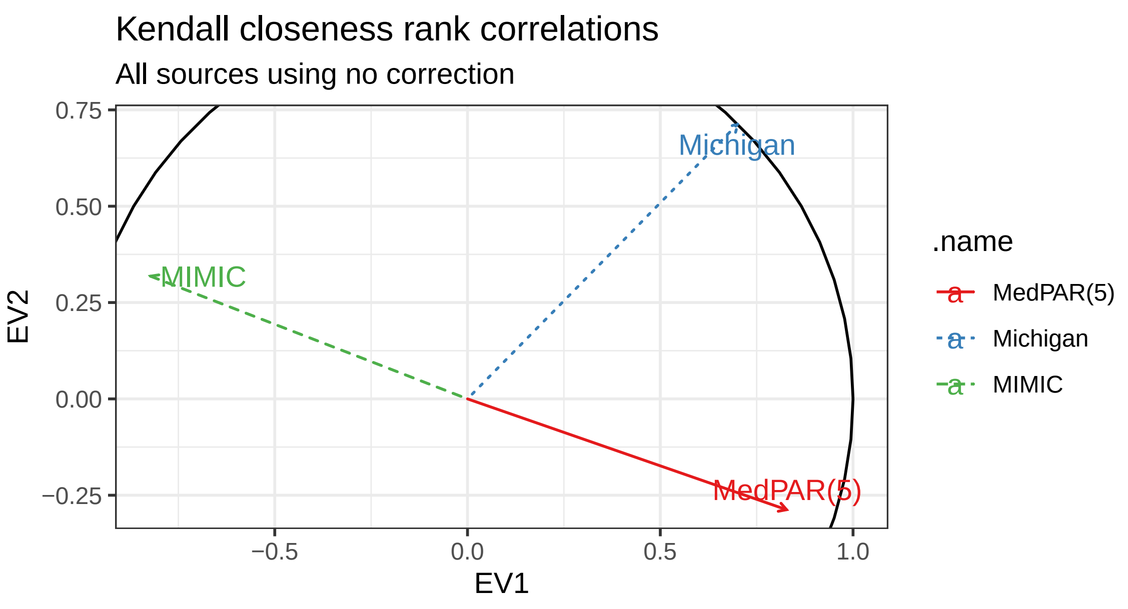

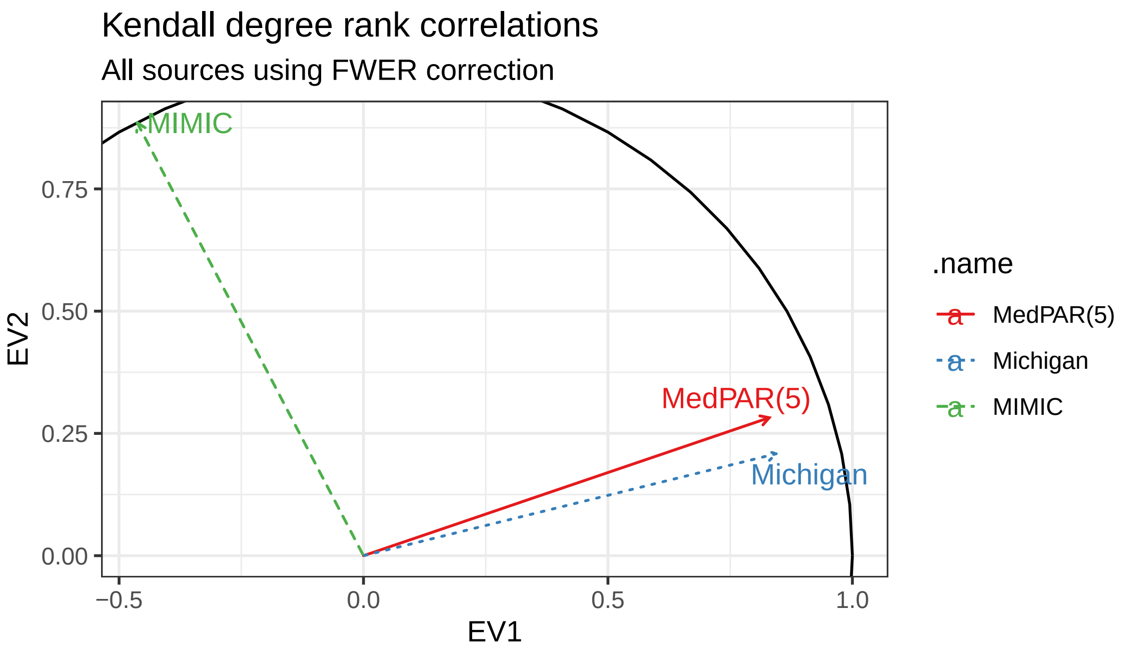

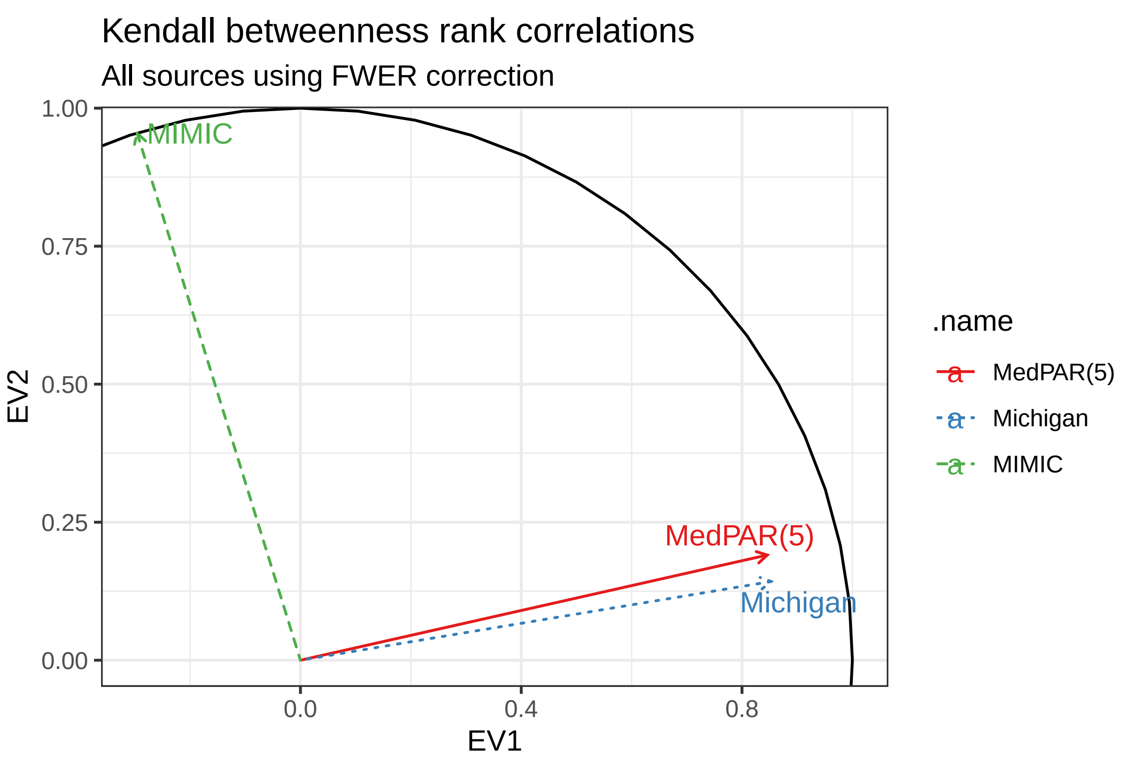

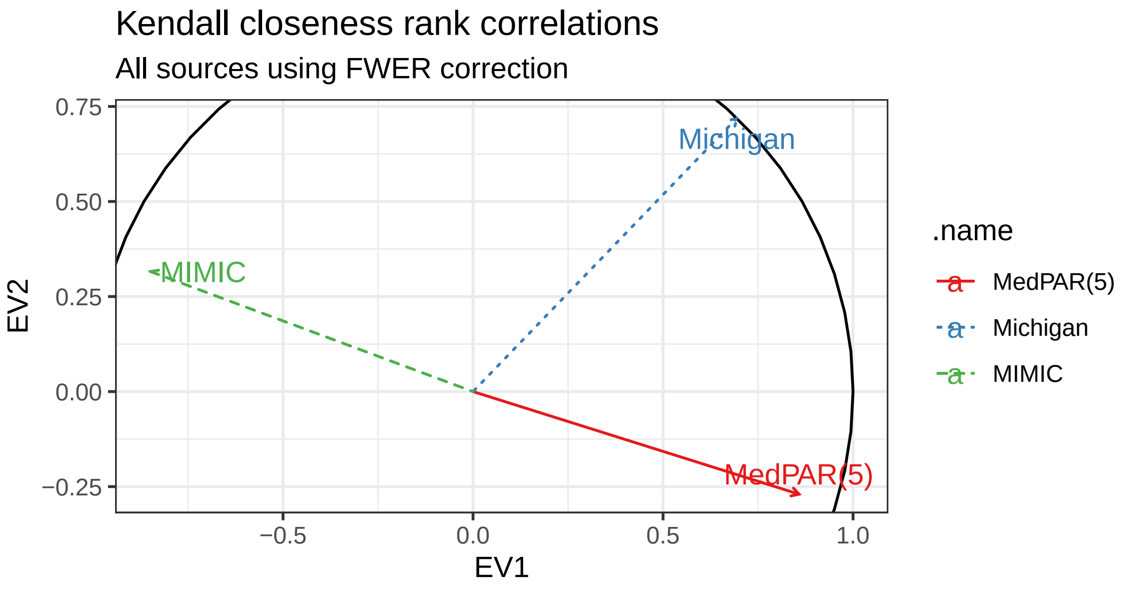

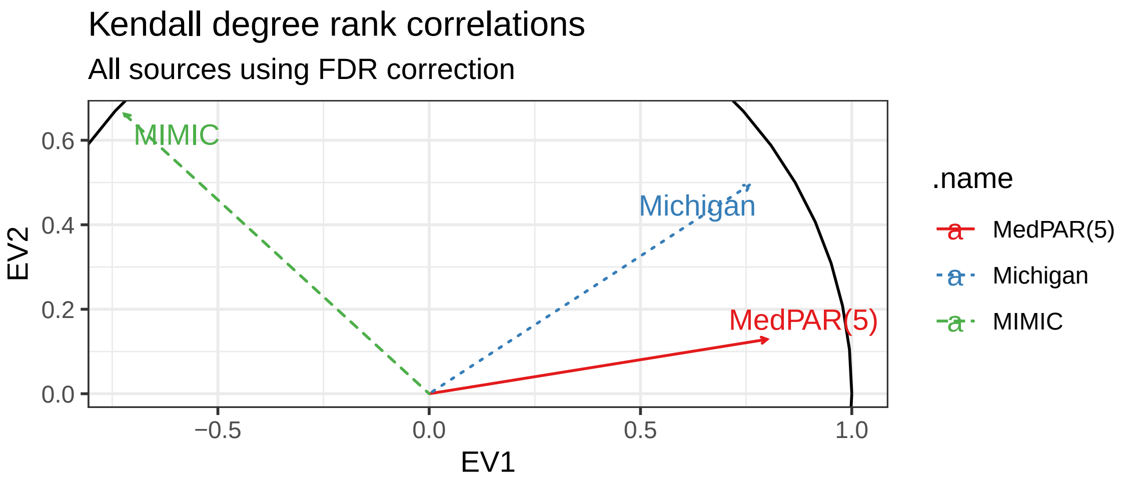

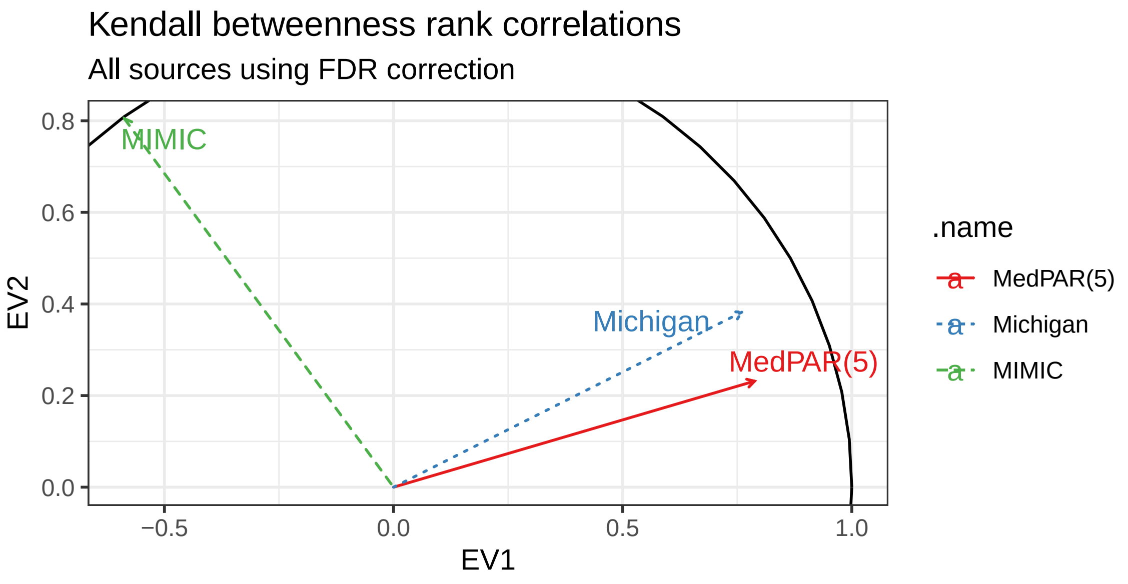

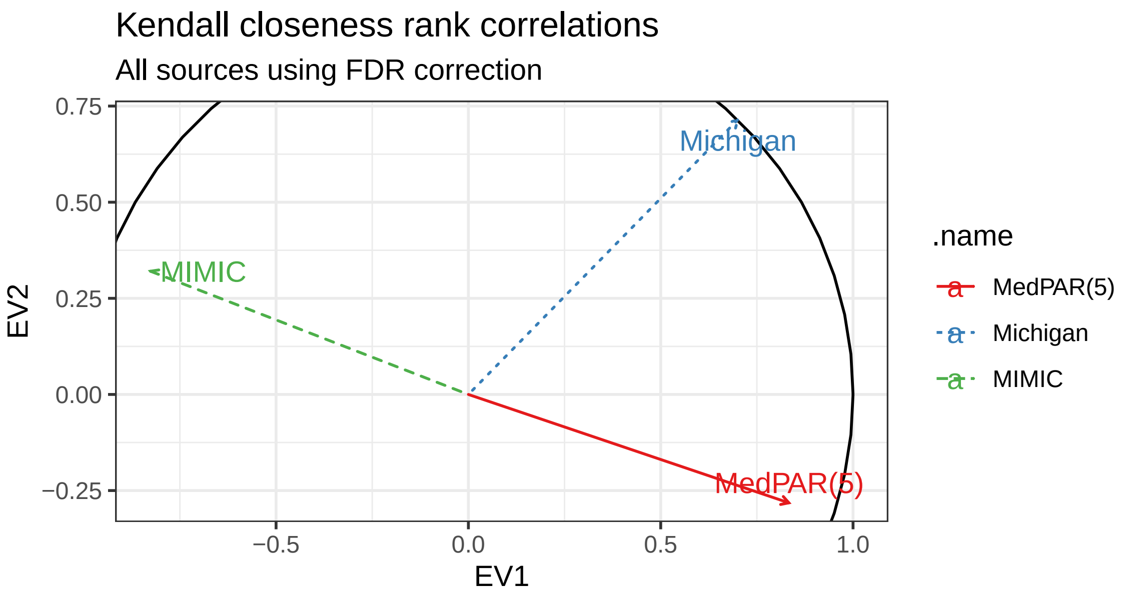

### Figure S19

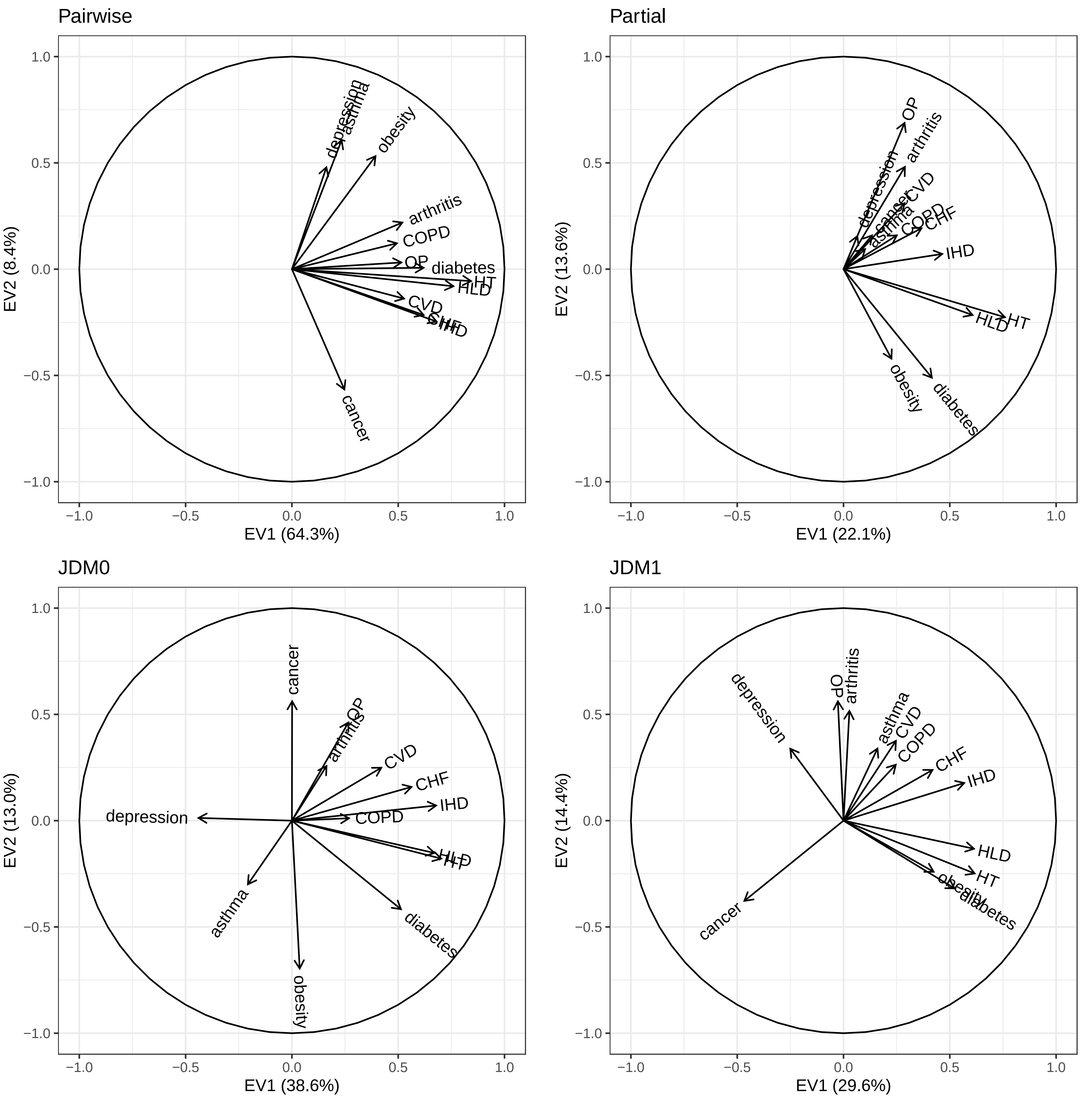

### Figure S20

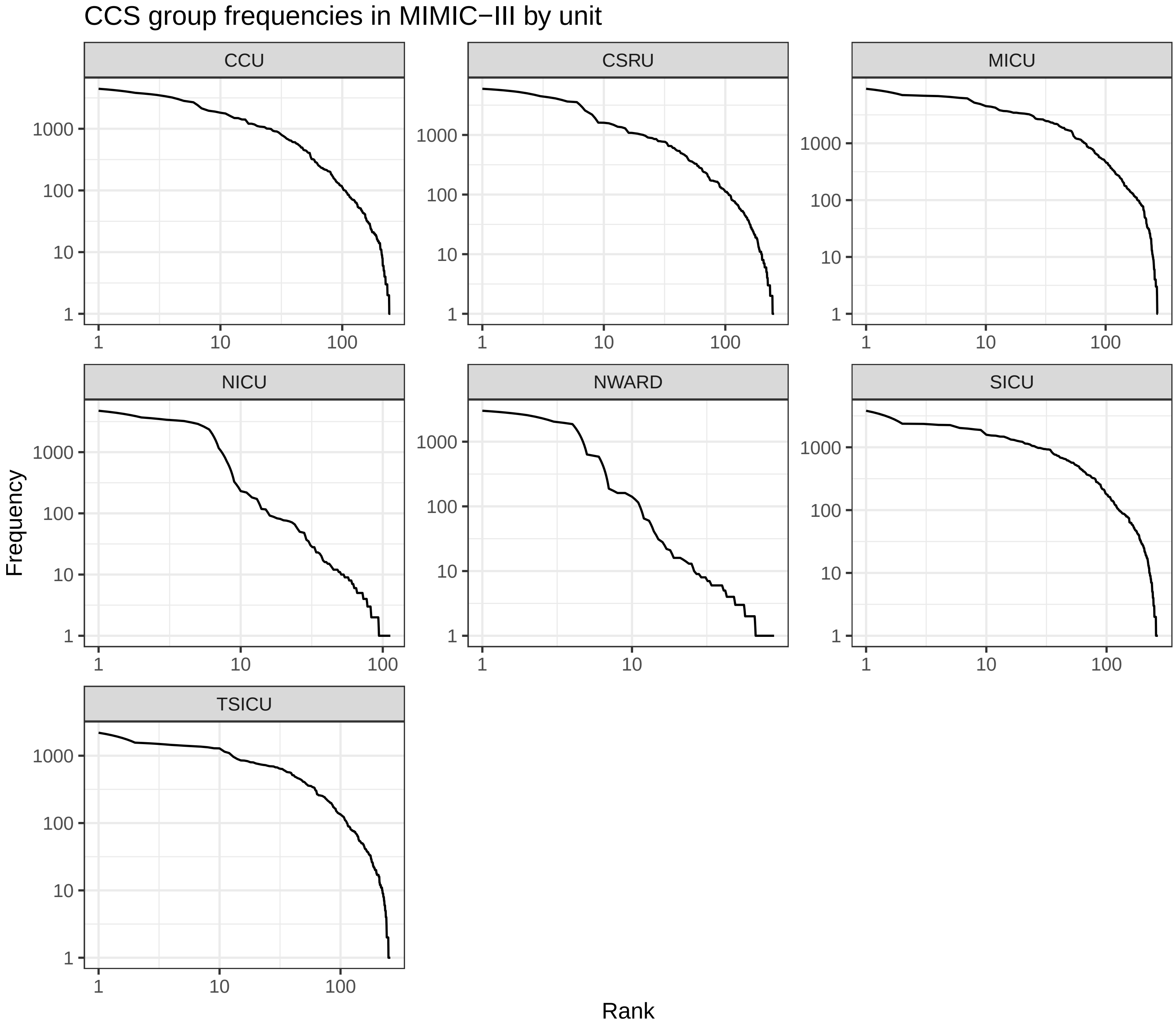

### Figure S23

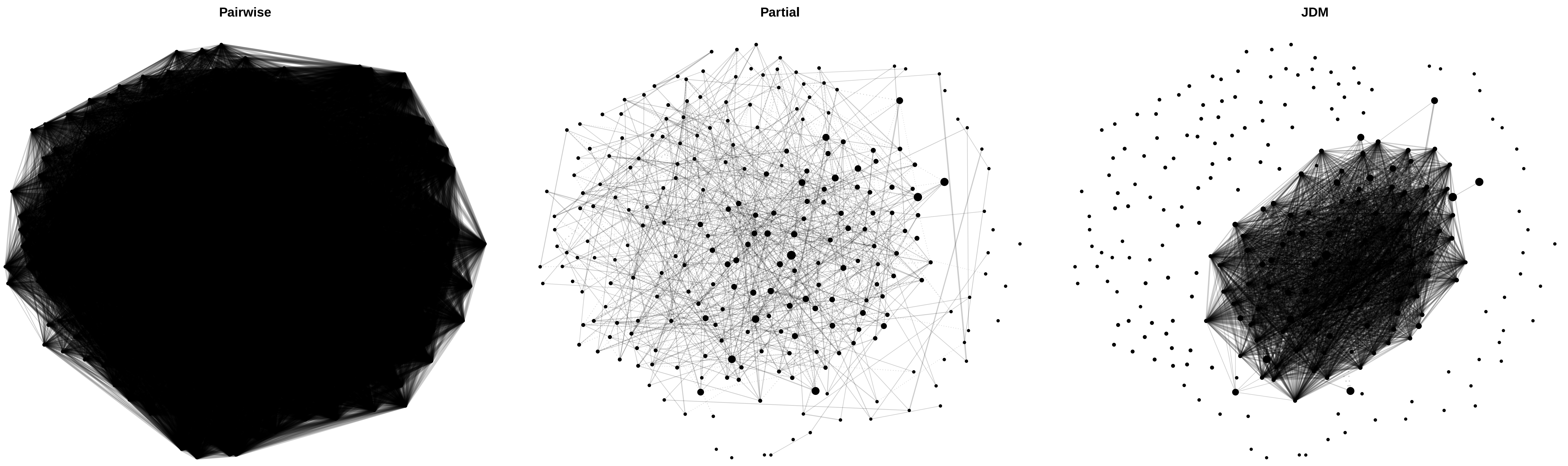

### Figure S24

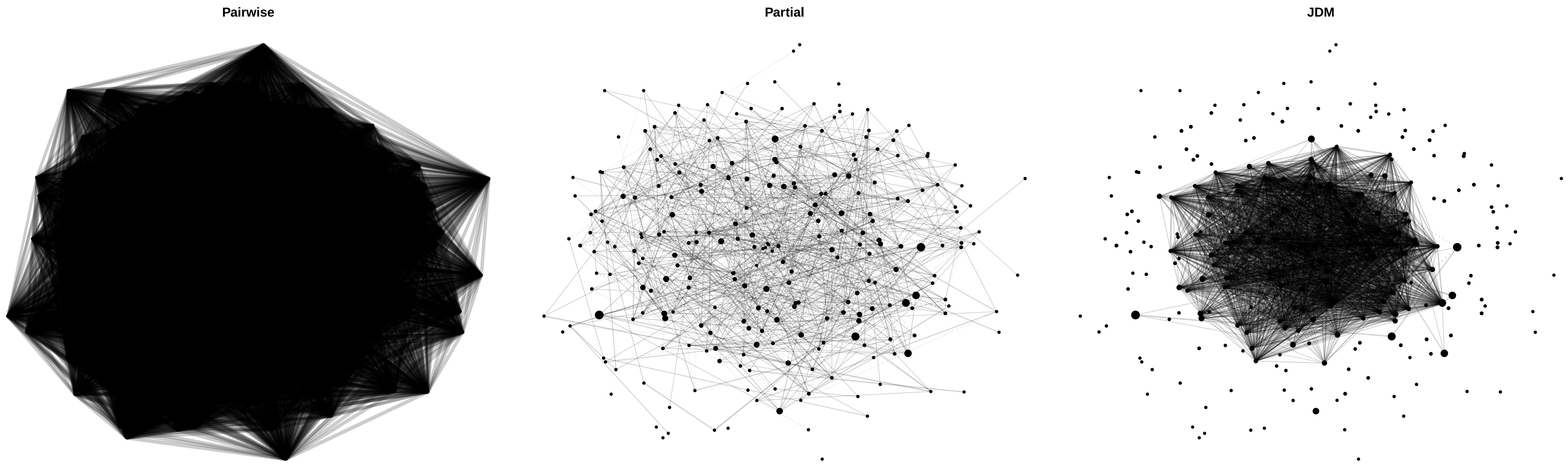

### Figure S25

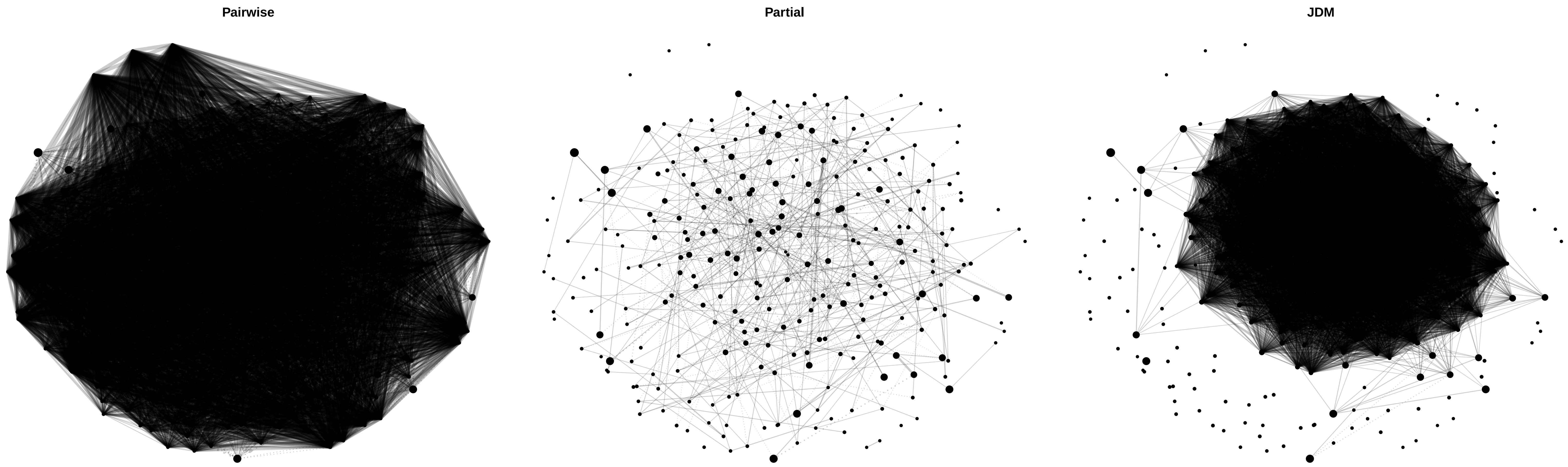

### Figure S26

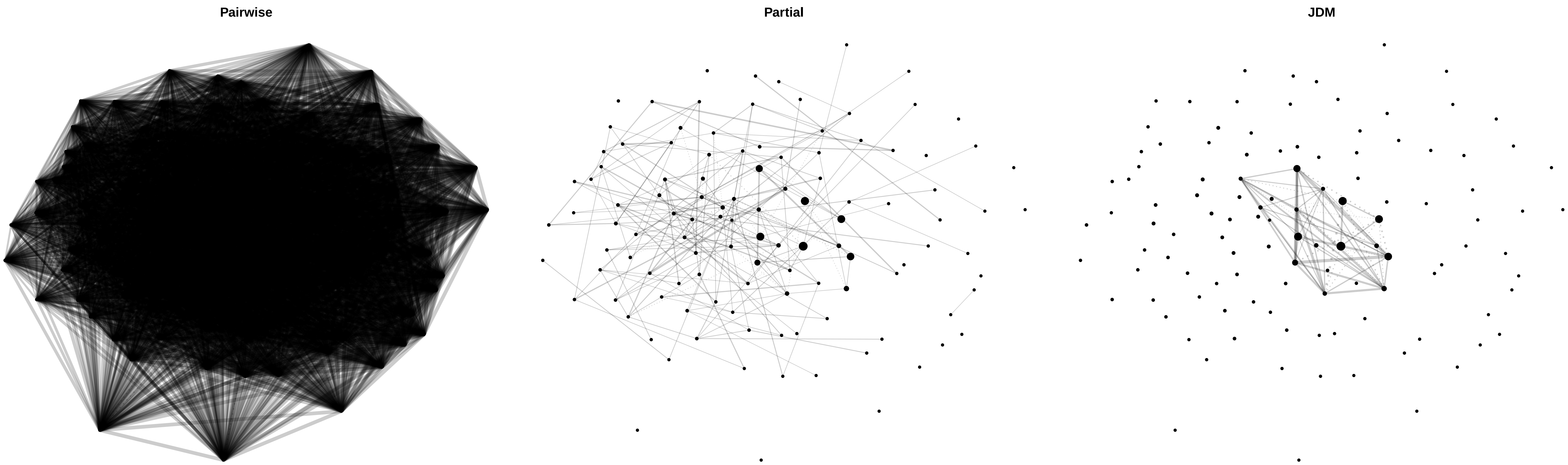

### Figure S27

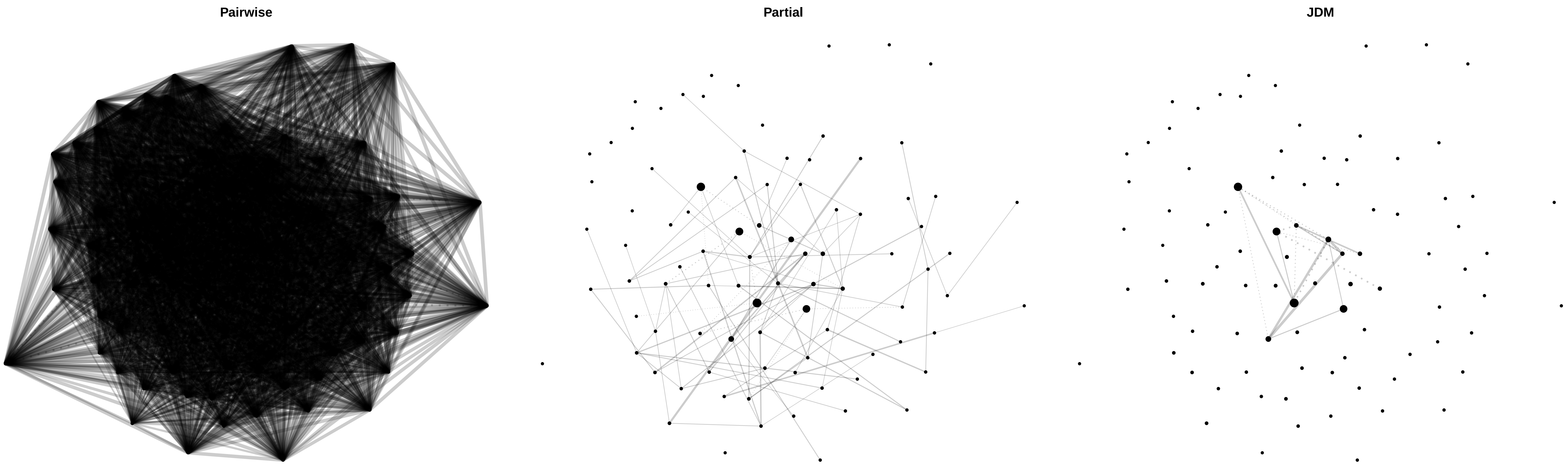

### Figure S28

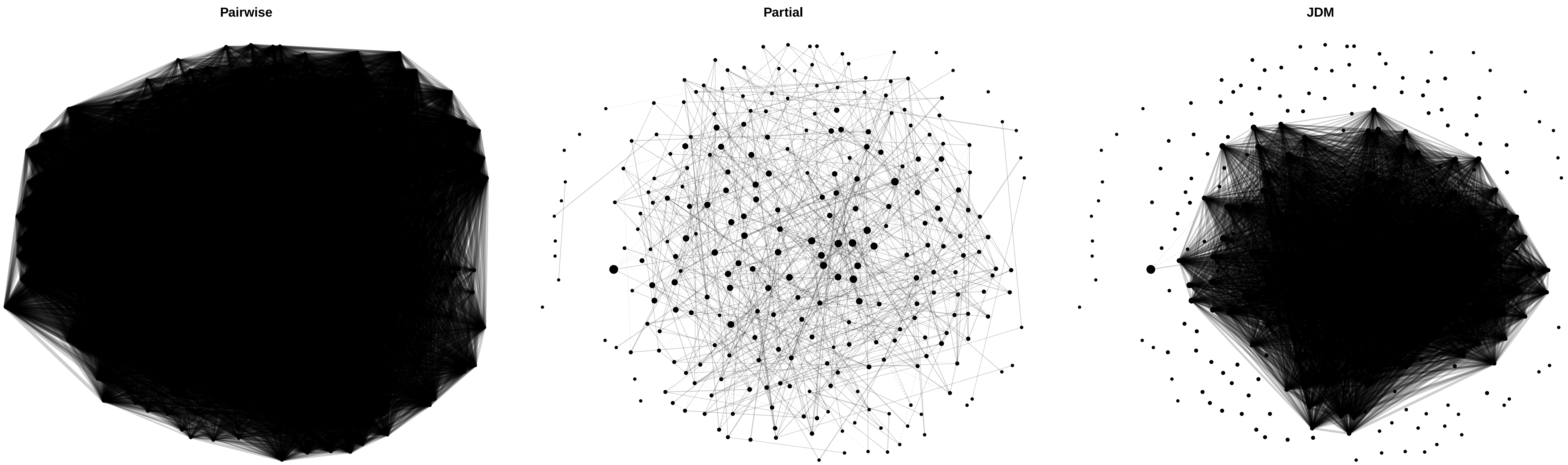

### Figure S29

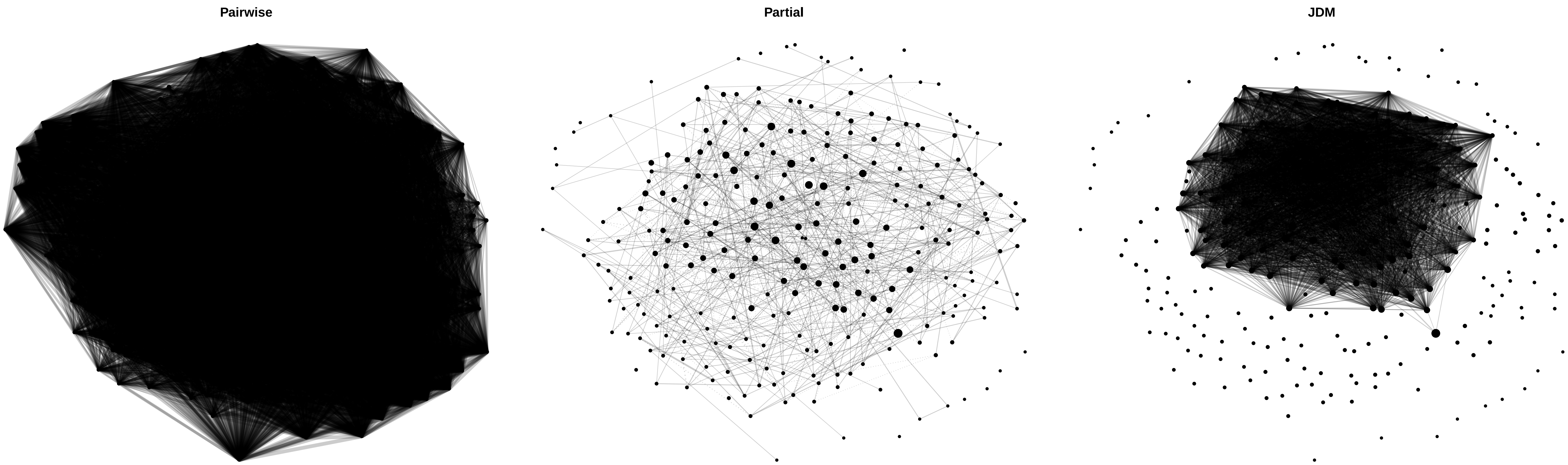

### Figure S30

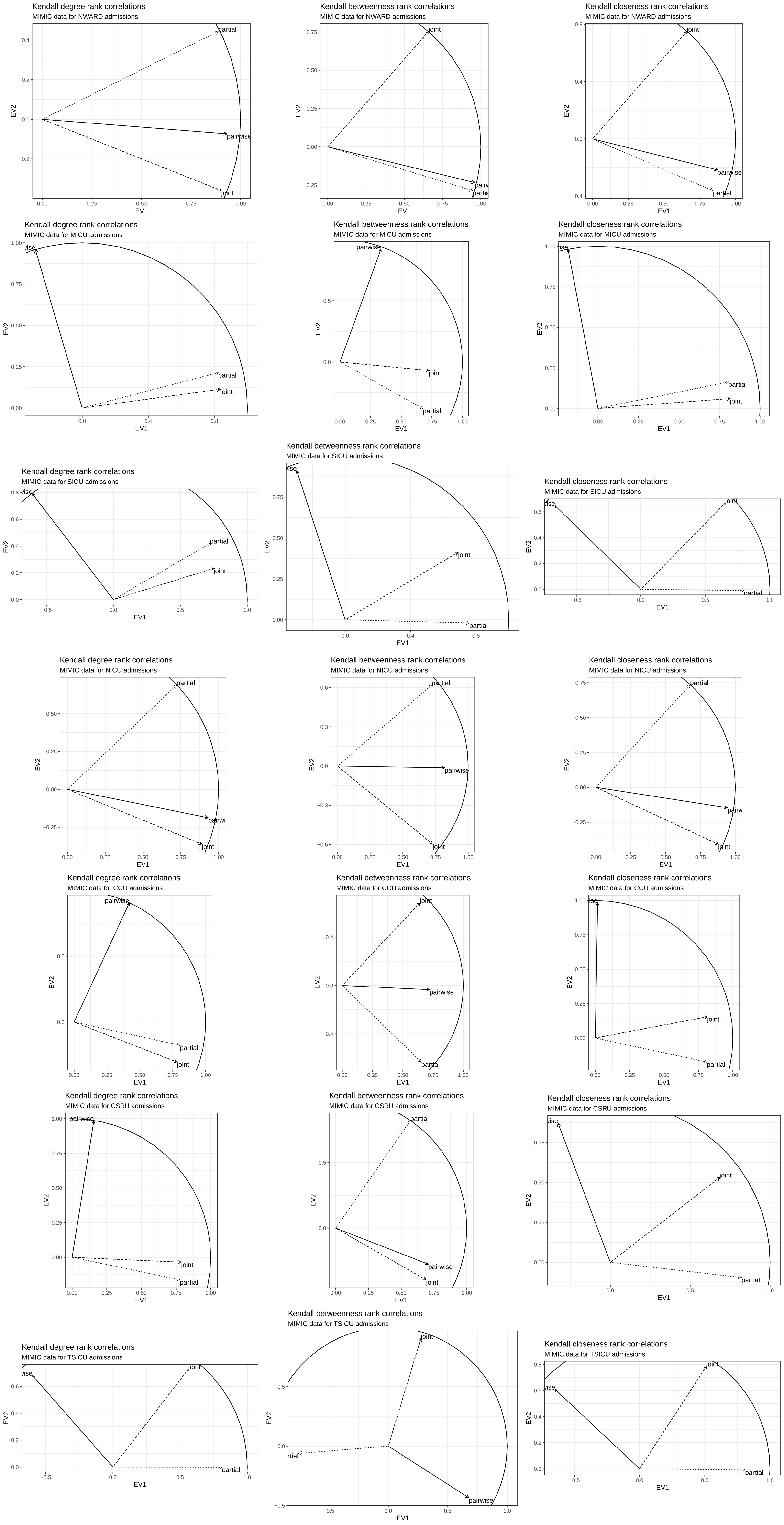
