## Supplementary material for "Sensitivity and robustness of comorbidity network analysis": Figure S13

Kendall degree rank correlations

All sources using no correction

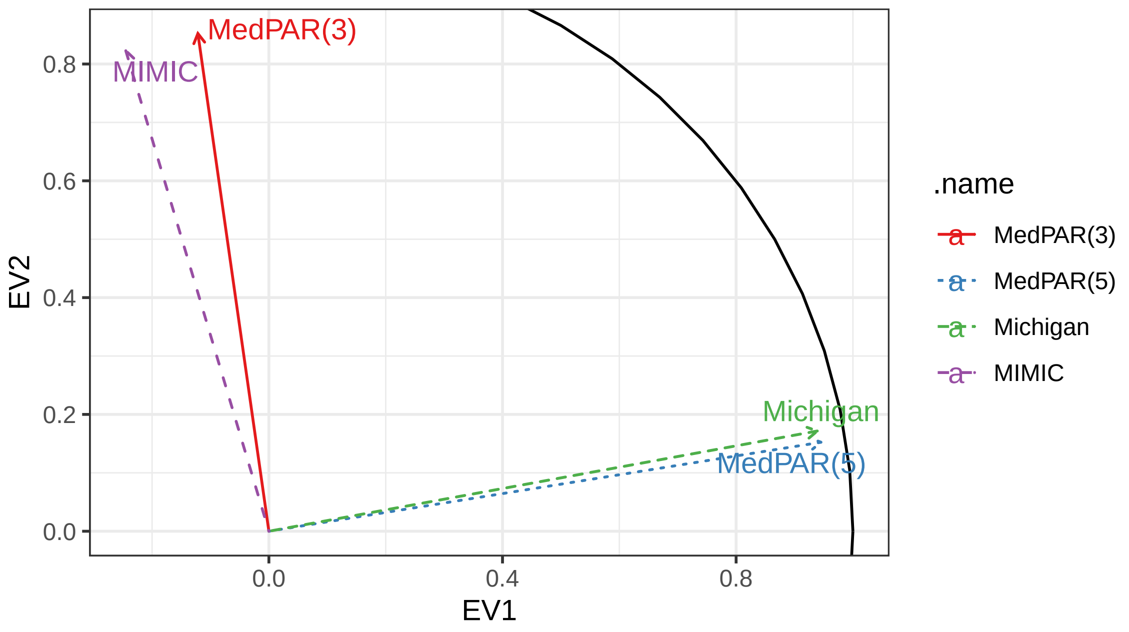


Kendall closeness rank correlations

All sources using no correction

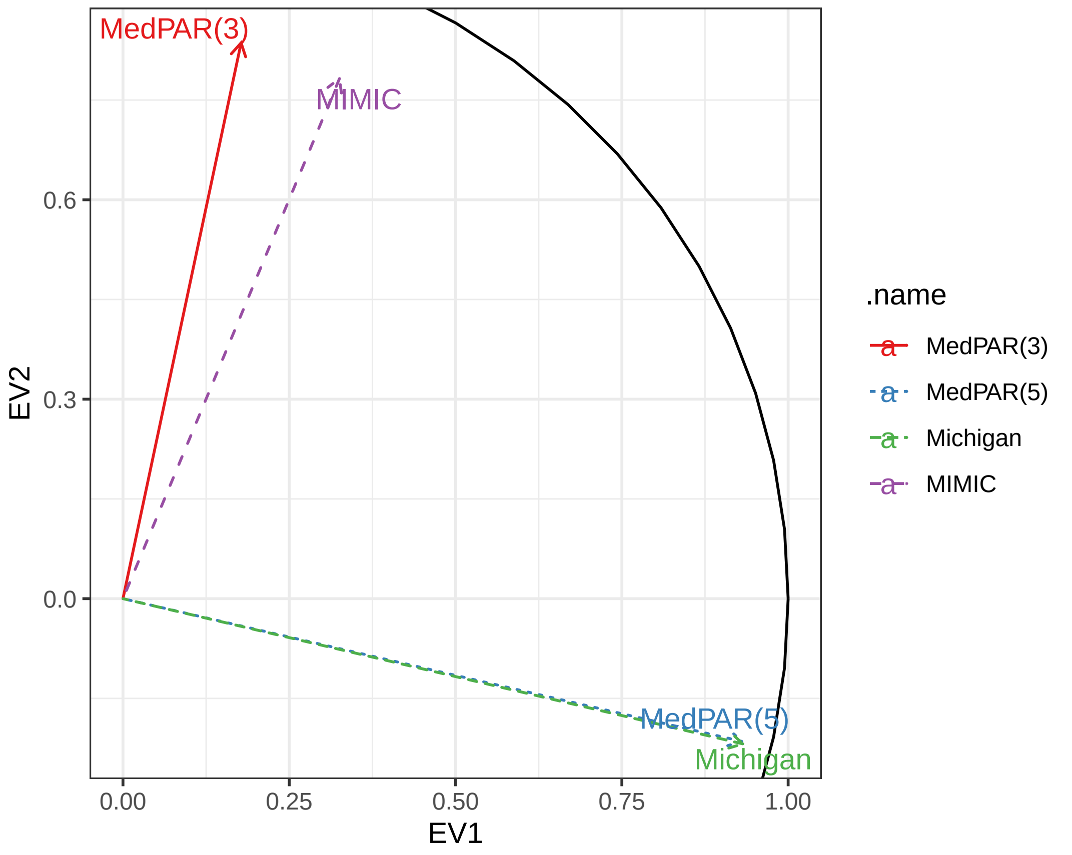


Kendall degree rank correlations

All sources using FWER correction

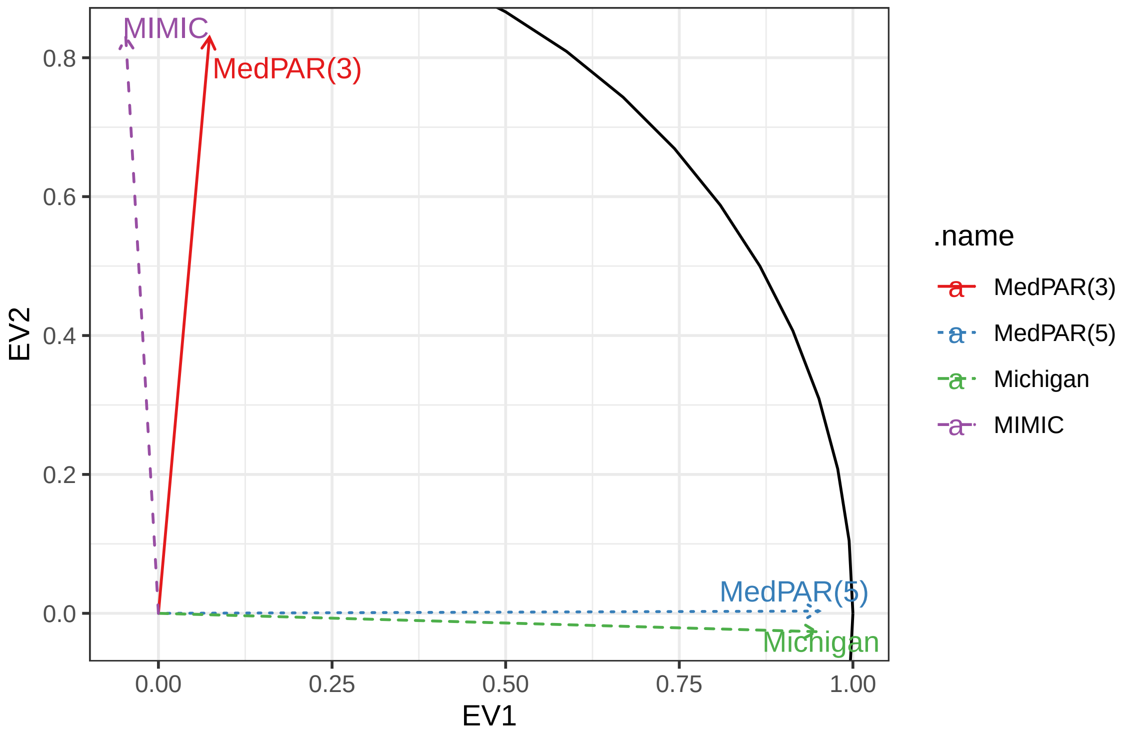


Kendall closeness rank correlations

All sources using FWER correction

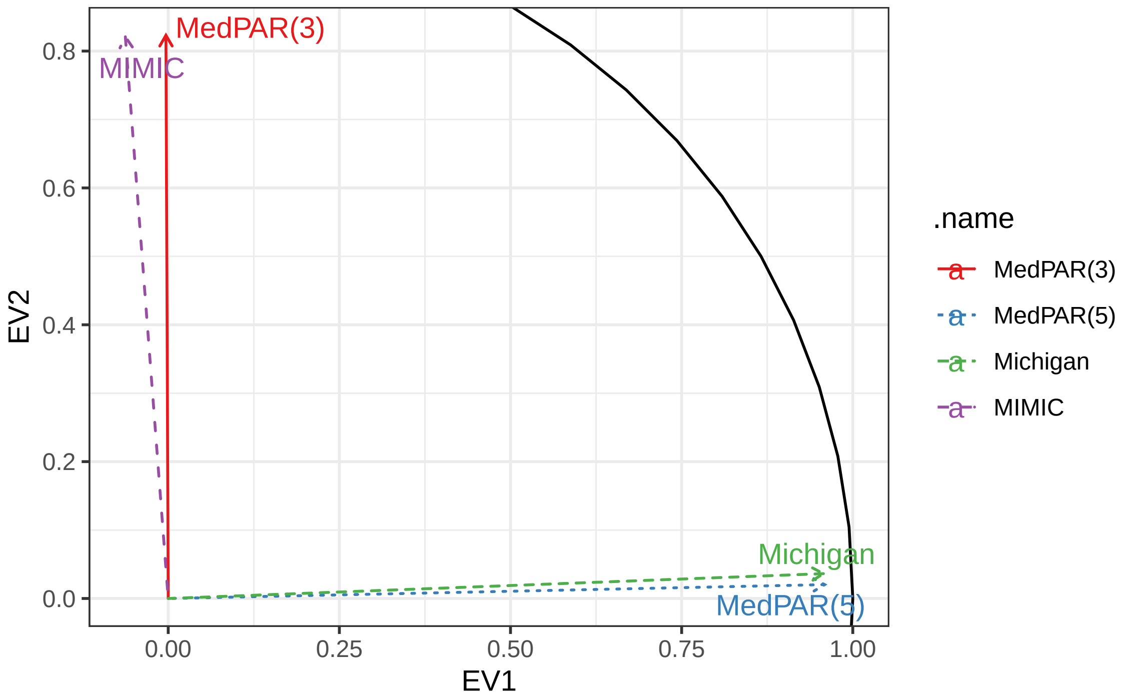


Kendall degree rank correlations

All sources using FDR correction

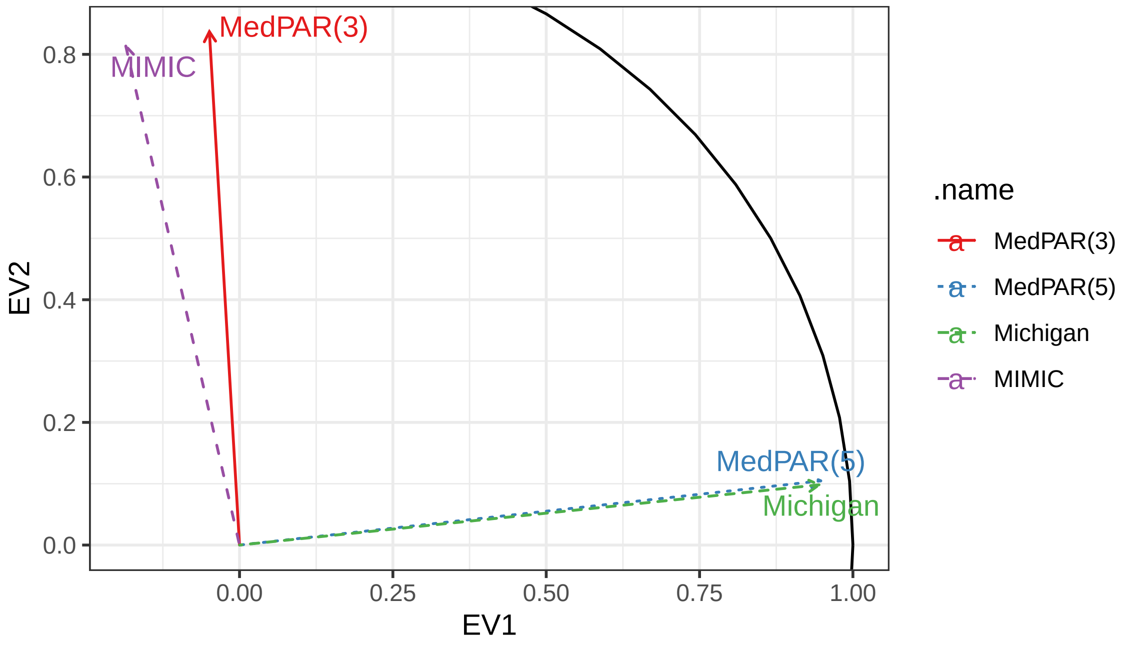


Kendall closeness rank correlations

All sources using FDR correction

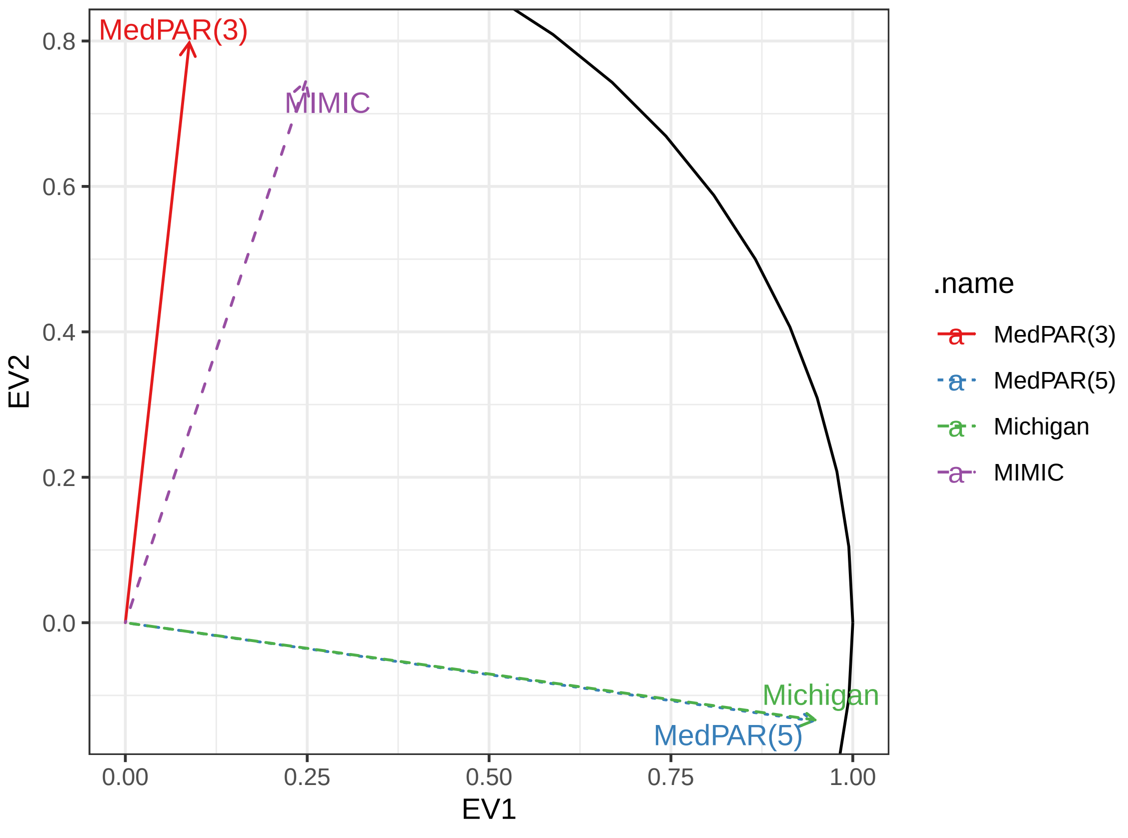
